# Learning evoked centrality dynamics in the schizophrenia brain: Entropy, heterogeneity and inflexibility of brain networks

**DOI:** 10.1101/2025.04.05.647398

**Authors:** Dhruval Bhatt, John Kopchick, Clifford C. Abel, Dalal Khatib, Patricia Thomas, Usha Rajan, Caroline Zajac–Benitez, Luay Haddad, Alireza Amirsadri, Jeffrey A Stanley, Vaibhav A Diwadkar

**Affiliations:** Department of Psychiatry and Behavioral Neurosciences, Wayne State University School of Medicine, Detroit, MI, 48201, USA

**Author notes:** Correspondence to: Vaibhav A. Diwadkar, PhD, Professor, Department of Psychiatry and Behavioral Neurosciences, Brain Imaging Research Division, Wayne State University School of Medicine, 3901 Chrysler Service Dr, Suite 5B, Tolan Park Medical Bldg, Detroit, MI, 48201, USA.

## Abstract

**Background:** Brain network dynamics are responsive to task induced fluctuations, but such responsivity may not hold in schizophrenia (SCZ). We introduce and implement Centrality Dynamics (CD), a method developed specifically to capture task-driven dynamic changes in graph theoretic measures of centrality. We applied CD to fMRI data in SCZ and Healthy Controls (HC) acquired during a learning paradigm.

**Methods:** fMRI (3T Siemens Verio) was acquired in 88 participants (49 SCZ). Time series were extracted from 246 functionally defined cerebral nodes. We applied a dynamic widowing technique to estimate 280 partially overlapping connectomes (30,135 region-pairs in each connectome). In each connectome we calculated every node’s Betweenness Centrality (BC) before building 246 unique time series (representing a node’s CD) from a node’s BC in successive connectomes. Next, in each group nodes were clustered based on similarities in CD.

**Results:** Clustering gave rise to fewer sub-networks in SCZ, and these were formed by nodes with greater functional heterogeneity. These sub-networks also showed greater ApEn (indicating greater stochasticity) but lower amplitude variability (suggesting less adaptability to task-induced dynamics). Higher ApEn was associated with worse clinical symptoms.

**Limitations:** Centrality Dynamics is a new method for network discovery in health and schizophrenia but will need further extension to other tasks and psychiatric conditions, before we achieve a fuller understanding of its promise.

**Conclusion:** The brain’s functional connectome is not static under task-driven conditions, and characterizing the dynamics of the connectome will provide new insight on the dysconnection syndrome that is schizophrenia. Centrality Dynamics provides novel characterization of task-induced changes in the brain’s connectome and shows that in the schizophrenia brain, learning-evoked sub-network dynamics were less responsive to learning evoked changes and showed greater stochasticity.

## Introduction

Schizophrenia is an aberrant dynamic brain process (1) that cannot be explained by lesion-based models (2). Unravelling schizophrenia may hinge on our understanding of disordered brain network dynamics, given that normative brain function emerges from dynamically evolving network interactions (3–5). These dynamics in fMRI data can be quantified in short time series “bursts” using continuous windows that span across the acquisition (6, 7). This approach for dynamic functional connectivity (FC) has been used to study a) state transitions in resting state fMRI signals (rsfMRI)(8, 9), b) disordered state transitions in schizophrenia (10), and c) increased stochasticity in brain states in the illness (11). However, at least two lacunae exist: 1) Dynamic FC has only infrequently been used with task-based fMRI despite the knowledge that they are significantly modulated by task-induced changes (12); 2) Changes in brain states can be captured in individual time series, but they may be more compellingly expressed in the dynamics of the connectome (13). The first lacuna can be addressed for example, by using dynamic FC to quantify contextually induced changes in schizophrenia-relevant tasks such as dynamic associative learning (14–17). Here, we address the second lacuna with a novel application of graph theoretic centrality techniques (18). We motivate the concept of Centrality Dynamics (CD) and then study it in fMRI data specifically acquired in a large group of SCZ patients and typical controls while participants engaged in dynamic learning (19).

### Networks, Dynamics, Schizophrenia

Brain networks are isomorphic to undirected (or directed) graphs (18, 20–23). Here, the functional importance of nodes can be efficiently estimated using node centrality measures. Thus, a measure like Betweenness Centrality (BC) efficiently summarizes information across the cumulative set of graph edges (20) and is particularly useful because a node’s BC quantifies the extent to which it lies on any number of shortest paths between other nodes in the network. BC is therefore, a heuristic estimate of a node’s integrative importance or “hubness” (24) and is a widely used in summarizing node importance in stationary connectomes (21). However, connectomes in natural biological systems possess complex dynamics (and these dynamics often reflect systemic interactions with the external environment)(25). Furthermore, these dynamics may be reflected in how centrality measures like BC change over the course of a task (26). In this view, task-evoked connectomic changes will be reflected in dynamic changes of the BC of nodes, precisely because a node’s integrative role may change over the course of the task.

Here, we drove brain network dynamics using an established learning task(19, 27). From the co-acquired time series data, we summarized the connectomic properties of the cerebral network in each of a sequence of overlapping moving windows (designed to cover the entirely of the experiment). From each node’s derived BC in each window (18) we then built a time series which represents the centrality dynamics for that node (28). Next, nodes were clustered based on the similarity of their centrality dynamics (using agglomerative hierarchical clustering) (29, 30). This clustering revealed distinct sub-networks in each of SCZ and HC. In schizophrenia, the discovered sub-networks showed a) less functional diversity of their constituent nodes; b) greater entropy of their centrality dynamics (suggesting greater stochasticity), where this entropy was generally associated with greater clinical deficits and c) greater inflexibility. Coupled with recent evidence for disordered dynamics of the salience network in schizophrenia (31) and during cognitive processing (based on EEG signals)(32), our observations provide compelling evidence of the dynamics of dysconnection in schizophrenia.

## Materials and Methods

### Participants

Eighty-eight subjects gave informed consent to participate in the study (approved by the Institutional Review Board at the WSU School of Medicine) and received remuneration for their involvement. The cohort included 49 stabilized (with atypical antipsychotics) schizophrenia patients and 39 healthy controls. Patients were identified through their treating physicians (L.H., A.A.), and diagnosis was confirmed by a research psychologist (U.R.) using DSM-5 criteria (33). All patients were maintained on a regime of atypical antipsychotics (risperidone, olanzapine, or aripiprazole). The PANSS was used to assess clinical symptom severity ratings (34) and the Wechsler Abbreviated Scale of Intelligence to measure general intelligence (35). Duration of illness was estimated using the date of diagnosis for schizophrenia, the estimated date of onset of psychotic symptoms (hallucinations, delusions, or disorganization of thinking; bizarre or catatonic behavior), and medical records, reports by family members or significant others, and the Structured Clinical Interview for DSM Disorders interview. All healthy controls were free of past or present Axis-I psychopathology. All participants were screened before entering the study to exclude any significant past/current medical and/or neurological illness (e.g., hypertension, thyroid disease, diabetes, asthma requiring prophylaxis, seizures, or significant head injury with loss of consciousness). The two groups were similar in age or gender distribution. Table 1 provides demographic and clinical information.

**Table 1.** Demographic and clinical information for HC and SCZ participants. Patients were stabilized under a regime of antipsychotics. HC were, free of psychiatric symptoms (or medications) (see Methods).

|  | Healthy Controls | Schizophrenia Patients |
| --- | --- | --- |
| N | 39 | 49 |
| Age (Years) | 27.59 ± 6.71 | 31.47 ± 8.61 |
| Gender | 10 F / 29 M | 9 F / 40 M |
| Handedness | 30 Right | 37 Right |
| FSIQ-4 Score | 101.85 ± 9.46 | 88.29 ± 7.94 |
| PANSS Positive Score | | 13.76 ± 3.44 |
| PANSS Negative Score | | 14.24 ± 3.92 |
| PANSS General Score | | 26.02 ± 5.81 |
| Global Assessment Scale | | 65.86 ± 4.41 |
| Social Adaptation Self-evaluation Scale | | 27.53 ± 6.62 |

### MRI acquisition and pre-processing

Data (3 T Siemens Verio scanner, 32-channel volume head coil) were acquired using a multiband gradient EPI sequence (TR = 3 s, TE = 24.6 s, multiband factor = 3, FOV = 192 × 192 mm^2^, matrix = 96 × 96, 64 axial slices, resolution = 2 mm^3^). T_1_-weighted MRI images were collected for normalization and co-registration with the EPI scan (3D magnetization-prepared rapid gradient-echo sequence, TR = 2,150 ms, TE = 3.5 ms, TI = 1,100 ms, flip angle = 8°, FOV = 256 × 256 × 160 mm^3^, 160 axial slices, resolution = 1 mm^3^).

### Image Processing

MRI data were processed using standard temporal (slice-time correction) and spatial preprocessing methods in SPM 12. The EPI images were aligned to the AC-PC line, realigned to a reference image in the sequence to correct for head movement, and co-registered to the high-resolution T1 anatomical image. The EPI images were normalized to stereotactic space by applying the deformations from normalizing the high-resolution T1 image. A low-pass filter (128 s) was used to remove low frequency components associated with respiratory and cardiac rhythms. At the first level, boxcar stimulus functions were convolved with a canonical hemodynamic response function modeled epochs as regressors of interest with the six motion parameters (3 for translation and 3 for rotation) from the co-registration modeled as covariates of no interest. The images were resliced (2 mm3) and with a spatial filter applied (8 mm FWHM). Images with more than 4 mm of movement (<1% of all images) were excluded from analyses.

### Dynamic Learning Task

Task dynamics were evoked using an established paired-associate learning paradigm (19, 21, 36). Participants learned associations between nine equi-familiar objects (37) and their designed unique grid locations to which they had been assigned. The eight task iterations cycled between epochs (27 s) for Encoding, Post-Encoding Consolidation, Retrieval, and Post-Retrieval Consolidation (Supplementary Figure 1 provides behavioral data from participants in both groups). The entire paradigm lasted 864 s (288 images).

### Estimating Centrality Dynamics

Figure 1 depicts our quantitative pipeline. First, from the pre- processed fMRI data, in each of the 88 study participants, we extracted averaged time series from 246 regions in the Brainnetome cerebral atlas (28) (chosen based on its use of multi-modal data for functional parcellation). Next, bivariate zero-lag correlations (undirected functional connectivity) (38) were estimated between all 30,135 pairs of regions (_246_C_2_) in each of 280 contiguous and partially overlapping windows. Window widths (27 s or 9 images) coincided with condition durations. We used a maximal overlap between successive windows to ensure maximal coverage across the acquisition (24 s/8 images).

**Figure 1.**
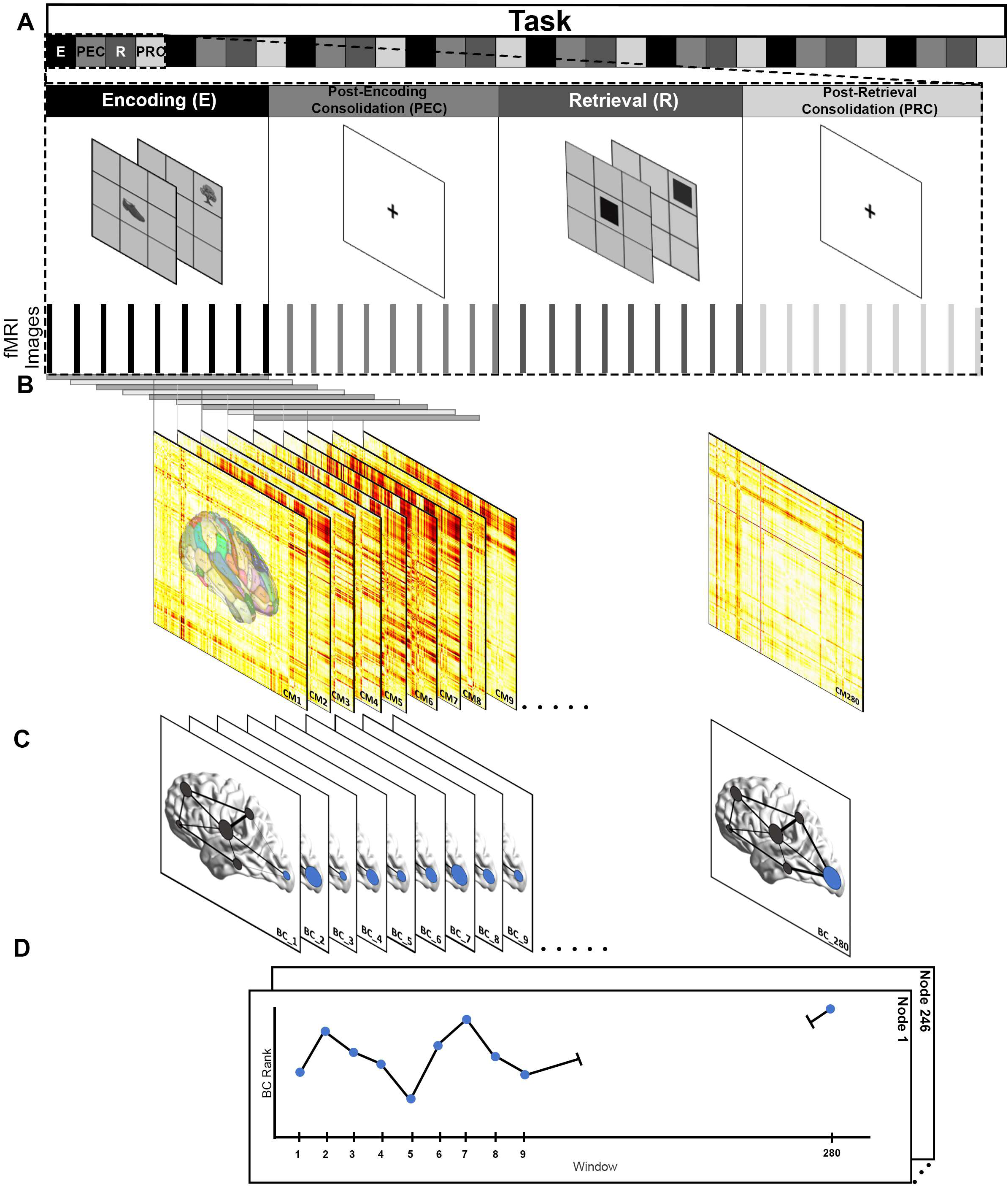
Overview of Analytical Pipeline. The Figure provides an overview of the arc of the analyses, beginning with a depiction of the task (a) and ending in a schematic depiction of the centrality dynamics for a node (d). (a) The learning paradigm required participants to learn nine object-location associations over eight iterations of the paradigm (a single iteration is depicted). Each iteration consisted of four successive epochs for Encoding, Post-Encoding Consolidation, Retrieval, and Post-Retrieval Consolidation (9 fMRI images/27 s). Data for the task were acquired over 288 images (864 s). (b) A moving window approach was adopted as the first step in computing centrality dynamics. Each window width corresponded to the width of each task epoch (9 images), and using a sliding approach each successive window was advanced by one image (resulting in 280 windows over which centrality dynamics were computed). Within each window, the full zero-lag connectivity matrix (CM) was estimated in the 246-region connectome (280 successive CMs, CM_1_ – CM_280_). (c) The next step in recovering centrality dynamics across the 280 windows was to estimate the Betweenness Centrality of each of the 246 nodes in each CM. From the estimated BC values, the integrative importance of nodes was ranked (see Methods). The schematic network depicts nodes where the node size is a representation of its ordinal rank in each CM, and changes in its size over successive CMs are a visual depiction of the node’s centrality dynamics. (d) Next, the centrality values were used to form a 280-point time series (henceforth t_BC_) for each node (a partial schematic is shown). Finally, within each group the 246 t_BC_ were averaged across participants, and the averaged t_BC_ was submitted for agglomerative clustering. The goal of clustering was to identify groups of nodes with homogenous centrality dynamics.

Coefficients (Pearson’s r) in the resultant symmetric connectivity matrix (CM) were normalized (Fisher’s Z) (39). Each participant contributed 280 CMs (CM_1_ – CM_280_) for subsequent analyses.

### Estimating and Ranking Betweenness Centrality (BC)

For every pair of regions in any connected graph (30,135 pairs in our case), there exists at least one shortest path between the regions such that the sum of the weights of the edges is minimized. Any region’s BC is the number of these shortest paths that pass through it (24). BC, captures a node’s integrative importance in any defined network where a higher BC suggests greater influence over the network (21). We estimated the BC for each of the 246 nodes for each CM_i_ using the ANTs library in R (see below)(40).

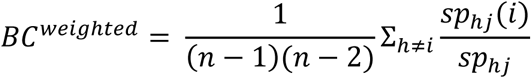

In each CM_i_ we ranked nodes (Spearman’s Rank method)(41) based on their BC value (246, Highest BC – 1, Lowest BC). As a result, for each of the 246 nodes we were able to form a 280-point (1 point for each CM_i_) time series (henceforth t_BC_) where the value at each time point represents a node’s rank within that CM_i_.

Notably, the shape of each t_BC_ captures the node’s Centrality Dynamics. Next, the t_BC_ were clustered (separately in HC and SCZ groups) using Agglomerative Hierarchical Clustering.

### Agglomerative Hierarchical Clustering (AHC)

Before clustering, the mean t_BC_’s in each of the HC and SCZ groups were computed and following this we computed the full cross-correlation matrix (30,135 unique correlations). Every cell in each matrix captures similarities in centrality dynamics between pairs of nodes. The resulting correlation coefficients served as inputs for AHC. We used AHC (Ward method) because a) it is robust in generating solutions independent of cluster size, b) it is effective in clustering high-dimensional data (29, 30), and c) it uses a simple similarity metric (coefficients) as a basis for clustering. In each clustering solution, we determined the optimal number of clusters through the convergence of a) the Within-Cluster Sum of Squares (elbow) plot and b) the threshold versus number of clusters plot (ensuring stable cluster sizes). Clusters were numbered in descending order of their intra-cluster correlation (HC_C1_ – HC_Cn_; SCZ_C1_ – SCZ_Cm_). We computed the Approximate Entropy (ApEn) and the Variance of each cluster (see below) over the average of all t_BC_ in the cluster (this average is referred to as t_Cluster_).

### Approximate Entropy

ApEn is an information theoretic index of the regularity or unpredictability of a time series’ fluctuation *where a higher ApEn indicates greater unpredictability* (i.e., lower regularity). ApEn was calculated in Matlab using the following formula.

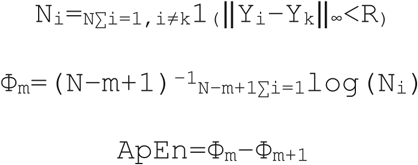

### Variance

We also calculated a complementary (to ApEn) metric, specifically the variance of each t_Cluster,_ where higher variability in a cluster’s centrality dynamics implies greater sub-network flexibility in responding to task dynamics (42, 43).

## Results

We organize our results as follows: 1) We first depict the unique AHC solutions reached from the t_BC_ in each of the HC and SCZ groups (Fig. 2), before (Fig. 3) showing that these sub- networks have differentiable bases. 2) We then examined the average centrality dynamics (t_Cluster_) of each sub-network (Fig. 4). In Fig. 5, we quantify the heterogeneity in periodicity and variability of clusters in Fig. 4, where ApEn and variance are depicted for each t_Cluster_, and wherein we also depict the relationship between ApEn and variance (across clusters). 3) We next explore how changes in the regional composition of sub- networks are related to ApEn (Fig. 6). 4) Finally (Fig. 7) we show that in schizophrenia, in two of the three sub-networks, key clinical variables are significantly associated with the ApEn.

**Figure 2.**
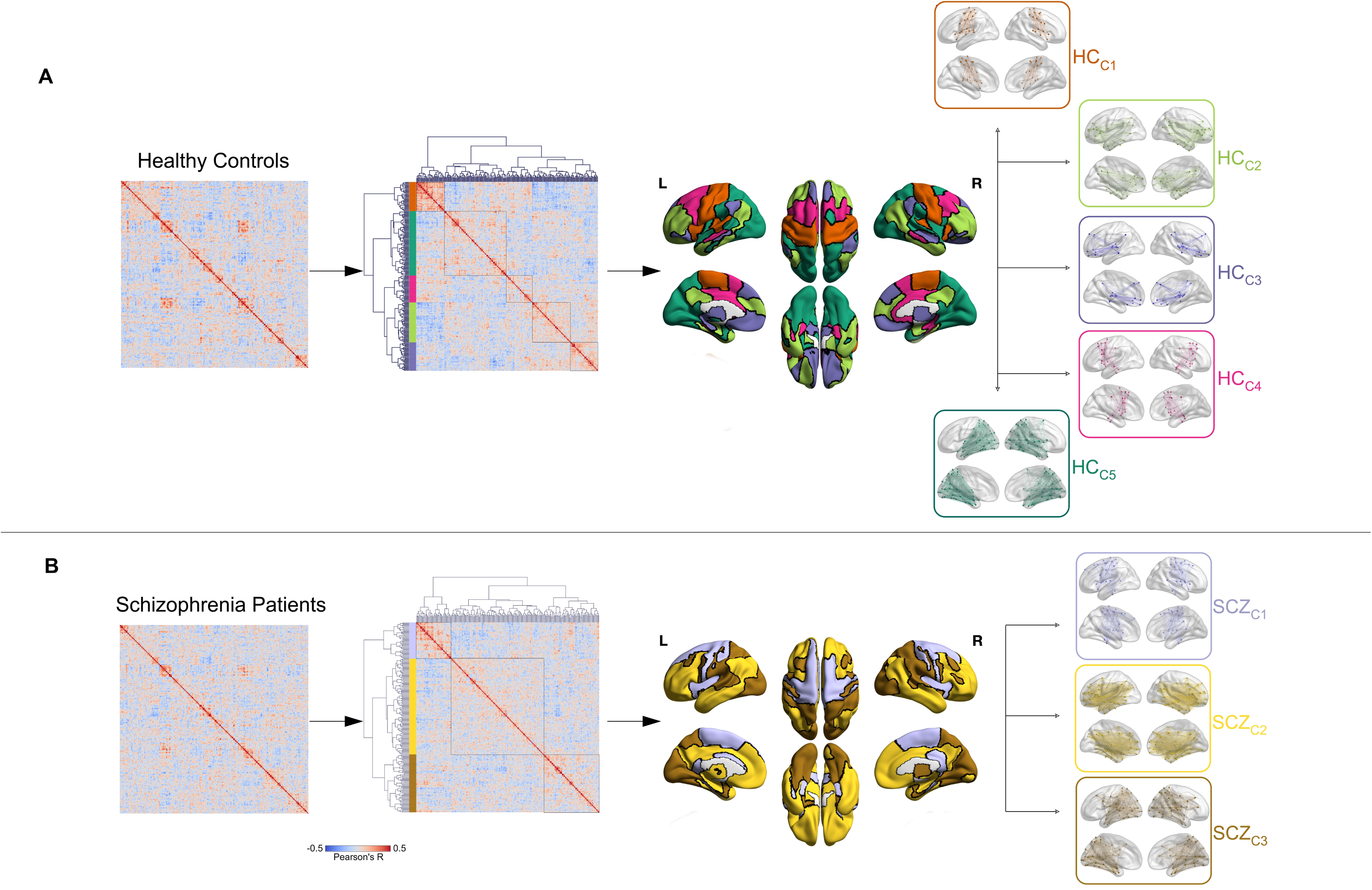
Agglomerative Hierarchical Clustering from Centrality Dynamics. The figure provides an overview of the clustering analysis and the resultant clustering solutions for each of (a) HC and (b) SCZ. In each sub-figure, the heat map of correlation coefficients between all pairs of t_BC_ captures the similarities in the centrality dynamics between all pairs of nodes in the group. These coefficients formed the data subsequently used for clustering. The choice of Agglomerative Hierarchical Clustering (Ward method) was motivated by its use of similarities (i.e., coefficients) as a basis for clustering, it’s robustness in generating clustering solutions independent of cluster size, and its effectiveness in clustering high-dimensional data. The original heat maps are reorganized (to the right) with the order of nodes reorganized to reflect the clustering solution (elbow plots (see Supplementary Figure 2) revealed the optimal numbers of clusters in HC and SCZ to be five and three respectively). Clusters are indicated by color bars on the left of each heat map (the color scheme is maintained for the remainder of the manuscript). Cluster names are based on intra-cluster correlation (where Cluster 1 in each group has highest intra-cluster correlation). The squares on each heat map represent intra-cluster correlations. Dendrograms show the hierarchical relationship between the observed clusters. In each group, the regions assigned to each cluster were then reverse mapped to the cerebral surfaces and the cumulative map is decomposed into separate depictions of each cluster (C_n_). For subsequent analyses, we computed the average cluster time series, t_Cluster_ (across all regions in the cluster) to represent that cluster’s centrality dynamics.

**Figure 3.**
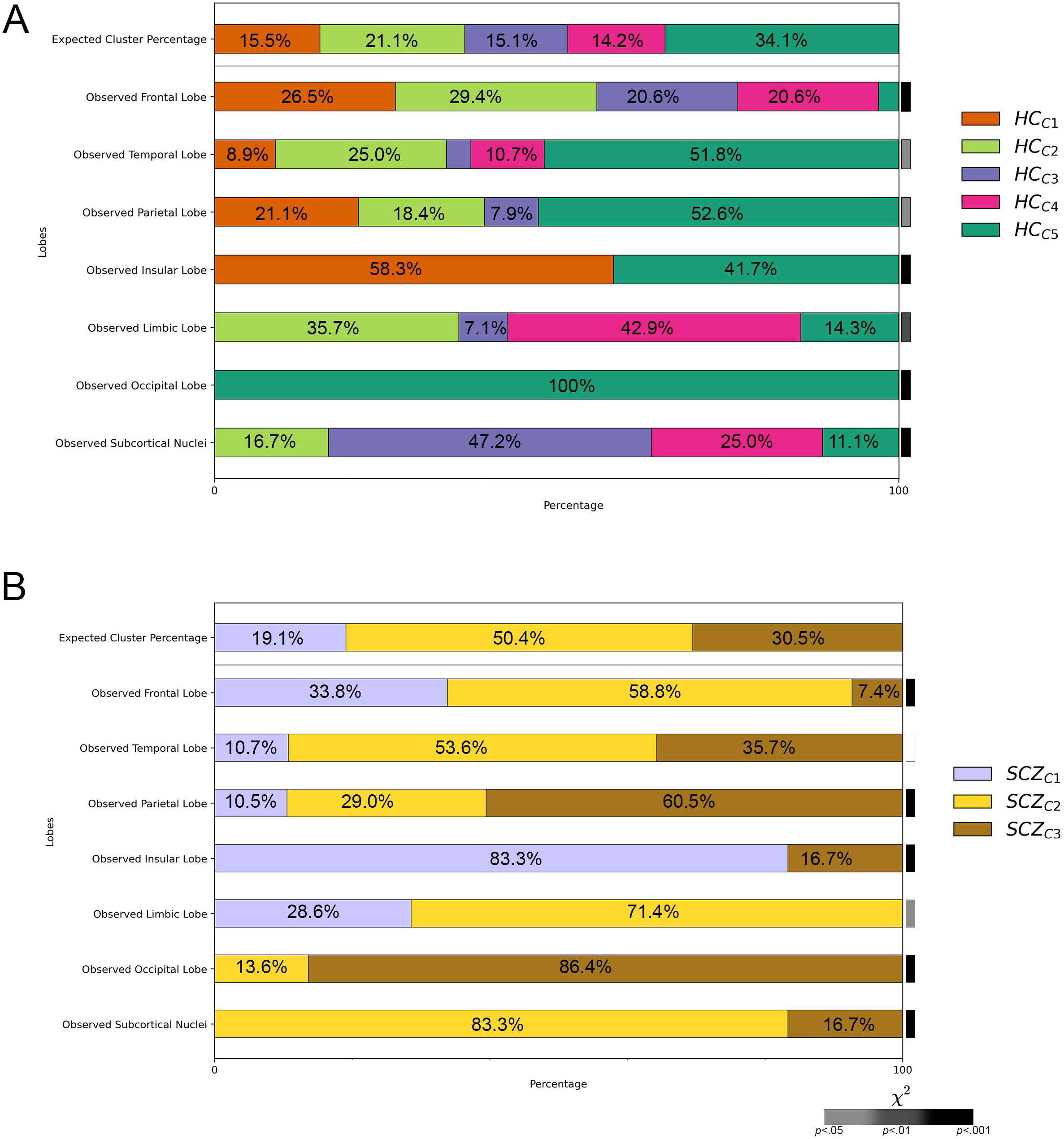
Cluster Composition by Brain Region. For each of (A) HC and (B) SCZ, the top stacked bar graph represents the percentage of nodes (across all lobes/regions) that were assigned to each of SCZ_C1_ – SCZ_C3_ or each of HC_C1_ – HC_C5_. The width of the bar provides a visual representation of the percentage of nodes (out of 246) assigned to each cluster (the actual percentages are noted in the bars). As seen, SCZ_C2_ was the largest cluster in patients (124 nodes, 50.4%) whereas HC_C5_ is the largest in controls (84 nodes, 34.1%). Each top graph is effectively a “null distribution”; thus, if each of the seven lobes/regions contributed proportionally to each cluster, the observed graphs for each region would look identical to the top graph. This representation allows us to inspect several visually compelling trends. For example, the frontal lobe is substantially under-represented in SCZ_C3_. Against an expected percentage of 34.1%, only 7.5% of frontal nodes (5 nodes out of 68) were assigned to this cluster. Conversely, the frontal lobe is over-represented in SCZ_C1_, where against an expected percentage of 19.1%, 33.8% of frontal nodes (23 nodes out of 68) were assigned. These observations become relevant when looking at the Approximate Entropy of the Centrality Dynamics of each of the sub-networks. Chi-Square Tests confirmed that the parcellations significantly diverged from the expected frequencies except for the temporal lobe in SCZ. Each bar on the right of the panel represents the p value of the Chi Square test (see color bar, bottom right for significance).

**Figure 4.**
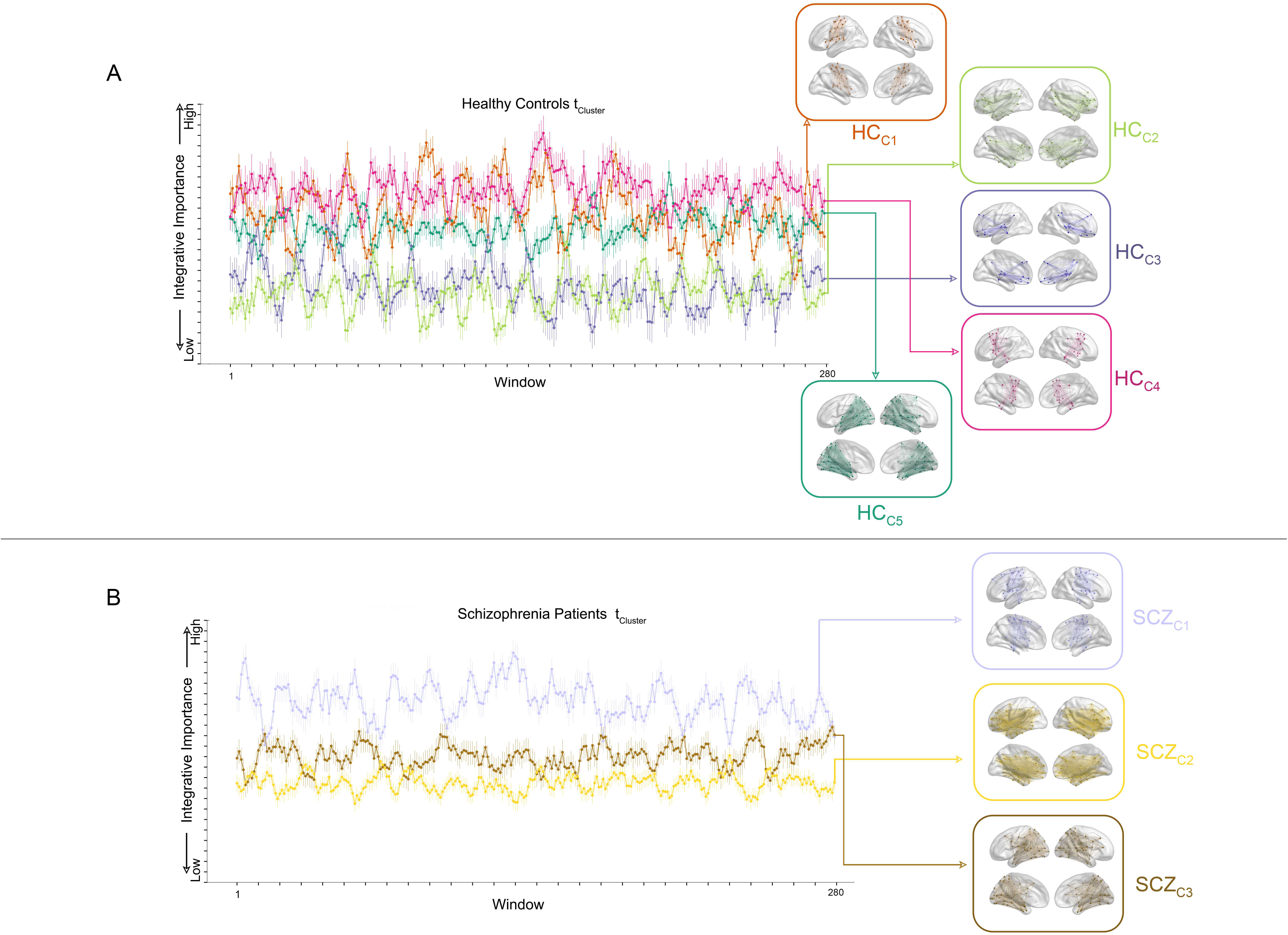
Centrality Dynamics for each Sub-Network. The Figure depicts the five t_Cluster_ in HC (error bars are ± sem). The color conventions are maintained from Figure 2 and 3 and the arrows link each time series with the cluster from which it was derived.

**Figure 5.**
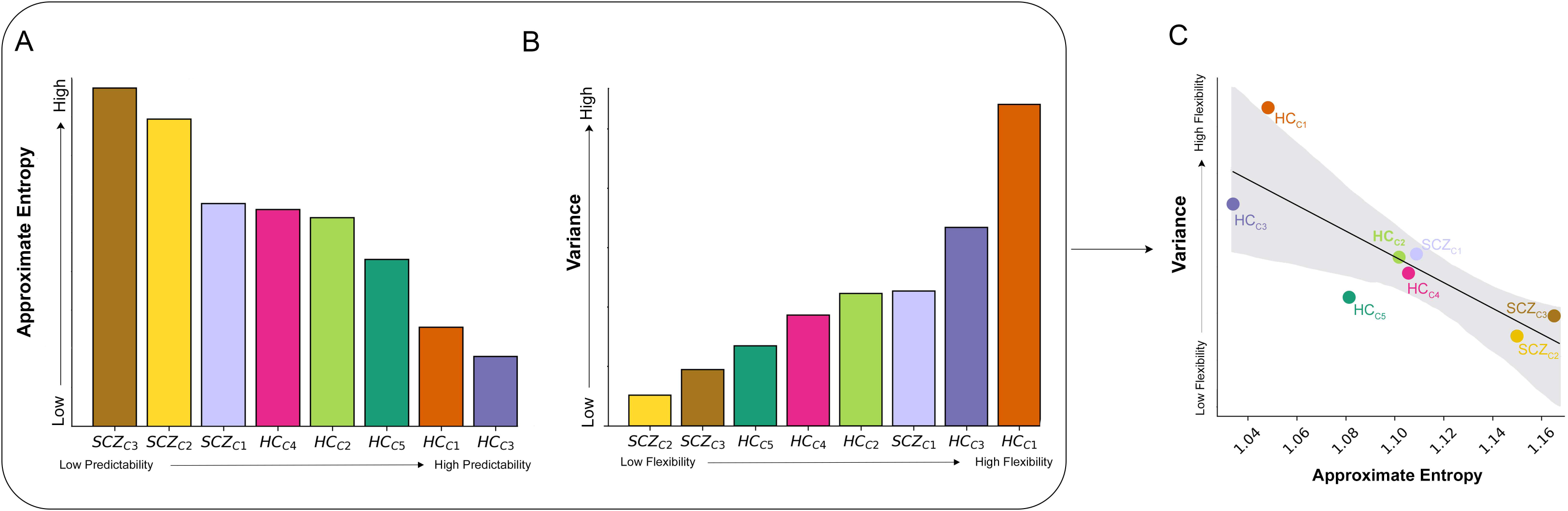
Approximate Entropy and Variance in Centrality Dynamics Across Sub-networks. (A) The Approximate Entropy (ApEn) values for each of the eight clusters (HC_C1_-HC_C5_ and SCZ_C1_-SCZ_C3_) are presented in descending order. ApEn is an index of the regularity or the unpredictability over the fluctuation of a time series; thus, higher ApEn values indicate greater unpredictability (i.e., lower regularity) of a time series. As seen t_Cluster_ in schizophrenia are characterized by higher ApEn values than those in healthy controls, indicating that centrality dynamics in schizophrenia were more irregular. (B) The variance for each of the 8 clusters is presented (calculated across the 280 values in each t_Cluster_, see Methods) in ascending order. A high degree of variability in the centrality dynamics of a time series is indicative of a greater range of dynamics of a sub-network, in turn suggestive of the sub-network’s flexibility over the task. As seen by the order of variance of t_Cluster_, this value was on average lower in SCZ, suggestive of lower sub-network flexibility in patients. (C) The negative relationship between ApEn and variance was highly significant (r^2^=.64, p<.02, shaded area is 95% confidence interval).

**Figure 6.**
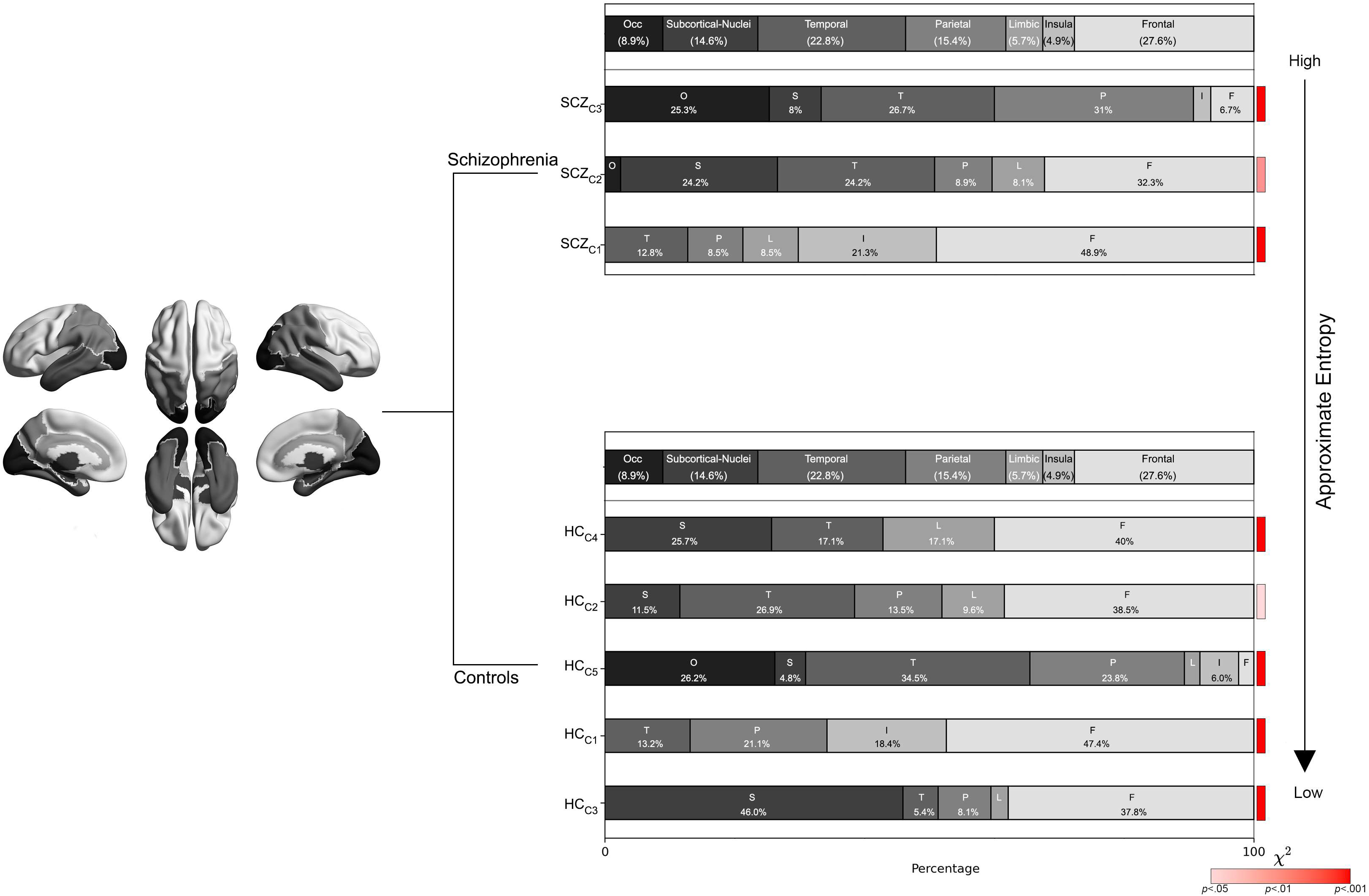
Regional Representation Across Sub-networks. In each panel, the stacked bar graph on the top depicts the actual representation of lobes/region in the cerebral connectome (calculated as a percentage of nodes across the 246-node connectome, with the actual percentage noted in the bars). Thus, in the atlas, frontal nodes constituted the largest set (27.6%). As with Figure 3, each top graph is effectively a “null distribution”; thus, if in each cluster, the representation of lobes/regions conformed to this null distribution, the observed graphs for each cluster would look identical to the top graph. In the figure, the clusters (SCZ_C1_ – SCZ_C3_, HC_C1_ – HCC5) are separated by group, and both within and across groups, are arranged in decreasing order of ApEn. Some of the visually observed trends are intriguing. For instance, in SCZ, the ApEn of the cluster decreases as the relative representation of frontal nodes increases. Moreover, the cluster with the lowest ApEn (HC_C3_) showed an absence of occipital nodes, but substantial over-representation of frontal nodes (37.8%, against an expected percentage of 27.6) and nodes like the thalamus and its sub-regions assigned to sub-cortical nuclei (45.9%, against an expected percentage of 14.6). Each bar on the right of the panel represents the p value of the Chi Square test (see color bar, bottom right for significance).

**Figure 7.**
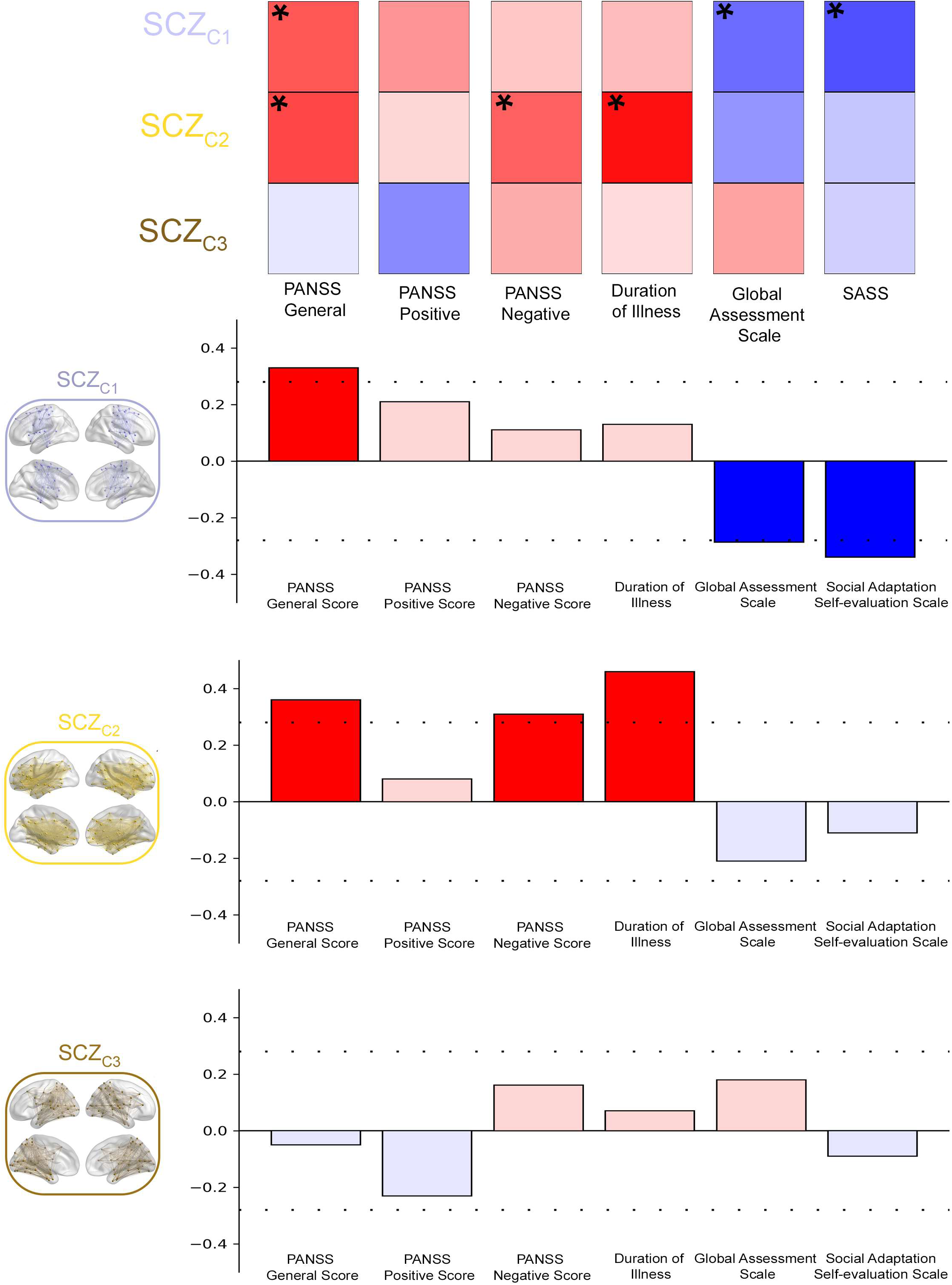
Approximate Entropy of Centrality Dynamics and Clinical Symptoms in Schizophrenia. The figure depicts the exploration of relationship between the entropy of centrality dynamics and schizophrenia clinical symptoms for each of SCZ_C1_-SCZ_C3_. Across the three panels, each bar represents the magnitude and direction of the correlation between the ApEn of t_Cluster_ and clinical measures across 49 SCZ patients. The dashed lines in each panel represent the significance threshold (p < .05). These results indicate increase entropy (greater unpredictability) of centrality dynamic of the sub-networks in SCZ is associated with worse clinical presentation.

### AHC Solutions in HC and SCZ

AHC identified unique solutions for each of the HC and SCZ groups (Fig. 2A and 2B, see Supplementary Fig. 1 for details). The original matrices (left) were numerically ordered by region number in the Brainnetome (28) (and organized by lobe).

Following AHC the matrices are reorganized where the dendrograms depict the resultant clustering structure. Colors assigned to represent each cluster (denoted on the vertical axis) are maintained going forward. Next, the regions in each cluster are projected to lateral, medial, dorsal and ventral cortical surfaces, following which we provide a sub-network representation (here, we depict each node and add edges to complete the representation). As noted, clusters are ordered (HC_C1_ - HC_C5_; SCZ_C1_ - SCZ_C3_) by decreasing intra-cluster correlation strength (see Supplemental Table 1 for a detailed listing of the brain regions assigned to each cluster).

Sub-networks in HC were characterized by greater anatomical diversity (made evident in the stacked bar graphs in Fig. 3). For each of HC and SCZ, the top bar in Fig. 3 represents the percentages of nodes (out of 246) that were assigned to each cluster (HC_C1_-HC_C5_, or SCZ_C1_-SCZ_C3_). If nodes were assigned to each cluster in the same proportion as the expected cluster percentage, then each subsequent bar graph would be identical to the top bar. The top bar therefore effectively serves as a null distribution of the expected relative frequencies of nodes. We observed several compelling trends. Nodes in the seven lobes/regions were not proportionally distributed across clusters. Thus, for example, in SCZ_C3_ the frontal lobe is notably *under-represented* (only 7.5% of nodes assigned, compared to an expected percentage of 19.1%). Conversely, in SCZ_C1_ the frontal lobe is *over-represented* (33.8% of nodes were assigned to this cluster). Chi-Square Tests confirmed that the observed parcellations significantly diverged from the expected frequencies (with the sole exception of the temporal lobe in SCZ).

### Approximate Entropy and flexibility of sub-networks

Next, across regions, mean centrality dynamics were represented in time series (t_cluster_) (Fig. 4). In HC the centrality dynamics evinced a) greater periodicity (evidence of lower stochasticity) and b) greater amplitude variation, suggesting greater flexibility to task-induced changes (44). The latter observation is notable because high variability is a feature of adaptive macroscopic brain networks (45), where common patterns of variability are associated with common inter-regional network dynamics.

We used ApEn to quantify the predictability of centrality dynamics specifically because it is well suited for studying shorter time series (46). We expect that in any t_Cluster_, stochastic patterns will have *low predictability* and *high ApEn*, but periodic patterns will have *high predictability*, and *low ApEn*.

Fig. 5A presents the observed ApEn (arranged in descending order of magnitude) for each t_Cluster_ in HC and SCZ. All three SCZ sub-networks evinced higher ApEn values (lower predictability). This observation formalizes the visual intuitions from Fig. 3.

Next (as shown in Fig. 5B), sub-network flexibility in SCZ was observed to be generally lower than in HC (in the graph the sub- networks are ordered in ascending order of variance). In exploratory analyses, we investigated *whether higher ApEn (less predictability) was associated with lower variance (less flexibility)*. Indeed, we observed a highly significant negative relationship (r²=.64, *p*<.02) across the eight sub-networks (Fig. 5C). This negative relationship indicates *that as the stochasticity of the t_cluster_ of sub-networks increased, the sub- network flexibility decreased*. In other words, sub-networks with a high degree of stochasticity of centrality dynamics were the ones with a low degree of flexibility.

Are regions/lobes evenly represented in each sub-network? The top bar in Fig. 6 depicts a null distribution constructed from the base rate of node assignments in the overall connectome. Thus, if node assignment to sub-networks conformed to the base rate, the distribution in each subsequent bar would emulate the null distribution. Subsequent bars depict the observed assignments of nodes to each sub-network (percentages are indicated) and sub-networks are arranged in decreasing order of ApEn (i.e., as centrality dynamics become increasingly periodic). We observed some intriguing visual trends. For example, SCZ_C1_, which has a larger than expected percentage of frontal nodes shows greater periodicity in centrality dynamics (relative to SCZ_C2_ and SCZ_C3_), whereas HC_C3_ which exhibits the highest periodicity in centrality dynamics sees a heavy over- representation of sub-cortical nodes such as the thalamus, as well as frontal nodes.

Is the ApEn of Centrality Dynamics in patients associated with clinical features? We addressed this question by first calculating ApEn from each patient’s data. That is, from each of SCZ_C1_-SCZ_C3_ and in each patient (n=49), we estimated the mean t_BC_ (averaging across all t_BC_ for nodes in the cluster). Next, ApEn was calculated for this mean t_BC_ (See Supplementary Fig. 3 for the methodological pipeline). We related these ApEn values to clinically salient measures including the PANSS General Score, the PANSS negative and positive scales (34), the Global Assessment Scale (47), the Social Adaptation Self-evaluation Scale (48) and duration of illness (49).

Fig. 7 depicts the significant results (as well as the direction of the relationship). Notably, the directionality of effects was highly consistent: regardless of the sub-network, increased ApEn of centrality dynamics was associated with poorer clinical presentation. This was true for SCZ_C1_ (PANSS General, GAS and SASS), and SCZ_C2_ (PANSS General, PANSS negative and DUI). *Thus, more irregular centrality dynamics were predictive of the greater poverty of the clinical state*.

## Discussion

In this paper we motivated the construct of task-driven Centrality Dynamics a measure that quantifies dynamic changes in the integrative importance of a node across a task (50). CD is based on the graph theoretic characterization of the betweenness centrality (BC) of nodes in a given functional connectome (18), where each such connectome is estimated in a succession of overlapping moving windows. Each node’s constructed time series captures dynamic changes in its’ BC. Because network connectomics will dynamically change in response to the task, fluctuations in BC are readily interpretable as lucid representations of such dynamics. These time series were partitioned (based on the similarities in the centrality dynamics) into non-overlapping sub-networks. Therefore, each sub-network was composed of nodes with homogenous centrality dynamics. Our scientific questions of interest were these. If connectomic dynamics are altered in schizophrenia: 1) Will the constituted sub-networks in SCZ be different from those in HC? 2) How predictable and how variable were the centrality dynamics of sub-networks in SCZ? 3) Were centrality dynamics predictive of clinical features in SCZ? These points are discussed in the remainder of the Discussion.

### Composition of the recovered sub-networks

Cognitive optimization is driven by the brain’s ability to dynamically parcellate itself into sub-networks (51). These functional modules (often “rich clubs”) allow sub-networks to communicate their outputs (to other networks) through hubs (52, 53). The dynamics of these sub-networks have largely (though not entirely) been derived from resting state fMRI time series data (54, 55). As the analytical focus shifts to the study of dynamics, the use of advanced models assumes greater importance. Here, such models should capture network dynamics and quantify how differential dynamics might drive sub-network segregation (56, 57). Our work fills this space because it elucidates the interjection between a) task-induced dynamics in b) the functional connectome and c) the nature of recovered sub-networks in heath and schizophrenia.

In this vein, our methods render our findings novel, and the outcomes of our analyses are straightforwardly recognizable. We observed greater sub-network parcellation in HC (HC_C1_ – HC_C5_) than in SCZ (SCZ_C1_ – SCZ_C3_) (see Fig. 3 & 6), and the sub-networks in HC were composed of regions with greater functional homogeneity. This effect indicates that in healthy participants, brain regions with greater functional homogeneity had more similar centrality dynamics. For instance, HC_C1_ is largely comprised of regions in the sensorimotor cortex (58), HC_C2_ of regions in the frontal, temporal, and parietal lobes (59–61), HC_C3_ of regions in the prefrontal cortex, hippocampus, and thalamus (62–64), and HC_C5_ almost exclusively of regions in the magnocellular and parvocellular forward visual pathways (65–67).

This pattern of relative homogeneity was violated in the observed sub-networks in SCZ. Here, sub-networks were composed of brain regions with greater functional heterogeneity. For instance, SCZ_C2_ is composed of regions from a cross-section of brain areas including the frontal, limbic, and temporal lobes, as well as regions in subcortical nuclei (61). Furthermore, SCZ_C3_ is a non-specific superset of regions that includes (but is not restricted to the magno- and parvocellular visual pathways; see HC_C5_ for comparison). Therefore, in SCZ, it appears that functionally heterogenous regions showed more similar centrality dynamics, an effect that is generally consistent with previous evidence showing a lack of network segregation in schizophrenia(68). The increase in heterogeneity is emphasized in the insets in each of the re-ordered adjacency matrices in Fig. 2a (HC) and 2b (SCZ). As seen, the r values (representing the similarities in centrality dynamics between regions) are more homogenous in HC than in SCZ.

Are the network properties captured by centrality dynamics capture distinct from those observed when analyzing fMRI time series data? In supplementary analyses we verified that the obtained results based on CD were wholly or partially distinct from those obtained from fMRI time series data. fMRI time series were clustered based on the similarity in their profiles and AHC was conducted (again separately in HC and SCZ) after computing 30,135 correlation coefficients between pairs of regions (representing the similarities in fMRI fluctuations). These analyses identified five and four unique clusters in HC and SCZ respectively (see Supplementary Fig. 4-6 and Supplementary Table 1-2 for more details), whose solutions were complementary to those observed in the primary analyses based on CD. In ongoing analyses, we are attempting to quantify similarities and differences in these discovered networks.

### Stochasticity and Flexibility in Schizophrenia

Not only did our primary analyses recover sub-networks in schizophrenia where the regions were more functionally heterogeneous, but the centrality dynamics of these sub-networks were also characterized by less predictability and lower amplitude variations. ApEn is a non-linear complexity measures that a) can lucidly classify complex system characteristics (69) and b) requires few assumptions about any putative neural processes underlying the data (70). More generally, entropy suitably characterizes stochastic biological data because unlike moment statistics, entropy measures do not depend on absolute or measured values of the signal. ApEn was originally designed to distinguish between a variety of systems from the nature of their time series outputs. The classes of systems that can be discriminated between include low-dimensional deterministic systems, periodic and multiply periodic systems, and high-dimensional chaotic, stochastic, and mixed systems (69, 71).

The centrality dynamics of the sub-networks in SCZ evinced increased stochasticity, an observation that suggests that the connectomic changes *were not systematically responsive to task effects*. Again, this is a recognizable generalization (to the network scale), of previously observed results; Schizophrenia has been associated with reduced signal-to-noise ratio in brain signals, that in turn drives sporadic and unpredictable statistical fluctuations in multiple cortical brain networks.

Therefore, the internal workings of the schizophrenia brain satisfy some of the hallmarks of stochastic systems (72, 73). While stochastic systems typically evince greater amplitude fluctuations as well, this was not true in our data. Greater stochasticity did not result from large amplitude fluctuations in the centrality of nodes. Contrarily, the amplitude range of centrality dynamics in schizophrenia was low (as evidenced in the observation of lower variance) (see Fig. 5). Effectively then, centrality dynamics in the schizophrenia brain display a high degree of stochasticity but inflexibility to task-induced change. Whereas dynamic instability is a form of complexity typical of neuronal systems and is crucial for adaptive brain function (74–76), the dynamics of the connectome in the schizophrenia brain is unstable and inflexible.

As previously noted, (Methods), window width was motivated by the condition width which means that our dynamic analyses coincided with the repeating nature of the task structure (see Fig. 1). Are the observed centrality dynamics statistically predicted by task dynamics? We conducted additional supplemental analyses in which we related the average centrality dynamics of each cluster (t_Cluster_) to the objective dynamics of the task.

Here, we used wave forms to represent each of the four task conditions, t_Condition_; Each constructed ordinal function represents a condition’s contribution at each time point (where a value of 0 reflects no contribution of that condition at that time point, whereas a value of 9 reflects full contribution of that condition at that time point; see Supplementary Fig. 7a for the composite function, and Supplementary Fig. 7b for its decomposition). Then, for each of HC_C1_ – HC_C5_ and SCZ_C1_ – SCZ_C3_, we examined correlations between each t_Cluster_ and each t_Condition_. *Any significant relationships would indicate that a cluster’s centrality dynamics were predicted by condition dynamics*. These analyses revealed thirteen significant effects in HC (out of a possible 20: five clusters and four conditions) but only five in SCZ (out of a possible 12: three clusters and four conditions, see Supplementary Fig. 8-11). These results indicate that unlike in typical controls, *centrality dynamics in schizophrenia are largely unyoked from the ongoing demands of the task*.

### Centrality dynamics and clinical symptoms

Because schizophrenia’s core symptoms are not directly captured in imaging data, the relationships between these symptoms and cognitively driven imaging measures become relevant (77, 78).

Here, our study of the relationship between entropy measures of centrality dynamics and clinical symptoms (Fig. 7) were highly revealing. Regardless of sub-network parcellation, the increased stochasticity of centrality dynamics (increased ApEn) predicted poorer clinical presentation (whether based on the cumulative score on the Positive and Negative Syndrome Scale for Schizophrenia, PANSS). This result indicates that more stochastic dynamics of the schizophrenia connectome are related to worse clinical presentation in schizophrenia (79).

### Limitations and Conclusions

Sentient brains do not rest (80), and are characterized by incessant and spontaneous dynamics (6). However, fMRI signals are also highly sensitive to task-based contextual modulation (12, 81), because tasks evoke unique network states. Task-based studies are particularly valuable in studying schizophrenia, precisely because induced network perturbations expand the evidentiary range of underlying network deficits and permit better classification and diagnosis (82–84). Our investigations are responsive to these issues, and our results suggest that task-evoked centrality dynamics may complement spontaneous brain fluctuations (85). However, our study inherits many limitations inherent in fMRI-based connectomics. Graph theoretic constructs like Betweenness Centrality have obscure neurophysiological bases because they are not derived from the physiological drivers (e.g., neuronal spiking or low or high frequency neuronal oscillations) that drive fMRI signals (86, 87). Still, the brain’s functional states span multiple spatial and temporal scales (88), and this very ubiquity motivates the development of concepts like centrality dynamics. Future studies should further elucidate the brain’s task-evoked and spontaneous centrality dynamics to better understand the dynamics of dysconnection in schizophrenia.

## Data Availability

The original EPI data have been deposited in the NIH repository (MH111177) and can be freely accessed. The code used to generate these results, and the processed data are freely available upon request.

## Funding

This work was supported by the National Institutes of Mental Health (MH111177), the Cohen Neuroscience Endowment, the Dorsey Endowment, the Ethel & James Flinn Foundation, and the Lycaki-Young Funds from the State of Michigan. The authors declare that there are no conflicts of interest to disclose.

## Supplementary Figure Legends

Supplementary Figure 1. The behavioral data are provided for each participant in each of the HC and SCZ groups. Each heat map represents data from each group with participants arranged in rows and epochs in columns. The color represents the retrieval proficiency in the epoch (see color bar).

Supplementary Figure 2. The figure depicts the results from the agglomerative clustering separately conducted in each of the a) HC and b) SCZ groups based on the t_BC_. The optimal cluster solution was derived from the convergence of the Elbow Plots (left panels) and the Threshold vs Number of Cluster Plot (right panels). In the former case, the optimal cluster solution is conventionally based on a deceleration in the change in the slope of the function. As seen (arrows), this deceleration is observed after a cluster solution of size five in HC and of three in SCZ. Our choice is confirmed by the Threshold vs Number of Cluster Plot where longer vertical lines for each solution size reflect greater stability in the cluster solution.

Supplementary Figure 3. The figure depicts the pipeline used to derive relationships between clinical measures (in schizophrenia) and ApEn. In each cluster (panel A) and for each patient, across all n nodes in the cluster (panel B), we first estimated the mean t_BC_ for each patient (panel C). Finally, we estimated the ApEn for each patient’s mean tBC (49 ApEn values for each of SCZ_C1_ – SCZ_C3_). We then examined the statistical relationship between these ApEn values and clinical measures (panel D).

Supplementary Figure 4. The figure provides the results of the clustering analysis (based on the fMRI time series) for each of (a) HC and (b) SCZ. In each sub-figure, at left, the heat map of correlation coefficients captures the similarities in the fMRI signal across the task between all unique pairs of nodes (30,135 pairs) for each group. These coefficients formed the data subsequently used for Agglomerative Hierarchical Clustering. The original heat maps are reorganized (to the right) with the order of nodes reorganized to reflect the clustering solution (the elbow plots shown in Supplementary Fig. 5 reveal the optimal numbers of clusters in HC and SCZ to be five and four respectively). Color bars at the left of these heat maps denote cluster identity. Dendrograms show the hierarchy of the observed clusters. In each group, the regions assigned to each cluster were then reverse mapped to the cerebral surfaces and the cumulative map is decomposed into separate depictions of each cluster (e.g., HC_fMRIn_).

Supplementary Figure 5. Optimal cluster solutions are depicted for the agglomerative clustering (based on fMRI time series) separately conducted in each of a) HC and b) SCZ (see Supplementary Fig. 4). The optimal cluster solution was derived from the convergence of the Elbow Plots (left panels) and the Threshold vs Number of Cluster Plot (right panels). In the former case, the optimal cluster solution is conventionally based on a deceleration in the change in the slope of the function. As seen (arrows), this deceleration is observed after a cluster solution of size five in HC and of four in SCZ. Our choice is confirmed by the Threshold vs Number of Cluster Plot where longer vertical lines for each solution size reflect greater stability in the cluster solution.

Supplementary Figure 6. A direct comparison of the cluster partitions based on a) Centrality Dynamics and b) fMRI time series emphasized distinct solutions. This visual observation is further supported when comparing regional composition for both clusters formed based on centrality dynamics (Supplementary Table 1) and fMRI time (Supplementary Table 2).

Supplementary Figure 7. The task dynamics were represented by creating temporal functions for each of the four task conditions (t_Condition)_. In each function, we represent the degree to which that condition is represented at any point in time. This value ranged from 0 (i.e., there was no representation of that condition at that point in time) to 9 (i.e., at that point in time that was the only condition represented). The formed condition-related time series are depicted in Panel B and are superimposed on the same timeline in Panel A.

Supplementary Figure 8. The figure explores the statistical relationships between the centrality dynamics of each cluster (t_Cluster_) and the t_Condition_ for Encoding. For each of HC_C1_ – HC_C5_ and SCZ_C1_ – SCZ_C3_, we overlay each t_Cluster_ on each of t_Condition_. Significant relationships (p < 0.05, Bonferroni corrected) between t_Cluster_ and the t_Condition_ for Encoding are depicted. The scatter plots (far right) depict average centrality measures in each of eight epochs (error bars are ± sem). The fitted function is the line of best linear fit. Across clusters, we observed multiple significant relationships between centrality dynamics and task conditions. This scheme of presenting effects is carried forward in Supplementary Figures 9 – 11.

Supplementary Figure 9. The figure explores the statistical relationships between the centrality dynamics of each cluster (t_Cluster_) and the t_Condition_ for Post-Encoding Consolidation.

Supplementary Figure 10. The figure explores the statistical relationships between the centrality dynamics of each cluster (t_Cluster_) and the t_Condition_ for Retrieval.

Supplementary Figure 11. The figure explores the statistical relationships between the centrality dynamics of each cluster (t_Cluster_) and the t_Condition_ for Post-Retrieval Consolidation.

## Notes

### Competing Interest Statement

The authors have declared no competing interest.

