## Supplementary Figures and Tables for "Learning evoked centrality dynamics in the schizophrenia brain: Entropy, heterogeneity and inflexibility of brain networks"

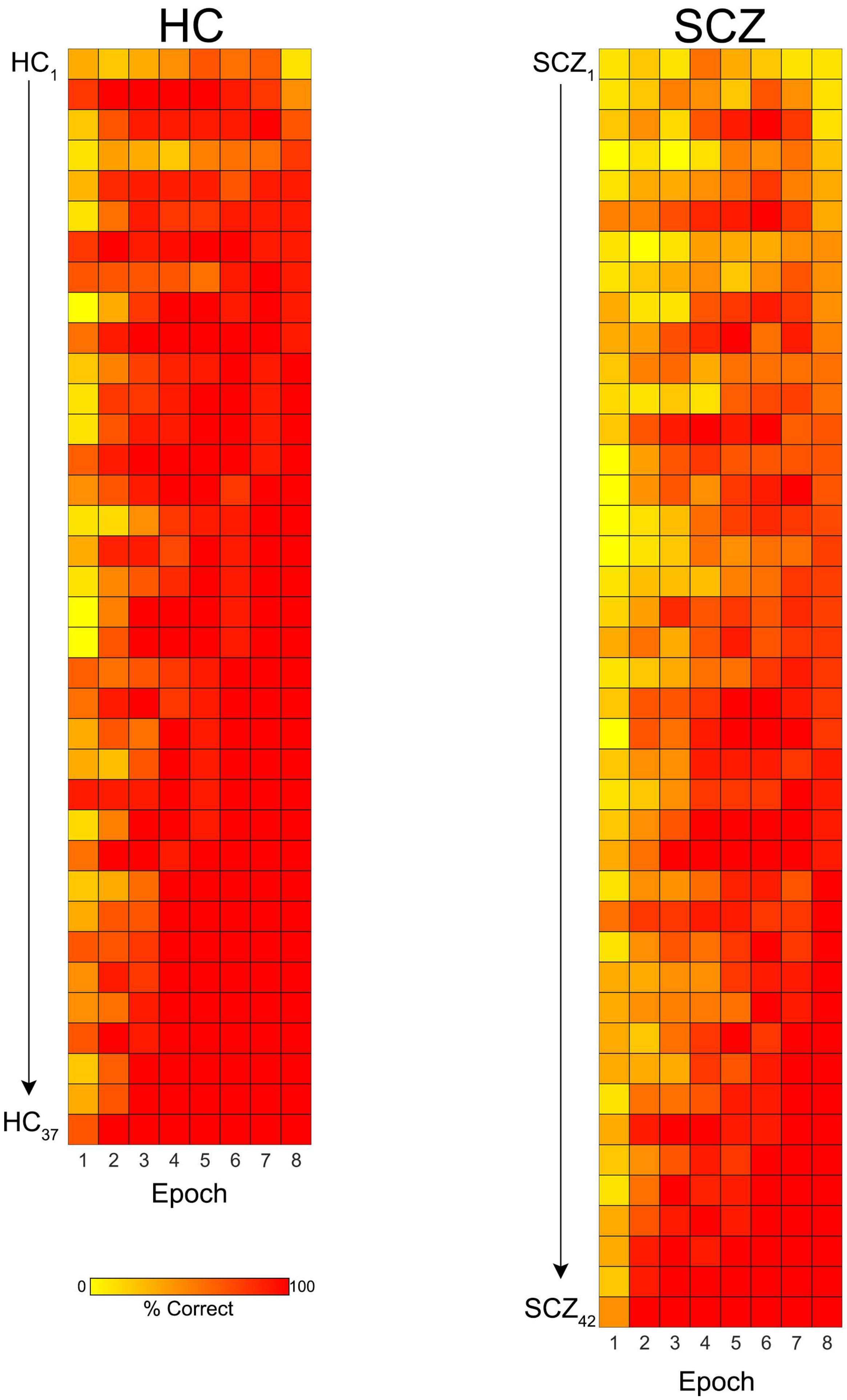


The behavioral data are provided for each participant in each of the HC and SCZ groups. Each heat map represents data from each group with participants arranged in rows and epochs in columns. The color represents the retrieval proficiency in the epoch (see color bar).

Supplementary Figure 2

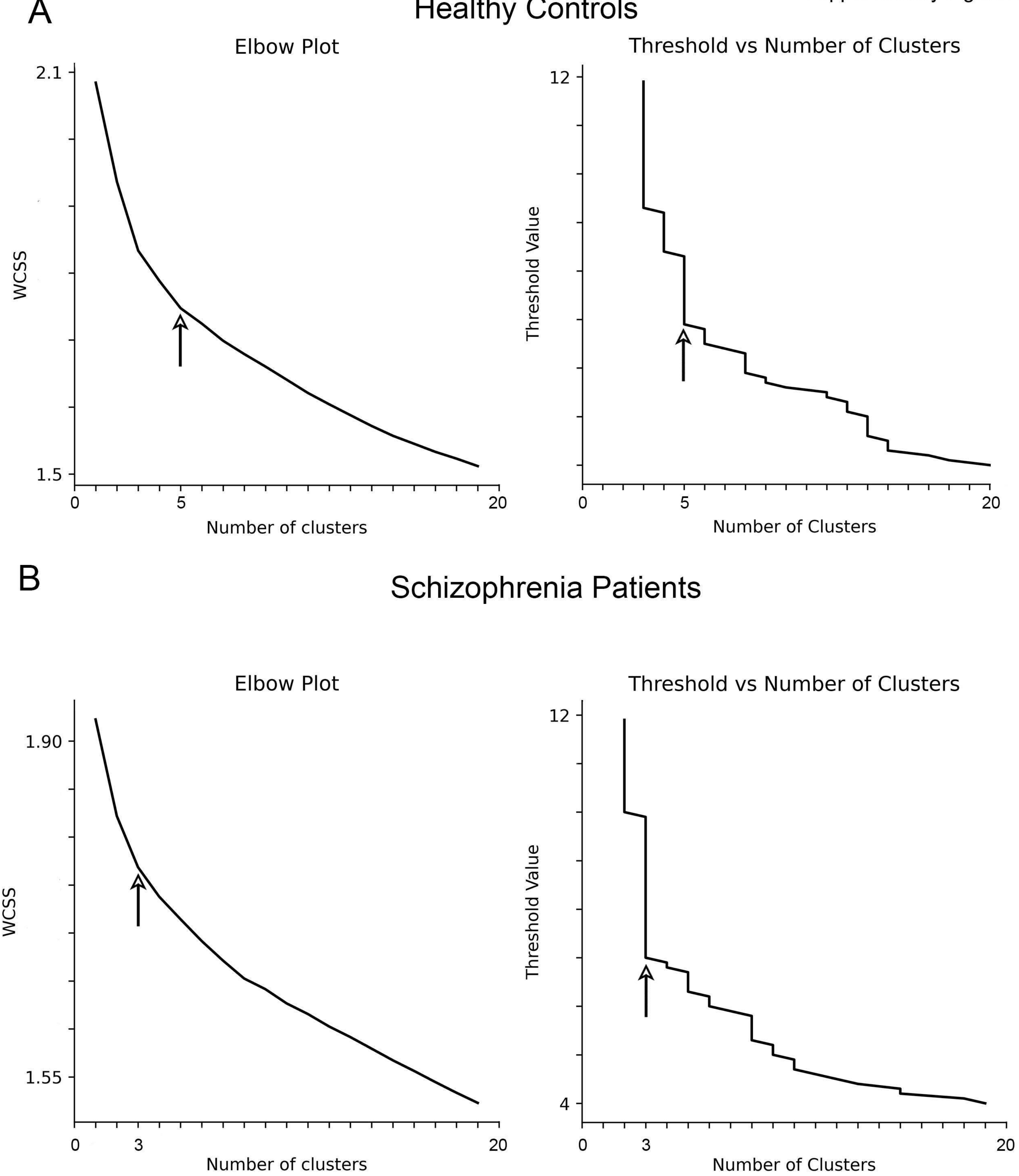


The figure depicts the results from the agglomerative clustering separately conducted in each of the a) HC and b) SCZ groups based on the $t_{BC}$. The optimal cluster solution was derived from the convergence of the Elbow Plots (left panels) and the Threshold vs Number of Cluster Plot (right panels). In the former case, the optimal cluster solution is conventionally based on a deceleration in the change in the slope of the function. As seen (arrows), this deceleration is observed after a cluster solution of size five in HC and of three in SCZ. Our choice is confirmed by the Threshold vs Number of Cluster Plot where longer vertical lines for each solution size reflect greater stability in the cluster solution.

Supplementary Figure 3

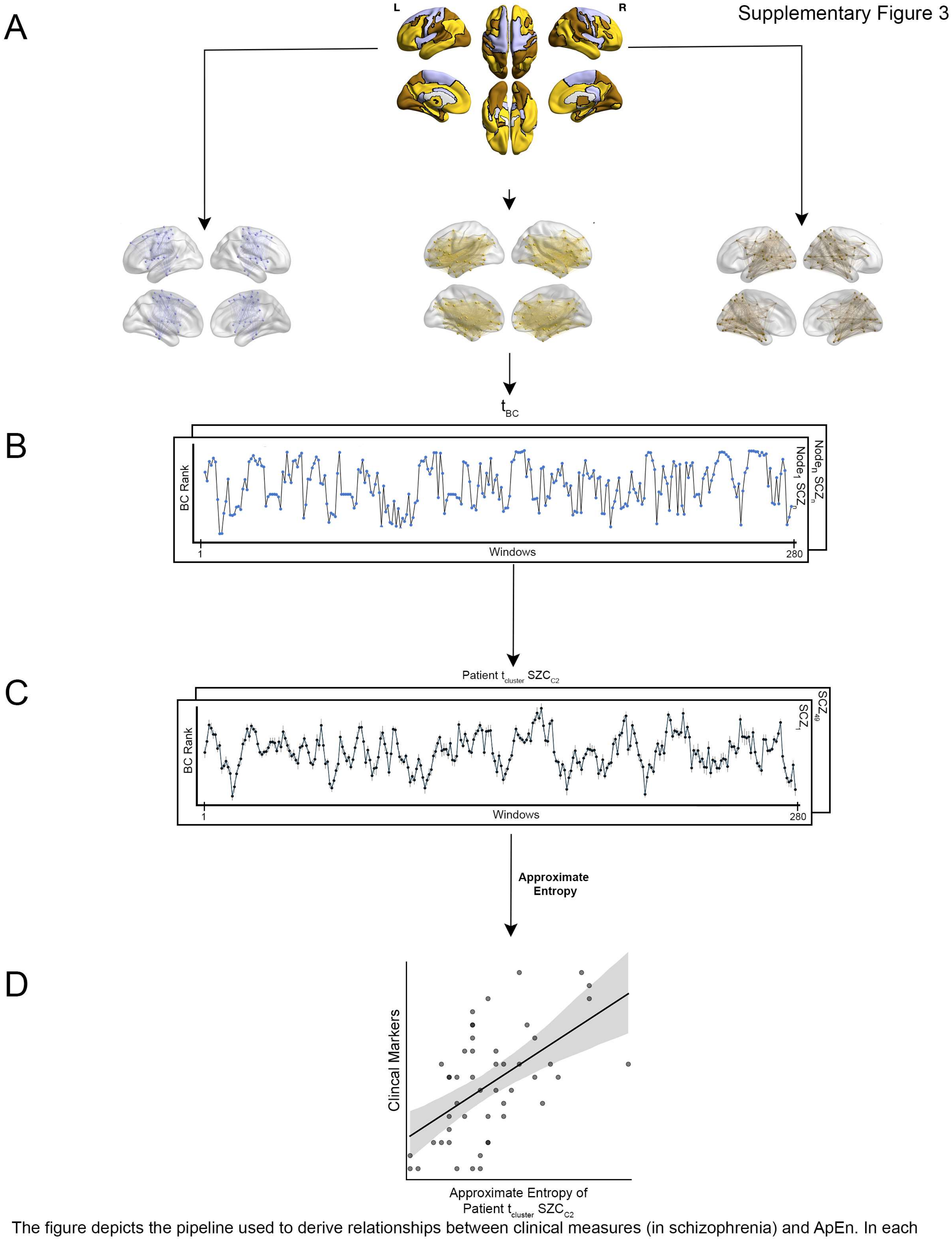


The figure depicts the pipeline used to derive relationships between clinical measures (in schizophrenia) and ApEn. In each cluster (panel A) and for each patient, across all n nodes in the cluster (panel B), we first estimated the mean $t_{BC}$ for each patient (panel C). Finally, we estimated the ApEn for each patient's mean $t_{BC}$ (49 ApEn values for each of SCZC1 – SCZC3). We then examined the statistical relationship between these ApEn values and clinical measures (panel D).

Supplementary Figure 4

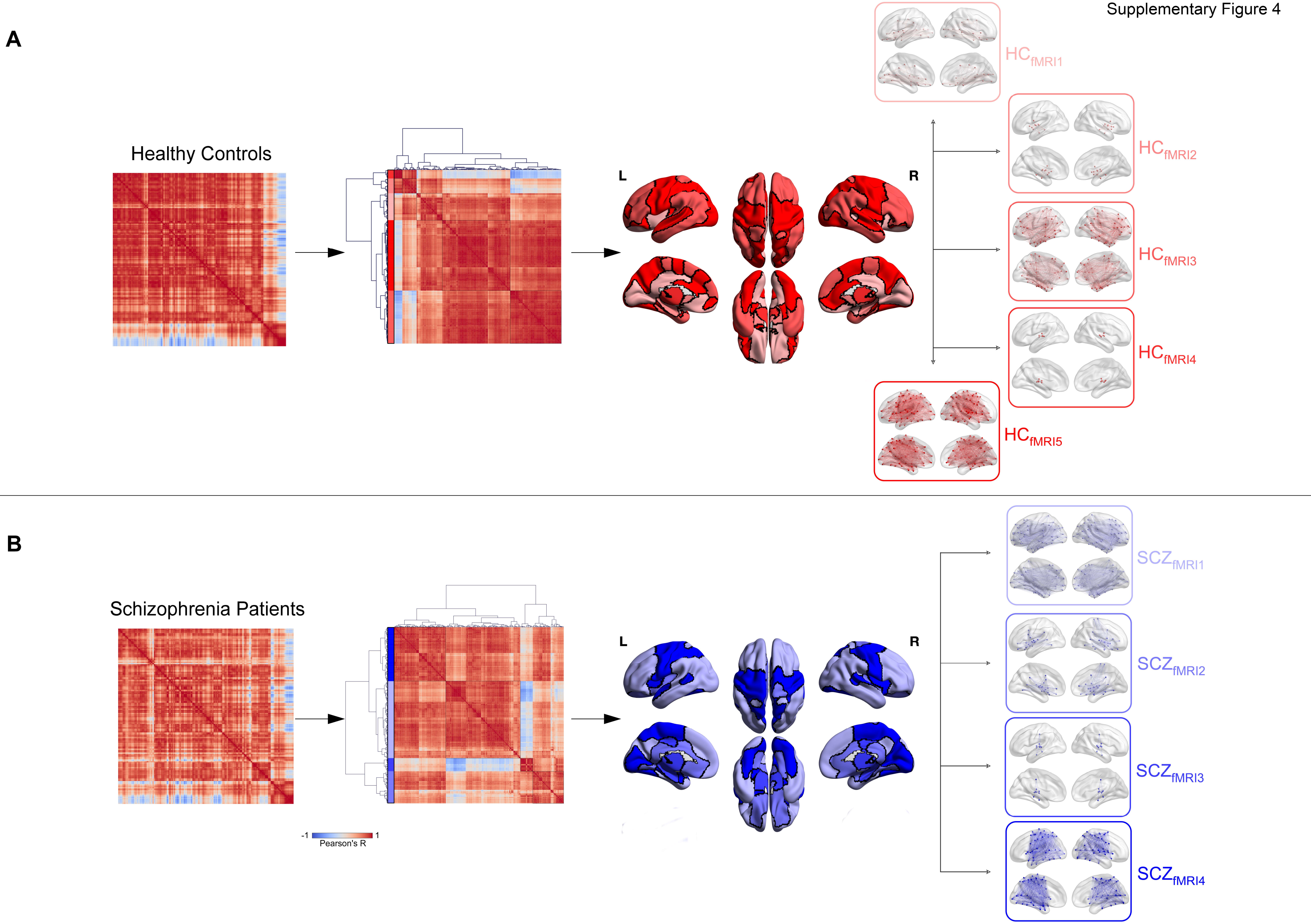


The figure provides the results of the clustering analysis (based on the fMRI time series) for each of (a) HC and (b) SCZ. In each sub-figure, at left, the heat map of correlation coefficients captures the similarities in the fMRI signal across the task between all unique pairs of nodes (30,135 pairs) for each group. These coefficients formed the data subsequently used for Agglomerative Hierarchical Clustering. The original heat maps are reorganized (to the right) with the order of nodes reorganized to reflect the clustering solution (the elbow plots shown in Supplementary Fig. 5 reveal the optimal numbers of clusters in HC and SCZ to be five and four respectively). Color bars at the left of these heat maps denote cluster identity. Dendrograms show the hierarchy of the observed clusters. In each group, the regions assigned to each cluster were then reverse-mapped to the cerebral surfaces and the cumulative map is decomposed into separate depictions of each cluster (e.g., $HC_{fMRIn}$).

Supplementary Figure 5

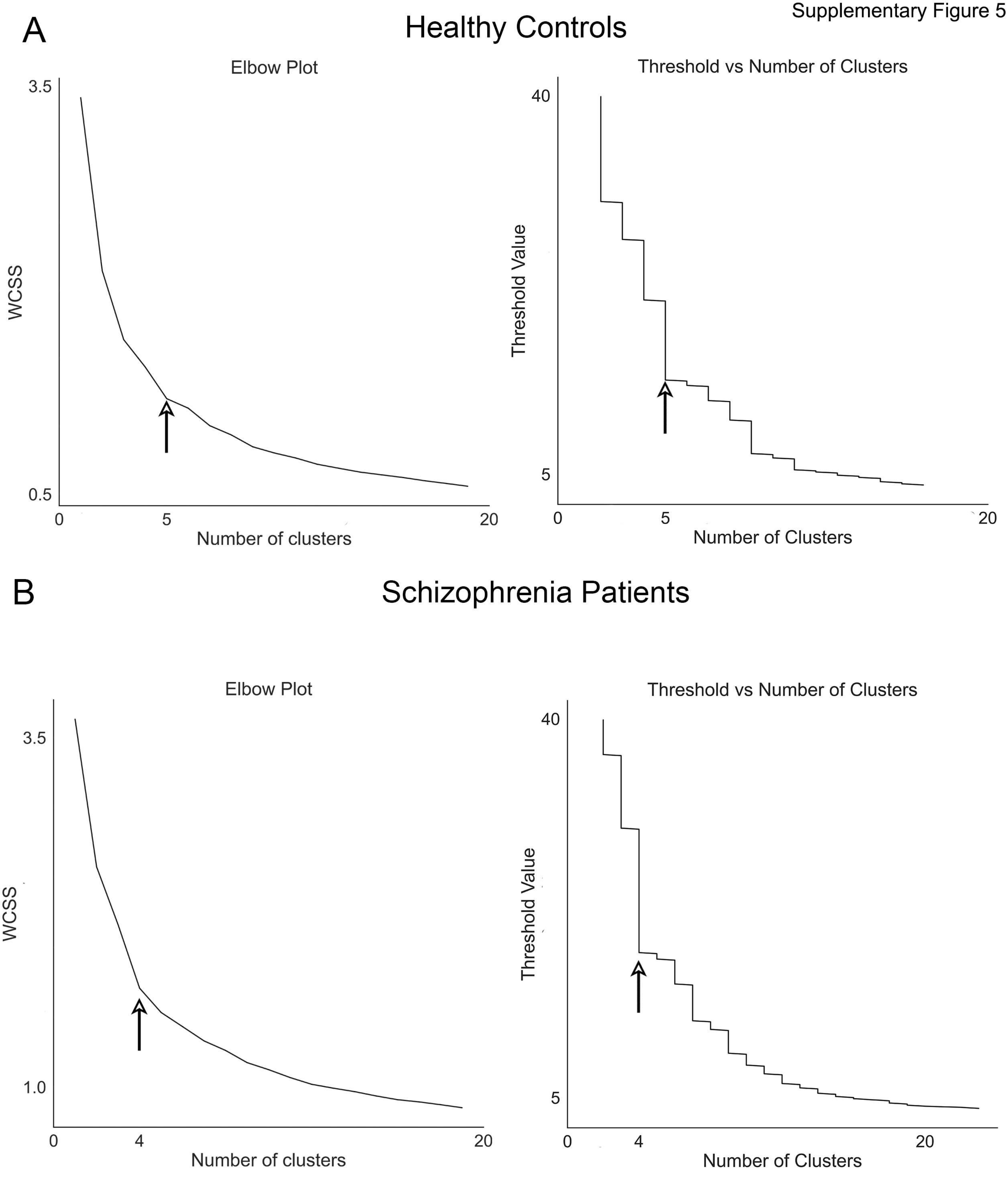


Optimal cluster solutions are depicted for the agglomerative clustering (based on fMRI time series) separately conducted in each of a) HC and b) SCZ (see Supplementary Fig. 4). The optimal cluster solution was derived from the convergence of the Elbow Plots (left panels) and the Threshold vs Number of Cluster Plot (right panels). In the former case, the optimal cluster solution is conventionally based on a deceleration in the change in the slope of the function. As seen (arrows), this deceleration is observed after a cluster solution of size five in HC and of four in SCZ. Our choice is confirmed by the Threshold vs Number of Cluster Plot where longer vertical lines for each solution size reflect greater stability in the cluster solution.

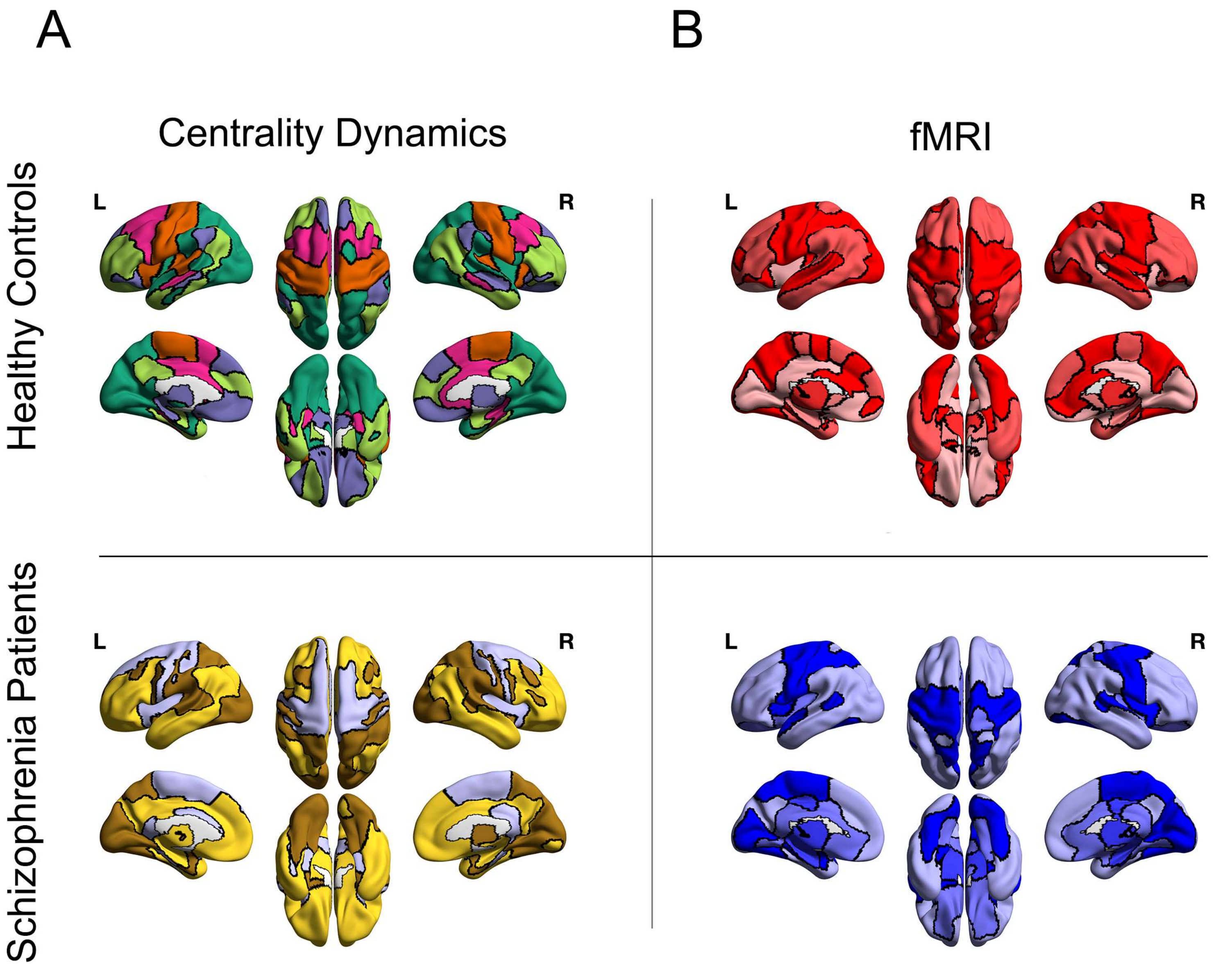


A direct comparison of the cluster partitions based on a) Centrality Dynamics and b) fMRI time series emphasized distinct solutions. This visual observation is further supported when comparing regional composition for both clusters formed based on centrality dynamics (Supplementary Table 1) and fMRI time (Supplementary Table 2).

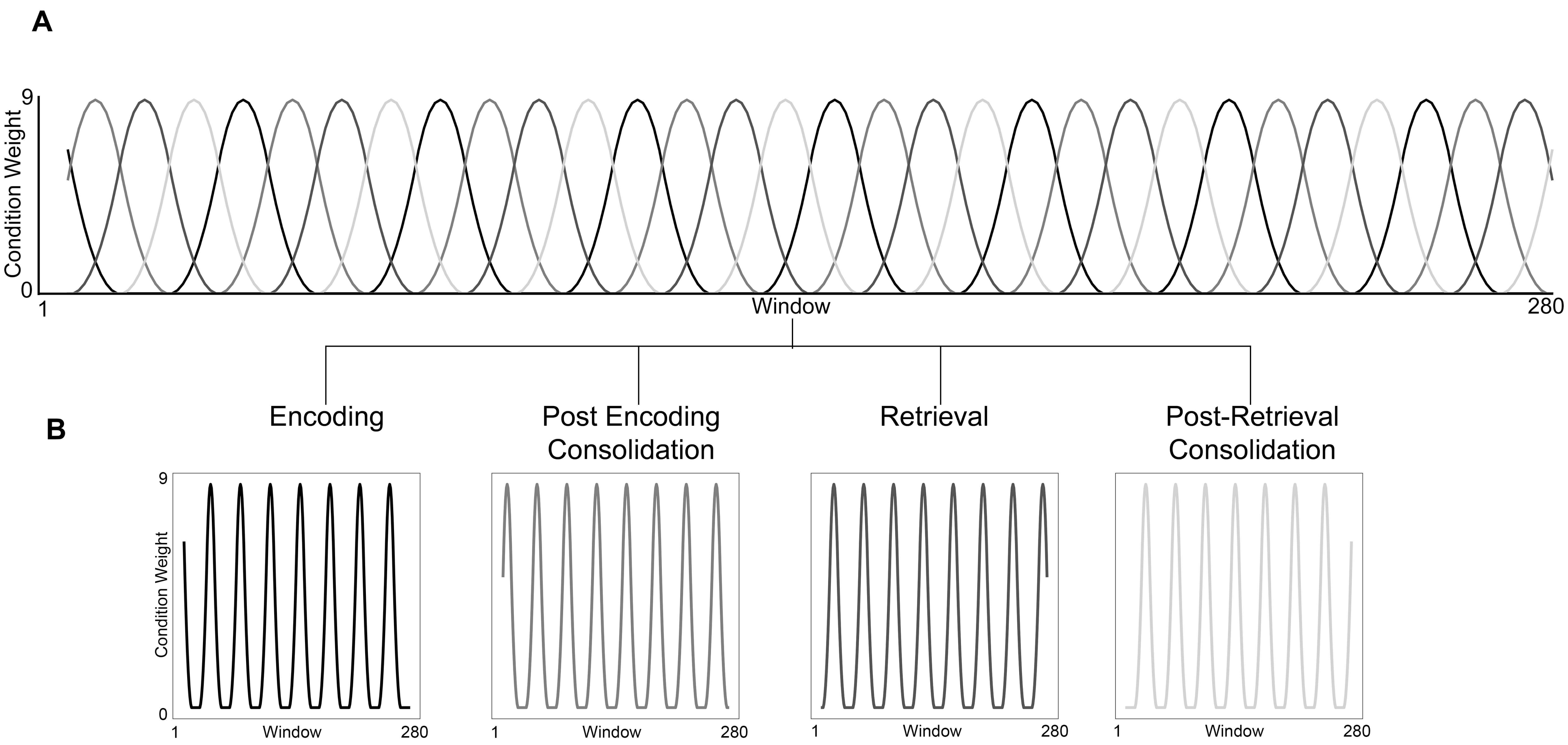


The task dynamics were represented by creating temporal functions for each of the four task conditions ($t_{Condition}$). In each function, we represent the degree to which that condition is represented at any point in time. This value ranged from 0 (i.e., there was no representation of that condition at that point in time) to 9 (i.e., at that point in time that was the only condition represented). The formed condition-related time series are depicted in Panel B and are superimposed on the same time line in Panel A.

Supplementary Figure 8

#### Encoding

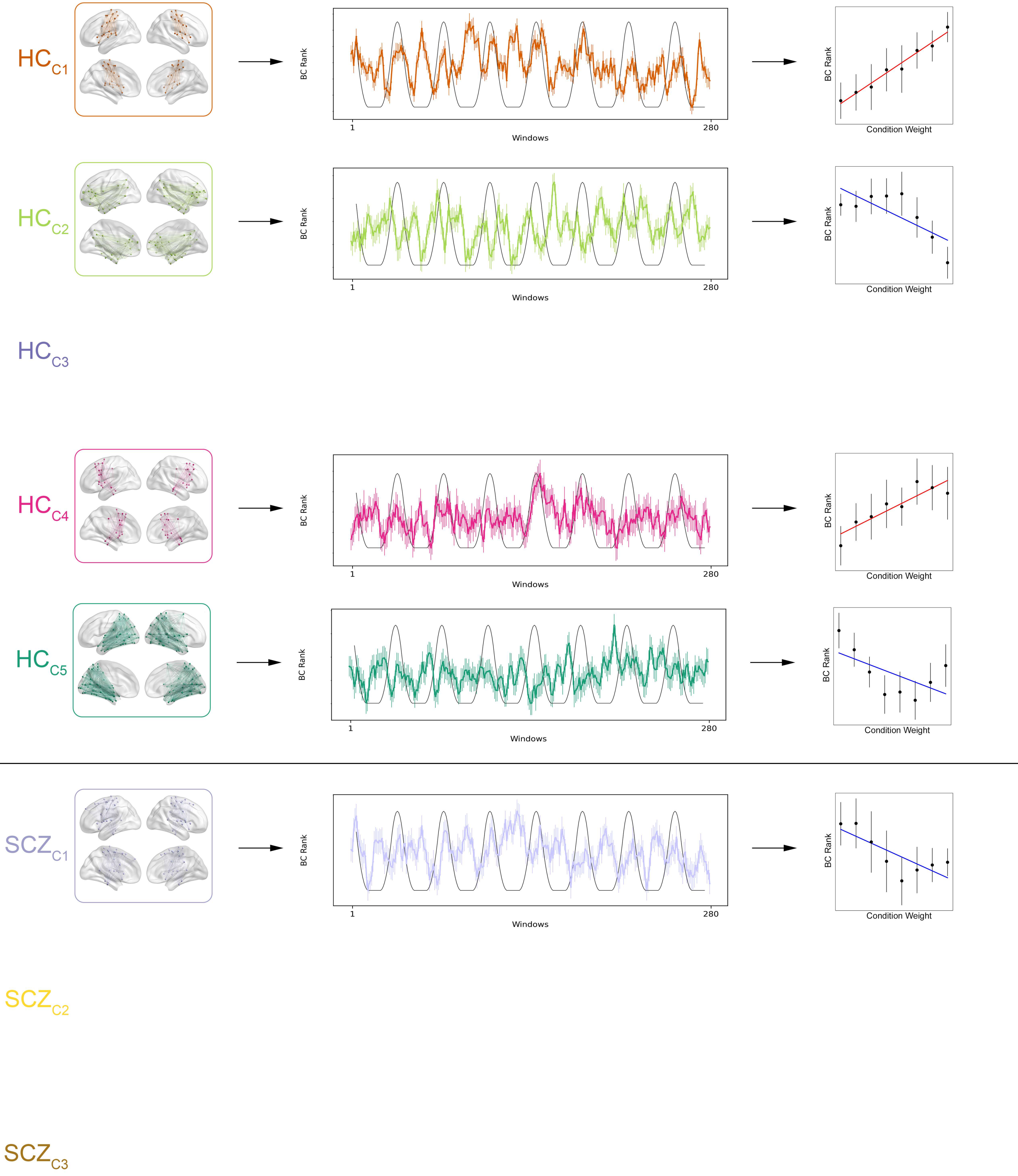


The figure explores the statistical relationships between the centrality dynamics of each cluster ($t_{Cluster}$) and the $t_{Condition}$ for Encoding. For each of $HC_{C1}$ – $HC_{C5}$ and $SCZ_{C1}$ – $SCZ_{C3}$, we overlay each $t_{Cluster}$ on each of $t_{Condition}$. Significant relationships ($p < 0.05$, Bonferroni corrected) between $t_{Cluster}$ and the $t_{Condition}$ for Encoding are depicted. The scatter plots (far right) depict average centrality measures in each of eight epochs (error bars are ± sem). The fitted function is the line of best linear fit. Across clusters, we observed multiple significant relationships between centrality dynamics and task conditions. This scheme of presenting effects is carried forward in Supplementary Figures 9 – 11.

### Post Encoding Consolidation

Supplementary Figure 9

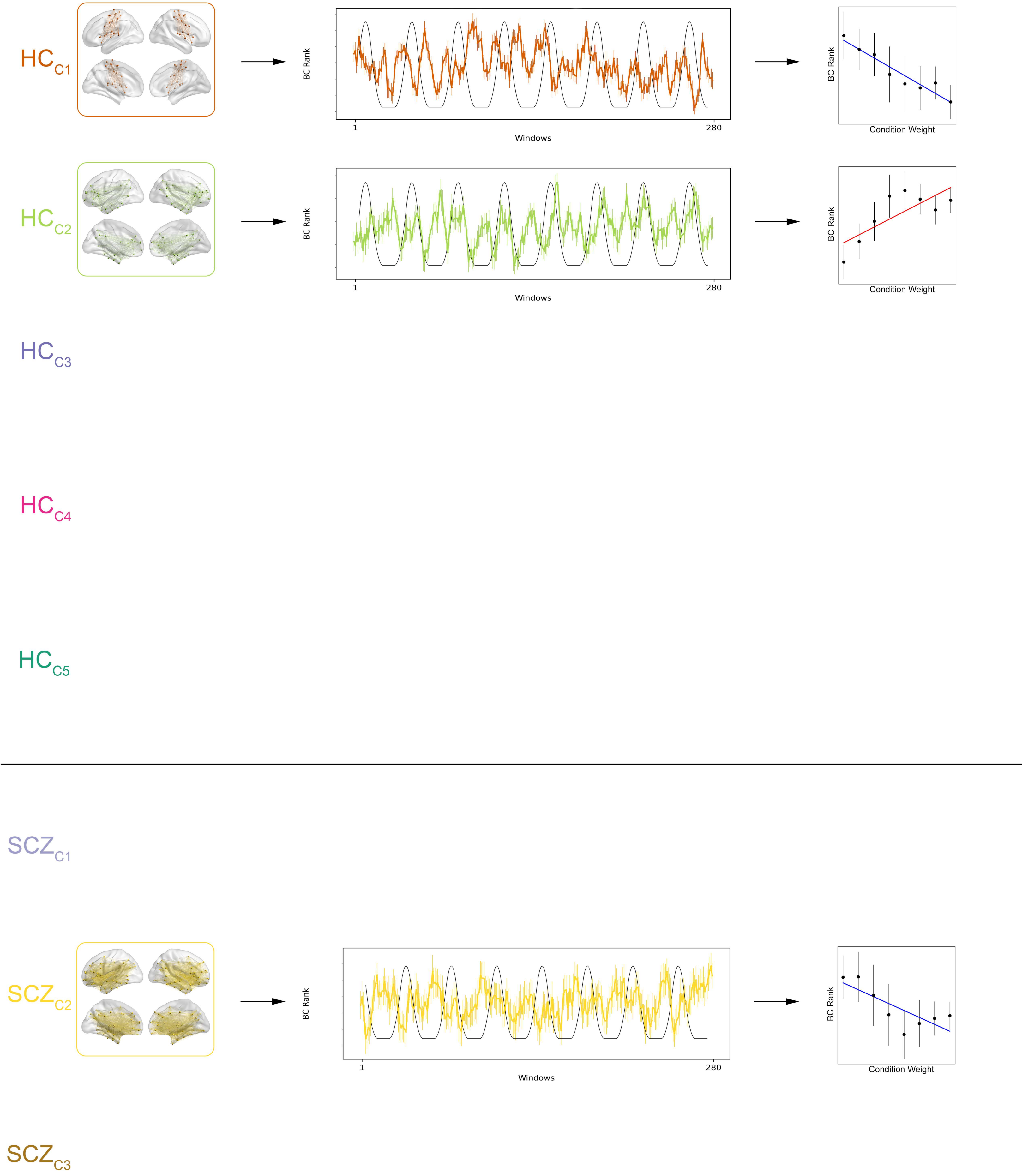


The figure explores the statistical relationships between the centrality dynamics of each cluster ($t_{Cluster}$) and the $t_{Condition}$ for Post-Encoding Consolidation.

Supplementary Figure 10

### Retrieval

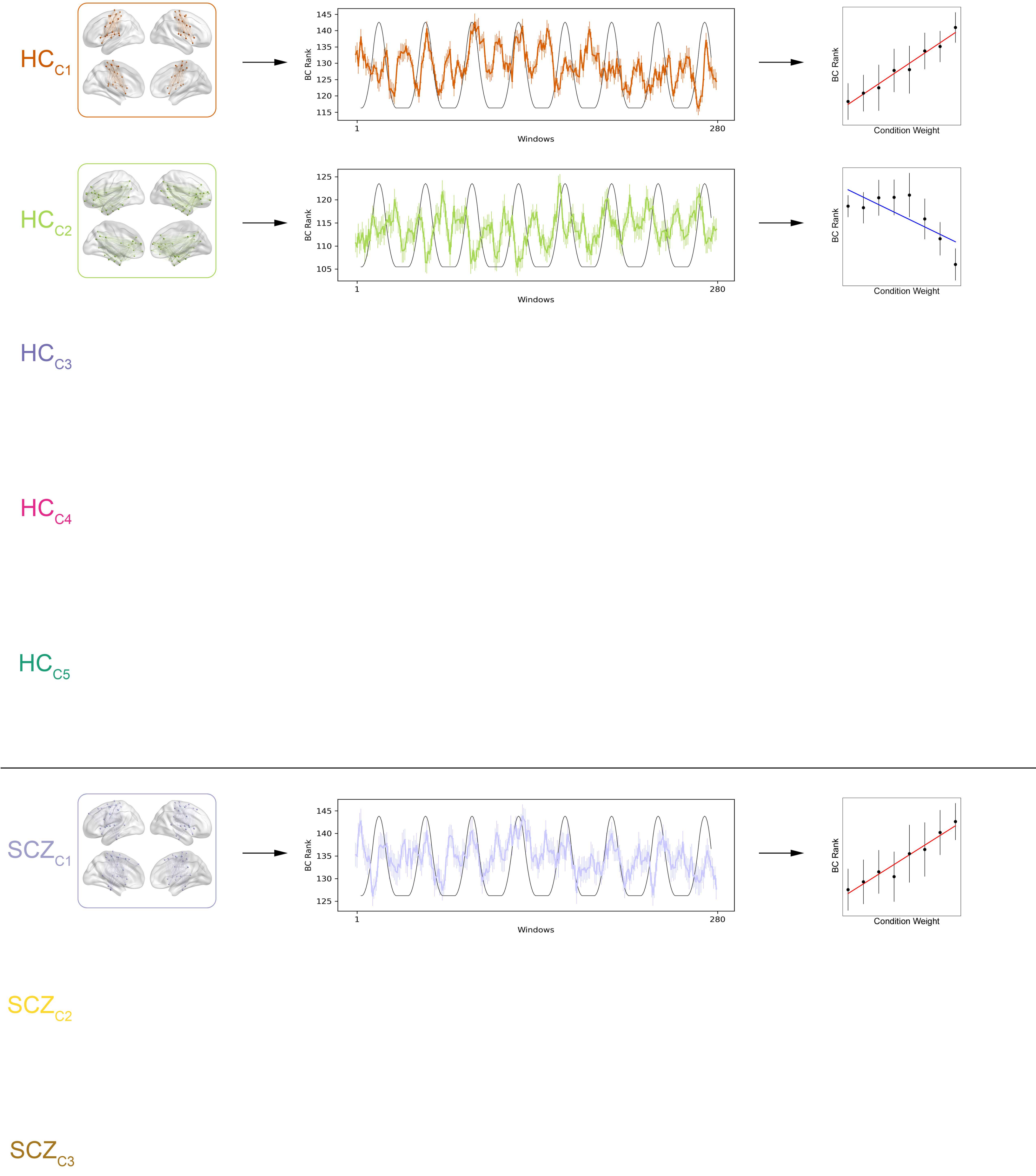


The figure explores the statistical relationships between the centrality dynamics of each cluster ($t_{Cluster}$) and the $t_{Condition}$ for Retrieval.

Supplementary Figure 11

### Post Retrieval Consolidation

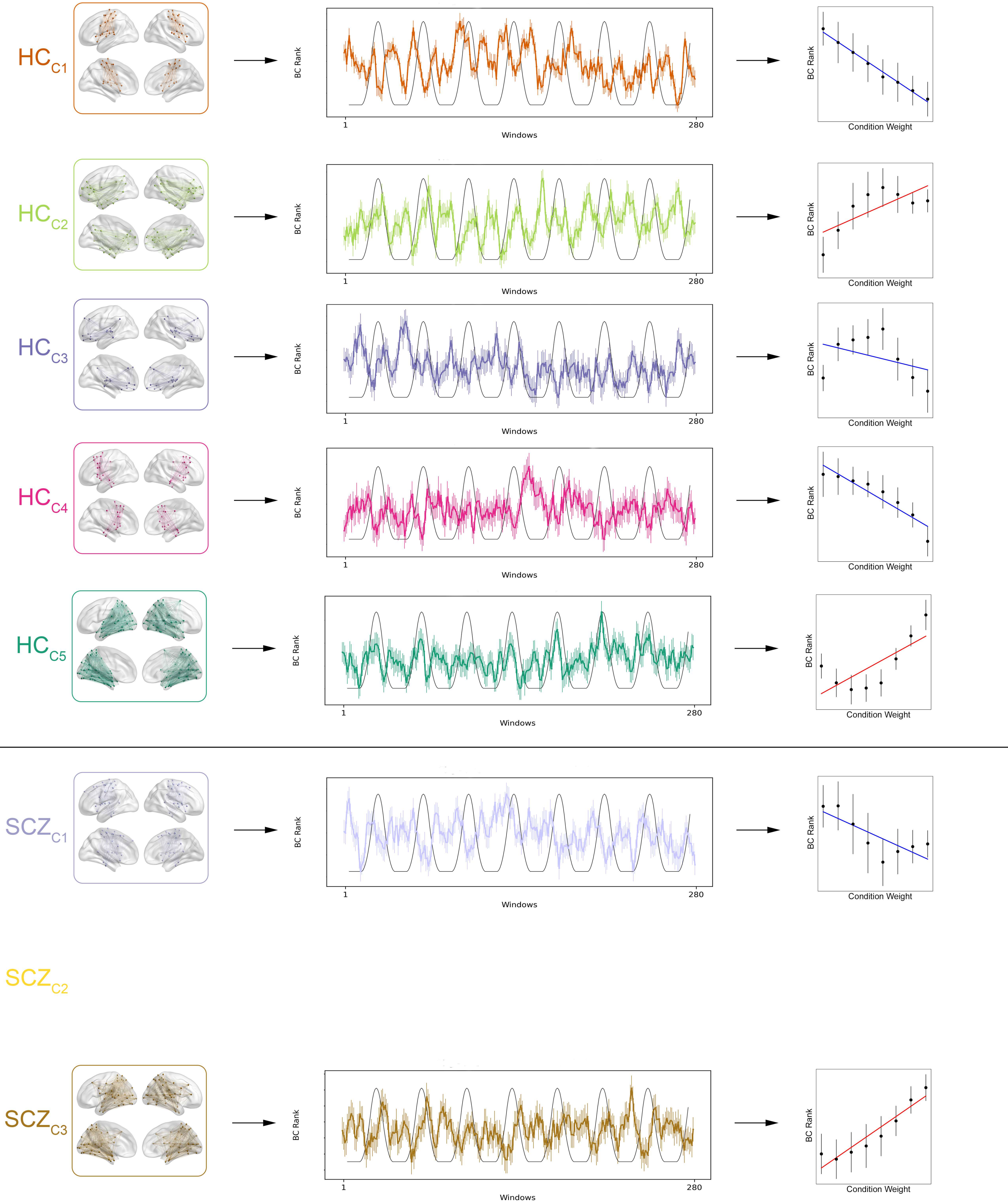


The figure explores the statistical relationships between the centrality dynamics of each cluster ($t_{Cluster}$) and the $t_{Condition}$ for Post-Retrieval Consolidation.

Supplementary Table 1. The table provides a detailed listing of the brain regions assigned to each cluster formed from applying AHC to centrality dynamics.

| Subnetwork | Brainnetome Region Number | Lobe | Gyrus | Anatomical and modified Cyto-architectonic descriptions |
|---|---|---|---|---|
| $HC_{C1}$ | 9 | Frontal Lobe | SFG, Superior Frontal Gyrus | A6m, medial area 6 |
| | 10 | Frontal Lobe | SFG, Superior Frontal Gyrus | A6m, medial area 6 |
| | 53 | Frontal Lobe | PrG, Precentral Gyrus | A4hf, area 4(head and face region) |
| | 54 | Frontal Lobe | PrG, Precentral Gyrus | A4hf, area 4(head and face region) |
| | 55 | Frontal Lobe | PrG, Precentral Gyrus | A6cdl, caudal dorsolateral area 6 |
| | 56 | Frontal Lobe | PrG, Precentral Gyrus | A6cdl, caudal dorsolateral area 6 |
| | 57 | Frontal Lobe | PrG, Precentral Gyrus | A4ul, area 4(upper limb region) |
| | 58 | Frontal Lobe | PrG, Precentral Gyrus | A4ul, area 4(upper limb region) |
| | 59 | Frontal Lobe | PrG, Precentral Gyrus | A4t, area 4(trunk region) |
| | 60 | Frontal Lobe | PrG, Precentral Gyrus | A4t, area 4(trunk region) |
| | 61 | Frontal Lobe | PrG, Precentral Gyrus | A4tl, area 4(tongue and larynx region) |
| | 62 | Frontal Lobe | PrG, Precentral Gyrus | A4tl, area 4(tongue and larynx region) |

| | | | |
|---|---|---|---|
| 63 | Frontal Lobe | PrG, Precentral Gyrus | A6cvl, caudal ventrolateral area 6 |
| 64 | Frontal Lobe | PrG, Precentral Gyrus | A6cvl, caudal ventrolateral area 6 |
| 65 | Frontal Lobe | PCL, Paracentral Lobule | A1/2/3ll, area1/2/3 (lower limb region) |
| 66 | Frontal Lobe | PCL, Paracentral Lobule | A1/2/3ll, area1/2/3 (lower limb region) |
| 67 | Frontal Lobe | PCL, Paracentral Lobule | A4ll, area 4, (lower limb region) |
| 68 | Frontal Lobe | PCL, Paracentral Lobule | A4ll, area 4, (lower limb region) |
| 71 | Temporal Lobe | STG, Superior Temporal Gyrus | A41/42, area 41/42 |
| 72 | Temporal Lobe | STG, Superior Temporal Gyrus | A41/42, area 41/42 |
| 73 | Temporal Lobe | STG, Superior Temporal Gyrus | TE1.0 and TE1.2 |
| 74 | Temporal Lobe | STG, Superior Temporal Gyrus | TE1.0 and TE1.2 |
| 75 | Temporal Lobe | STG, Superior Temporal Gyrus | A22c, caudal area 22 |
| 155 | Parietal Lobe | PoG, Postcentral Gyrus | A1/2/3ulhf, area 1/2/3(upper limb, head and face region) |
| 156 | Parietal Lobe | PoG, Postcentral Gyrus | A1/2/3ulhf, area 1/2/3(upper limb, head and face region) |

| | | | | |
|---|---|---|---|---|
| | 157 | Parietal Lobe | PoG, Postcentral Gyrus | A1/2/3tonIa, area 1/2/3(tongue and larynx region) |
| | 158 | Parietal Lobe | PoG, Postcentral Gyrus | A1/2/3tonIa, area 1/2/3(tongue and larynx region) |
| | 159 | Parietal Lobe | PoG, Postcentral Gyrus | A2, area 2 |
| | 160 | Parietal Lobe | PoG, Postcentral Gyrus | A2, area 2 |
| | 161 | Parietal Lobe | PoG, Postcentral Gyrus | A1/2/3tru, area1/2/3(trunk region) |
| | 162 | Parietal Lobe | PoG, Postcentral Gyrus | A1/2/3tru, area1/2/3(trunk region) |
| | 163 | Insular Lobe | INS, Insular Gyrus | G, hypergranular insula |
| | 164 | Insular Lobe | INS, Insular Gyrus | G, hypergranular insula |
| | 165 | Insular Lobe | INS, Insular Gyrus | vIa, ventral agranular insula |
| | 166 | Insular Lobe | INS, Insular Gyrus | vIa, ventral agranular insula |
| | 167 | Insular Lobe | INS, Insular Gyrus | dIa, dorsal agranular insula |
| | 168 | Insular Lobe | INS, Insular Gyrus | dIa, dorsal agranular insula |
| | 173 | Insular Lobe | INS, Insular Gyrus | dId, dorsal dysgranular insula |
| $HC_{C2}$ | 13 | Frontal Lobe | SFG, Superior Frontal Gyrus | A10m, medial area 10 |

| | | | |
|---|---|---|---|
| 14 | Frontal Lobe | SFG, Superior Frontal Gyrus | A10m, medial area 10 |
| 15 | Frontal Lobe | MFG, Middle Frontal Gyrus | A9/46d, dorsal area 9/46 |
| 16 | Frontal Lobe | MFG, Middle Frontal Gyrus | A9/46d, dorsal area 9/46 |
| 19 | Frontal Lobe | MFG, Middle Frontal Gyrus | A46, area 46 |
| 20 | Frontal Lobe | MFG, Middle Frontal Gyrus | A46, area 46 |
| 21 | Frontal Lobe | MFG, Middle Frontal Gyrus | A9/46v, ventral area 9/46 |
| 22 | Frontal Lobe | MFG, Middle Frontal Gyrus | A9/46v, ventral area 9/46 |
| 27 | Frontal Lobe | MFG, Middle Frontal Gyrus | A10l, lateral area10 |
| 28 | Frontal Lobe | MFG, Middle Frontal Gyrus | A10l, lateral area10 |
| 31 | Frontal Lobe | IFG, Inferior Frontal Gyrus | IFS, inferior frontal sulcus |
| 32 | Frontal Lobe | IFG, Inferior Frontal Gyrus | IFS, inferior frontal sulcus |
| 33 | Frontal Lobe | IFG, Inferior Frontal Gyrus | A45c, caudal area 45 |
| 34 | Frontal Lobe | IFG, Inferior Frontal Gyrus | A45c, caudal area 45 |
| 35 | Frontal Lobe | IFG, Inferior Frontal Gyrus | A45r, rostral area 45 |
| 36 | Frontal Lobe | IFG, Inferior Frontal Gyrus | A45r, rostral area 45 |
| 37 | Frontal Lobe | IFG, Inferior Frontal Gyrus | A44op, opercular area 44 |

| | | | |
|---|---|---|---|
| 38 | Frontal Lobe | IFG, Inferior Frontal Gyrus | A44op, opercular area 44 |
| 44 | Frontal Lobe | OrG, Orbital Gyrus | A12/47o, orbital area 12/47 |
| 45 | Frontal Lobe | OrG, Orbital Gyrus | A11l, lateral area 11 |
| 69 | Temporal Lobe | STG, Superior Temporal Gyrus | A38m, medial area 38 |
| 83 | Temporal Lobe | MTG, Middle Temporal Gyrus | A21r, rostral area 21 |
| 84 | Temporal Lobe | MTG, Middle Temporal Gyrus | A21r, rostral area 21 |
| 85 | Temporal Lobe | MTG, Middle Temporal Gyrus | A37dl, dorsolateral area37 |
| 86 | Temporal Lobe | MTG, Middle Temporal Gyrus | A37dl, dorsolateral area37 |
| 93 | Temporal Lobe | ITG, Inferior Temporal Gyrus | A20r, rostral area 20 |
| 94 | Temporal Lobe | ITG, Inferior Temporal Gyrus | A20r, rostral area 20 |
| 96 | Temporal Lobe | ITG, Inferior Temporal Gyrus | A20il, intermediate lateral area 20 |
| 99 | Temporal Lobe | ITG, Inferior Temporal Gyrus | A20cl, caudolateral of area 20 |
| 100 | Temporal Lobe | ITG, Inferior Temporal Gyrus | A20cl, caudolateral of area 20 |
| 101 | Temporal Lobe | ITG, Inferior Temporal Gyrus | A20cv, caudoventral of area 20 |

| | | | |
|---|---|---|---|
| 102 | Temporal Lobe | ITG, Inferior Temporal Gyrus | A20cv, caudoventral of area 20 |
| 103 | Temporal Lobe | FuG, Fusiform Gyrus | A20rv, rostroventral area 20 |
| 104 | Temporal Lobe | FuG, Fusiform Gyrus | A20rv, rostroventral area 20 |
| 136 | Parietal Lobe | IPL, Inferior Parietal Lobule | A39c, caudal area 39(PGp) |
| 137 | Parietal Lobe | IPL, Inferior Parietal Lobule | A39rd, rostrodorsal area 39(Hip3) |
| 138 | Parietal Lobe | IPL, Inferior Parietal Lobule | A39rd, rostrodorsal area 39(Hip3) |
| 143 | Parietal Lobe | IPL, Inferior Parietal Lobule | A39rv, rostroventral area 39(PGa) |
| 144 | Parietal Lobe | IPL, Inferior Parietal Lobule | A39rv, rostroventral area 39(PGa) |
| 153 | Parietal Lobe | Pcun, Precuneus | A31, area 31 (Lc1) |
| 154 | Parietal Lobe | Pcun, Precuneus | A31, area 31 (Lc1) |
| 175 | Limbic Lobe | CG, Cingulate Gyrus | A23d, dorsal area 23 |
| 176 | Limbic Lobe | CG, Cingulate Gyrus | A23d, dorsal area 23 |
| 179 | Limbic Lobe | CG, Cingulate Gyrus | A32p, pregenual area 32 |
| 180 | Limbic Lobe | CG, Cingulate Gyrus | A32p, pregenual area 32 |

| | | | | |
|---|---|---|---|---|
| | 188 | Limbic Lobe | CG, Cingulate Gyrus | A32sg, subgenual area 32 |
| | 212 | Subcortical Nuclei | Amyg, Amygdala | mAmyg, medial amygdala |
| | 214 | Subcortical Nuclei | Amyg, Amygdala | lAmyg, lateral amygdala |
| | 215 | Subcortical Nuclei | Hipp, Hippocampus | rHipp, rostral hippocampus |
| | 216 | Subcortical Nuclei | Hipp, Hippocampus | rHipp, rostral hippocampus |
| | 227 | Subcortical Nuclei | BG, Basal Ganglia | dCa, dorsal caudate |
| | 228 | Subcortical Nuclei | BG, Basal Ganglia | dCa, dorsal caudate |
| $HC_{C3}$ | 5 | Frontal Lobe | SFG, Superior Frontal Gyrus | A9l, lateral area 9 |
| | 6 | Frontal Lobe | SFG, Superior Frontal Gyrus | A9l, lateral area 9 |
| | 11 | Frontal Lobe | SFG, Superior Frontal Gyrus | A9m,medial area 9 |
| | 12 | Frontal Lobe | SFG, Superior Frontal Gyrus | A9m,medial area 9 |
| | 41 | Frontal Lobe | OrG, Orbital Gyrus | A14m, medial area 14 |
| | 42 | Frontal Lobe | OrG, Orbital Gyrus | A14m, medial area 14 |
| | 43 | Frontal Lobe | OrG, Orbital Gyrus | A12/47o, orbital area 12/47 |
| | 46 | Frontal Lobe | OrG, Orbital Gyrus | A11l, lateral area 11 |
| | 47 | Frontal Lobe | OrG, Orbital Gyrus | A11m, medial area 11 |

| | | | |
|---|---|---|---|
| 48 | Frontal Lobe | OrG, Orbital Gyrus | A11m, medial area 11 |
| 49 | Frontal Lobe | OrG, Orbital Gyrus | A13, area 13 |
| 50 | Frontal Lobe | OrG, Orbital Gyrus | A13, area 13 |
| 51 | Frontal Lobe | OrG, Orbital Gyrus | A12/47l, lateral area 12/47 |
| 52 | Frontal Lobe | OrG, Orbital Gyrus | A12/47l, lateral area 12/47 |
| 81 | Temporal Lobe | MTG, Middle Temporal Gyrus | A21c, caudal area 21 |
| 88 | Temporal Lobe | MTG, Middle Temporal Gyrus | aSTS, anterior superior temporal sulcus |
| 140 | Parietal Lobe | IPL, Inferior Parietal Lobule | A40rd, rostrodorsal area 40(PFt) |
| 141 | Parietal Lobe | IPL, Inferior Parietal Lobule | A40c, caudal area 40(PFm) |
| 142 | Parietal Lobe | IPL, Inferior Parietal Lobule | A40c, caudal area 40(PFm) |
| 187 | Limbic Lobe | CG, Cingulate Gyrus | A32sg, subgenual area 32 |
| 217 | Subcortical Nuclei | Hipp, Hippocampus | cHipp, caudal hippocampus |
| 230 | Subcortical Nuclei | BG, Basal Ganglia | dlPu, dorsolateral putamen |
| 231 | Subcortical Nuclei | Tha, Thalamus | mPFtha, medial pre-frontal thalamus |
| 232 | Subcortical Nuclei | Tha, Thalamus | mPFtha, medial pre-frontal thalamus |

| | | | | |
|---|---|---|---|---|
| | 233 | Subcortical Nuclei | Tha, Thalamus | mPMtha, pre-motor thalamus |
| | 234 | Subcortical Nuclei | Tha, Thalamus | mPMtha, pre-motor thalamus |
| | 235 | Subcortical Nuclei | Tha, Thalamus | Stha, sensory thalamus |
| | 236 | Subcortical Nuclei | Tha, Thalamus | Stha, sensory thalamus |
| | 237 | Subcortical Nuclei | Tha, Thalamus | rTtha, rostral temporal thalamus |
| | 238 | Subcortical Nuclei | Tha, Thalamus | rTtha, rostral temporal thalamus |
| | 239 | Subcortical Nuclei | Tha, Thalamus | PPtha, posterior parietal thalamus |
| | 240 | Subcortical Nuclei | Tha, Thalamus | PPtha, posterior parietal thalamus |
| | 241 | Subcortical Nuclei | Tha, Thalamus | Otha, occipital thalamus |
| | 242 | Subcortical Nuclei | Tha, Thalamus | Otha, occipital thalamus |
| | 243 | Subcortical Nuclei | Tha, Thalamus | cTtha, caudal temporal thalamus |
| | 244 | Subcortical Nuclei | Tha, Thalamus | cTtha, caudal temporal thalamus |
| | 246 | Subcortical Nuclei | Tha, Thalamus | lPFtha, lateral pre-frontal thalamus |
| $HC_{C4}$ | 1 | Frontal Lobe | SFG, Superior Frontal Gyrus | A8m, medial area 8 |
| | 2 | Frontal Lobe | SFG, Superior Frontal Gyrus | A8m, medial area 8 |
| | 3 | Frontal Lobe | SFG, Superior Frontal Gyrus | A8dl, dorsolateral area 8 |

| | | | |
|---|---|---|---|
| 4 | Frontal Lobe | SFG, Superior Frontal Gyrus | A8dl, dorsolateral area 8 |
| 7 | Frontal Lobe | SFG, Superior Frontal Gyrus | A6dl, dorsolateral area 6 |
| 17 | Frontal Lobe | MFG, Middle Frontal Gyrus | IFJ, inferior frontal junction |
| 18 | Frontal Lobe | MFG, Middle Frontal Gyrus | IFJ, inferior frontal junction |
| 23 | Frontal Lobe | MFG, Middle Frontal Gyrus | A8vl, ventrolateral area 8 |
| 24 | Frontal Lobe | MFG, Middle Frontal Gyrus | A8vl, ventrolateral area 8 |
| 25 | Frontal Lobe | MFG, Middle Frontal Gyrus | A6vl, ventrolateral area 6 |
| 26 | Frontal Lobe | MFG, Middle Frontal Gyrus | A6vl, ventrolateral area 6 |
| 29 | Frontal Lobe | IFG, Inferior Frontal Gyrus | A44d,dorsal area 44 |
| 30 | Frontal Lobe | IFG, Inferior Frontal Gyrus | A44d,dorsal area 44 |
| 39 | Frontal Lobe | IFG, Inferior Frontal Gyrus | A44v, ventral area 44 |
| 82 | Temporal Lobe | MTG, Middle Temporal Gyrus | A21c, caudal area 21 |
| 87 | Temporal Lobe | MTG, Middle Temporal Gyrus | aSTS, anterior superior temporal sulcus |
| 89 | Temporal Lobe | ITG, Inferior Temporal Gyrus | A20iv, intermediate ventral area 20 |

| | | | |
|---|---|---|---|
| 112 | Temporal Lobe | PhG, Parahippocampal Gyrus | A35/36c, caudal area 35/36 |
| 113 | Temporal Lobe | PhG, Parahippocampal Gyrus | TL, area TL (lateral PPHC, posterior parahippocampal gyrus) |
| 114 | Temporal Lobe | PhG, Parahippocampal Gyrus | TL, area TL (lateral PPHC, posterior parahippocampal gyrus) |
| 177 | Limbic Lobe | CG, Cingulate Gyrus | A24rv, rostroventral area 24 |
| 178 | Limbic Lobe | CG, Cingulate Gyrus | A24rv, rostroventral area 24 |
| 183 | Limbic Lobe | CG, Cingulate Gyrus | A24cd, caudodorsal area 24 |
| 184 | Limbic Lobe | CG, Cingulate Gyrus | A24cd, caudodorsal area 24 |
| 185 | Limbic Lobe | CG, Cingulate Gyrus | A23c, caudal area 23 |
| 186 | Limbic Lobe | CG, Cingulate Gyrus | A23c, caudal area 23 |
| 218 | Subcortical Nuclei | Hipp, Hippocampus | cHipp, caudal hippocampus |
| 219 | Subcortical Nuclei | BG, Basal Ganglia | vCa, ventral caudate |
| 220 | Subcortical Nuclei | BG, Basal Ganglia | vCa, ventral caudate |
| 221 | Subcortical Nuclei | BG, Basal Ganglia | GP, globus pallidus |

| | | | | |
|---|---|---|---|---|
| | 222 | Subcortical Nuclei | BG, Basal Ganglia | GP, globus pallidus |
| | 223 | Subcortical Nuclei | BG, Basal Ganglia | NAC, nucleus accumbens |
| | 224 | Subcortical Nuclei | BG, Basal Ganglia | NAC, nucleus accumbens |
| | 225 | Subcortical Nuclei | BG, Basal Ganglia | vmPu, ventromedial putamen |
| | 226 | Subcortical Nuclei | BG, Basal Ganglia | vmPu, ventromedial putamen |
| $HC_{C5}$ | 8 | Frontal Lobe | SFG, Superior Frontal Gyrus | A6dl, dorsolateral area 6 |
| | 40 | Frontal Lobe | IFG, Inferior Frontal Gyrus | A44v, ventral area 44 |
| | 70 | Temporal Lobe | STG, Superior Temporal Gyrus | A38m, medial area 38 |
| | 76 | Temporal Lobe | STG, Superior Temporal Gyrus | A22c, caudal area 22 |
| | 77 | Temporal Lobe | STG, Superior Temporal Gyrus | A38l, lateral area 38 |
| | 78 | Temporal Lobe | STG, Superior Temporal Gyrus | A38l, lateral area 38 |
| | 79 | Temporal Lobe | STG, Superior Temporal Gyrus | A22r, rostral area 22 |
| | 80 | Temporal Lobe | STG, Superior Temporal Gyrus | A22r, rostral area 22 |
| | 90 | Temporal Lobe | ITG, Inferior Temporal Gyrus | A20iv, intermediate ventral area 20 |
| | 91 | Temporal Lobe | ITG, Inferior Temporal Gyrus | A37elv, extreme lateroventral area37 |

| | | | |
|---|---|---|---|
| 92 | Temporal Lobe | ITG, Inferior Temporal Gyrus | A37elv, extreme lateroventral area37 |
| 95 | Temporal Lobe | ITG, Inferior Temporal Gyrus | A20il, intermediate lateral area 20 |
| 97 | Temporal Lobe | ITG, Inferior Temporal Gyrus | A37vl, ventrolateral area 37 |
| 98 | Temporal Lobe | ITG, Inferior Temporal Gyrus | A37vl, ventrolateral area 37 |
| 105 | Temporal Lobe | FuG, Fusiform Gyrus | A37mv, medioventral area37 |
| 106 | Temporal Lobe | FuG, Fusiform Gyrus | A37mv, medioventral area37 |
| 107 | Temporal Lobe | FuG, Fusiform Gyrus | A37lv, lateroventral area37 |
| 108 | Temporal Lobe | FuG, Fusiform Gyrus | A37lv, lateroventral area37 |
| 109 | Temporal Lobe | PhG, Parahippocampal Gyrus | A35/36r, rostral area 35/36 |
| 110 | Temporal Lobe | PhG, Parahippocampal Gyrus | A35/36r, rostral area 35/36 |
| 111 | Temporal Lobe | PhG, Parahippocampal Gyrus | A35/36c, caudal area 35/36 |
| 115 | Temporal Lobe | PhG, Parahippocampal Gyrus | A28/34, area 28/34 (EC, entorhinal cortex) |

| | | | |
|---|---|---|---|
| 116 | Temporal Lobe | PhG, Parahippocampal Gyrus | A28/34, area 28/34 (EC, entorhinal cortex) |
| 117 | Temporal Lobe | PhG, Parahippocampal Gyrus | TI, area TI(temporal agranular insular cortex) |
| 118 | Temporal Lobe | PhG, Parahippocampal Gyrus | TI, area TI(temporal agranular insular cortex) |
| 119 | Temporal Lobe | PhG, Parahippocampal Gyrus | TH, area TH (medial PPHC) |
| 120 | Temporal Lobe | PhG, Parahippocampal Gyrus | TH, area TH (medial PPHC) |
| 121 | Temporal Lobe | pSTS, posterior Superior Temporal Sulcus | rpSTS, rostroposterior superior temporal sulcus |
| 122 | Temporal Lobe | pSTS, posterior Superior Temporal Sulcus | rpSTS, rostroposterior superior temporal sulcus |
| 123 | Temporal Lobe | pSTS, posterior Superior Temporal Sulcus | cpSTS, caudoposterior superior temporal sulcus |
| 124 | Temporal Lobe | pSTS, posterior Superior Temporal Sulcus | cpSTS, caudoposterior superior temporal sulcus |
| 125 | Parietal Lobe | SPL, Superior Parietal Lobule | A7r, rostral area 7 |
| 126 | Parietal Lobe | SPL, Superior Parietal Lobule | A7r, rostral area 7 |

| | | | |
|---|---|---|---|
| 127 | Parietal Lobe | SPL, Superior Parietal Lobule | A7c, caudal area 7 |
| 128 | Parietal Lobe | SPL, Superior Parietal Lobule | A7c, caudal area 7 |
| 129 | Parietal Lobe | SPL, Superior Parietal Lobule | A5l, lateral area 5 |
| 130 | Parietal Lobe | SPL, Superior Parietal Lobule | A5l, lateral area 5 |
| 131 | Parietal Lobe | SPL, Superior Parietal Lobule | A7pc, postcentral area 7 |
| 132 | Parietal Lobe | SPL, Superior Parietal Lobule | A7pc, postcentral area 7 |
| 133 | Parietal Lobe | SPL, Superior Parietal Lobule | A7ip, intraparietal area 7(hIP3) |
| 134 | Parietal Lobe | SPL, Superior Parietal Lobule | A7ip, intraparietal area 7(hIP3) |
| 135 | Parietal Lobe | IPL, Inferior Parietal Lobule | A39c, caudal area 39(PGp) |
| 139 | Parietal Lobe | IPL, Inferior Parietal Lobule | A40rd, rostrodorsal area 40(PFt) |
| 145 | Parietal Lobe | IPL, Inferior Parietal Lobule | A40rv, rostroventral area 40(PFop) |
| 146 | Parietal Lobe | IPL, Inferior Parietal Lobule | A40rv, rostroventral area 40(PFop) |
| 147 | Parietal Lobe | Pcun, Precuneus | A7m, medial area 7(PEp) |
| 148 | Parietal Lobe | Pcun, Precuneus | A7m, medial area 7(PEp) |
| 149 | Parietal Lobe | Pcun, Precuneus | A5m, medial area 5(PEm) |

| | | | |
|---|---|---|---|
| 150 | Parietal Lobe | Pcun, Precuneus | A5m, medial area 5(PEm) |
| 151 | Parietal Lobe | Pcun, Precuneus | dmPOS, dorsomedial parietooccipital sulcus(PEr) |
| 152 | Parietal Lobe | Pcun, Precuneus | dmPOS, dorsomedial parietooccipital sulcus(PEr) |
| 169 | Insular Lobe | INS, Insular Gyrus | vId/vIg, ventral dysgranular and granular insula |
| 170 | Insular Lobe | INS, Insular Gyrus | vId/vIg, ventral dysgranular and granular insula |
| 171 | Insular Lobe | INS, Insular Gyrus | dIg, dorsal granular insula |
| 172 | Insular Lobe | INS, Insular Gyrus | dIg, dorsal granular insula |
| 174 | Insular Lobe | INS, Insular Gyrus | dId, dorsal dysgranular insula |
| 181 | Limbic Lobe | CG, Cingulate Gyrus | A23v, ventral area 23 |
| 182 | Limbic Lobe | CG, Cingulate Gyrus | A23v, ventral area 23 |
| 189 | Occipital Lobe | MVOcC, MedioVentral Occipital Cortex | cLinG, caudal lingual gyrus |
| 190 | Occipital Lobe | MVOcC, MedioVentral Occipital Cortex | cLinG, caudal lingual gyrus |
| 191 | Occipital Lobe | MVOcC, MedioVentral Occipital Cortex | rCunG, rostral cuneus gyrus |

| | | | |
|---|---|---|---|
| 192 | Occipital Lobe | MVOcC, MedioVentral Occipital Cortex | rCunG, rostral cuneus gyrus |
| 193 | Occipital Lobe | MVOcC, MedioVentral Occipital Cortex | cCunG, caudal cuneus gyrus |
| 194 | Occipital Lobe | MVOcC, MedioVentral Occipital Cortex | cCunG, caudal cuneus gyrus |
| 195 | Occipital Lobe | MVOcC, MedioVentral Occipital Cortex | rLinG, rostral lingual gyrus |
| 196 | Occipital Lobe | MVOcC, MedioVentral Occipital Cortex | rLinG, rostral lingual gyrus |
| 197 | Occipital Lobe | MVOcC, MedioVentral Occipital Cortex | vmPOS,ventromedial parietooccipital sulcus |
| 198 | Occipital Lobe | MVOcC, MedioVentral Occipital Cortex | vmPOS,ventromedial parietooccipital sulcus |
| 199 | Occipital Lobe | LOcC, lateral Occipital Cortex | mOccG, middle occipital gyrus |
| 200 | Occipital Lobe | LOcC, lateral Occipital Cortex | mOccG, middle occipital gyrus |
| 201 | Occipital Lobe | LOcC, lateral Occipital Cortex | V5/MT+, area V5/MT+ |

| | | | |
|---|---|---|---|
| 202 | Occipital Lobe | LOcC, lateral Occipital Cortex | V5/MT+, area V5/MT+ |
| 203 | Occipital Lobe | LOcC, lateral Occipital Cortex | OPC, occipital polar cortex |
| 204 | Occipital Lobe | LOcC, lateral Occipital Cortex | OPC, occipital polar cortex |
| 205 | Occipital Lobe | LOcC, lateral Occipital Cortex | iOccG, inferior occipital gyrus |
| 206 | Occipital Lobe | LOcC, lateral Occipital Cortex | iOccG, inferior occipital gyrus |
| 207 | Occipital Lobe | LOcC, lateral Occipital Cortex | msOccG, medial superior occipital gyrus |
| 208 | Occipital Lobe | LOcC, lateral Occipital Cortex | msOccG, medial superior occipital gyrus |
| 209 | Occipital Lobe | LOcC, lateral Occipital Cortex | lsOccG, lateral superior occipital gyrus |
| 210 | Occipital Lobe | LOcC, lateral Occipital Cortex | lsOccG, lateral superior occipital gyrus |
| 211 | Subcortical Nuclei | Amyg, Amygdala | mAmyg, medial amygdala |
| 213 | Subcortical Nuclei | Amyg, Amygdala | lAmyg, lateral amygdala |
| 229 | Subcortical Nuclei | BG, Basal Ganglia | dlPu, dorsolateral putamen |
| 245 | Subcortical Nuclei | Tha, Thalamus | lPFtha, lateral pre-frontal thalamus |

| | | | | |
|---|---|---|---|---|
| $SCZ_{C1}$ | 1 | Frontal Lobe | SFG, Superior Frontal Gyrus | A8m, medial area 8 |
| | 2 | Frontal Lobe | SFG, Superior Frontal Gyrus | A8m, medial area 8 |
| | 3 | Frontal Lobe | SFG, Superior Frontal Gyrus | A8dl, dorsolateral area 8 |
| | 4 | Frontal Lobe | SFG, Superior Frontal Gyrus | A8dl, dorsolateral area 8 |
| | 5 | Frontal Lobe | SFG, Superior Frontal Gyrus | A9l, lateral area 9 |
| | 7 | Frontal Lobe | SFG, Superior Frontal Gyrus | A6dl, dorsolateral area 6 |
| | 8 | Frontal Lobe | SFG, Superior Frontal Gyrus | A6dl, dorsolateral area 6 |
| | 9 | Frontal Lobe | SFG, Superior Frontal Gyrus | A6m, medial area 6 |
| | 10 | Frontal Lobe | SFG, Superior Frontal Gyrus | A6m, medial area 6 |
| | 53 | Frontal Lobe | PrG, Precentral Gyrus | A4hf, area 4(head and face region) |
| | 54 | Frontal Lobe | PrG, Precentral Gyrus | A4hf, area 4(head and face region) |
| | 55 | Frontal Lobe | PrG, Precentral Gyrus | A6cdl, caudal dorsolateral area 6 |
| | 56 | Frontal Lobe | PrG, Precentral Gyrus | A6cdl, caudal dorsolateral area 6 |
| | 57 | Frontal Lobe | PrG, Precentral Gyrus | A4ul, area 4(upper limb region) |
| | 58 | Frontal Lobe | PrG, Precentral Gyrus | A4ul, area 4(upper limb region) |
| | 59 | Frontal Lobe | PrG, Precentral Gyrus | A4t, area 4(trunk region) |

| | | | |
|---|---|---|---|
| 60 | Frontal Lobe | PrG, Precentral Gyrus | A4t, area 4(trunk region) |
| 61 | Frontal Lobe | PrG, Precentral Gyrus | A4tl, area 4(tongue and larynx region) |
| 62 | Frontal Lobe | PrG, Precentral Gyrus | A4tl, area 4(tongue and larynx region) |
| 65 | Frontal Lobe | PCL, Paracentral Lobule | A1/2/3ll, area1/2/3 (lower limb region) |
| 66 | Frontal Lobe | PCL, Paracentral Lobule | A1/2/3ll, area1/2/3 (lower limb region) |
| 67 | Frontal Lobe | PCL, Paracentral Lobule | A4ll, area 4, (lower limb region) |
| 68 | Frontal Lobe | PCL, Paracentral Lobule | A4ll, area 4, (lower limb region) |
| 74 | Temporal Lobe | STG, Superior Temporal Gyrus | TE1.0 and TE1.2 |
| 89 | Temporal Lobe | ITG, Inferior Temporal Gyrus | A20iv, intermediate ventral area 20 |
| 111 | Temporal Lobe | PhG, Parahippocampal Gyrus | A35/36c, caudal area 35/36 |
| 112 | Temporal Lobe | PhG, Parahippocampal Gyrus | A35/36c, caudal area 35/36 |
| 113 | Temporal Lobe | PhG, Parahippocampal Gyrus | TL, area TL (lateral PPHC, posterior parahippocampal gyrus) |

| | | | |
|---|---|---|---|
| 114 | Temporal Lobe | PhG, Parahippocampal Gyrus | TL, area TL (lateral PPHC, posterior parahippocampal gyrus) |
| 159 | Parietal Lobe | PoG, Postcentral Gyrus | A2, area 2 |
| 160 | Parietal Lobe | PoG, Postcentral Gyrus | A2, area 2 |
| 161 | Parietal Lobe | PoG, Postcentral Gyrus | A1/2/3tru, area1/2/3(trunk region) |
| 162 | Parietal Lobe | PoG, Postcentral Gyrus | A1/2/3tru, area1/2/3(trunk region) |
| 163 | Insular Lobe | INS, Insular Gyrus | G, hypergranular insula |
| 164 | Insular Lobe | INS, Insular Gyrus | G, hypergranular insula |
| 165 | Insular Lobe | INS, Insular Gyrus | vIa, ventral agranular insula |
| 166 | Insular Lobe | INS, Insular Gyrus | vIa, ventral agranular insula |
| 167 | Insular Lobe | INS, Insular Gyrus | dIa, dorsal agranular insula |
| 168 | Insular Lobe | INS, Insular Gyrus | dIa, dorsal agranular insula |
| 171 | Insular Lobe | INS, Insular Gyrus | dIg, dorsal granular insula |
| 172 | Insular Lobe | INS, Insular Gyrus | dIg, dorsal granular insula |
| 173 | Insular Lobe | INS, Insular Gyrus | dId, dorsal dysgranular insula |

| | | | | |
|---|---|---|---|---|
| | 174 | Insular Lobe | INS, Insular Gyrus | dId, dorsal dysgranular insula |
| | 176 | Limbic Lobe | CG, Cingulate Gyrus | A23d, dorsal area 23 |
| | 177 | Limbic Lobe | CG, Cingulate Gyrus | A24rv, rostroventral area 24 |
| | 181 | Limbic Lobe | CG, Cingulate Gyrus | A23v, ventral area 23 |
| | 182 | Limbic Lobe | CG, Cingulate Gyrus | A23v, ventral area 23 |
| $SCZ_{C2}$ | 6 | Frontal Lobe | SFG, Superior Frontal Gyrus | A9l, lateral area 9 |
| | 11 | Frontal Lobe | SFG, Superior Frontal Gyrus | A9m,medial area 9 |
| | 12 | Frontal Lobe | SFG, Superior Frontal Gyrus | A9m,medial area 9 |
| | 13 | Frontal Lobe | SFG, Superior Frontal Gyrus | A10m, medial area 10 |
| | 14 | Frontal Lobe | SFG, Superior Frontal Gyrus | A10m, medial area 10 |
| | 15 | Frontal Lobe | MFG, Middle Frontal Gyrus | A9/46d, dorsal area 9/46 |
| | 16 | Frontal Lobe | MFG, Middle Frontal Gyrus | A9/46d, dorsal area 9/46 |
| | 17 | Frontal Lobe | MFG, Middle Frontal Gyrus | IFJ, inferior frontal junction |
| | 18 | Frontal Lobe | MFG, Middle Frontal Gyrus | IFJ, inferior frontal junction |
| | 19 | Frontal Lobe | MFG, Middle Frontal Gyrus | A46, area 46 |
| | 20 | Frontal Lobe | MFG, Middle Frontal Gyrus | A46, area 46 |

| | | | |
|---|---|---|---|
| 21 | Frontal Lobe | MFG, Middle Frontal Gyrus | A9/46v, ventral area 9/46 |
| 22 | Frontal Lobe | MFG, Middle Frontal Gyrus | A9/46v, ventral area 9/46 |
| 25 | Frontal Lobe | MFG, Middle Frontal Gyrus | A6vl, ventrolateral area 6 |
| 26 | Frontal Lobe | MFG, Middle Frontal Gyrus | A6vl, ventrolateral area 6 |
| 27 | Frontal Lobe | MFG, Middle Frontal Gyrus | A10l, lateral area10 |
| 28 | Frontal Lobe | MFG, Middle Frontal Gyrus | A10l, lateral area10 |
| 29 | Frontal Lobe | IFG, Inferior Frontal Gyrus | A44d,dorsal area 44 |
| 30 | Frontal Lobe | IFG, Inferior Frontal Gyrus | A44d,dorsal area 44 |
| 31 | Frontal Lobe | IFG, Inferior Frontal Gyrus | IFS, inferior frontal sulcus |
| 33 | Frontal Lobe | IFG, Inferior Frontal Gyrus | A45c, caudal area 45 |
| 34 | Frontal Lobe | IFG, Inferior Frontal Gyrus | A45c, caudal area 45 |
| 35 | Frontal Lobe | IFG, Inferior Frontal Gyrus | A45r, rostral area 45 |
| 36 | Frontal Lobe | IFG, Inferior Frontal Gyrus | A45r, rostral area 45 |
| 37 | Frontal Lobe | IFG, Inferior Frontal Gyrus | A44op, opercular area 44 |
| 38 | Frontal Lobe | IFG, Inferior Frontal Gyrus | A44op, opercular area 44 |
| 39 | Frontal Lobe | IFG, Inferior Frontal Gyrus | A44v, ventral area 44 |

| | | | |
|---|---|---|---|
| 40 | Frontal Lobe | IFG, Inferior Frontal Gyrus | A44v, ventral area 44 |
| 41 | Frontal Lobe | OrG, Orbital Gyrus | A14m, medial area 14 |
| 42 | Frontal Lobe | OrG, Orbital Gyrus | A14m, medial area 14 |
| 43 | Frontal Lobe | OrG, Orbital Gyrus | A12/47o, orbital area 12/47 |
| 44 | Frontal Lobe | OrG, Orbital Gyrus | A12/47o, orbital area 12/47 |
| 45 | Frontal Lobe | OrG, Orbital Gyrus | A11l, lateral area 11 |
| 46 | Frontal Lobe | OrG, Orbital Gyrus | A11l, lateral area 11 |
| 47 | Frontal Lobe | OrG, Orbital Gyrus | A11m, medial area 11 |
| 48 | Frontal Lobe | OrG, Orbital Gyrus | A11m, medial area 11 |
| 49 | Frontal Lobe | OrG, Orbital Gyrus | A13, area 13 |
| 50 | Frontal Lobe | OrG, Orbital Gyrus | A13, area 13 |
| 51 | Frontal Lobe | OrG, Orbital Gyrus | A12/47l, lateral area 12/47 |
| 52 | Frontal Lobe | OrG, Orbital Gyrus | A12/47l, lateral area 12/47 |
| 69 | Temporal Lobe | STG, Superior Temporal Gyrus | A38m, medial area 38 |
| 70 | Temporal Lobe | STG, Superior Temporal Gyrus | A38m, medial area 38 |
| 77 | Temporal Lobe | STG, Superior Temporal Gyrus | A38l, lateral area 38 |

| | | | |
|---|---|---|---|
| 78 | Temporal Lobe | STG, Superior Temporal Gyrus | A38l, lateral area 38 |
| 79 | Temporal Lobe | STG, Superior Temporal Gyrus | A22r, rostral area 22 |
| 80 | Temporal Lobe | STG, Superior Temporal Gyrus | A22r, rostral area 22 |
| 81 | Temporal Lobe | MTG, Middle Temporal Gyrus | A21c, caudal area 21 |
| 82 | Temporal Lobe | MTG, Middle Temporal Gyrus | A21c, caudal area 21 |
| 83 | Temporal Lobe | MTG, Middle Temporal Gyrus | A21r, rostral area 21 |
| 84 | Temporal Lobe | MTG, Middle Temporal Gyrus | A21r, rostral area 21 |
| 85 | Temporal Lobe | MTG, Middle Temporal Gyrus | A37dl, dorsolateral area37 |
| 86 | Temporal Lobe | MTG, Middle Temporal Gyrus | A37dl, dorsolateral area37 |
| 87 | Temporal Lobe | MTG, Middle Temporal Gyrus | aSTS, anterior superior temporal sulcus |
| 88 | Temporal Lobe | MTG, Middle Temporal Gyrus | aSTS, anterior superior temporal sulcus |
| 90 | Temporal Lobe | ITG, Inferior Temporal Gyrus | A20iv, intermediate ventral area 20 |
| 91 | Temporal Lobe | ITG, Inferior Temporal Gyrus | A37elv, extreme lateroventral area37 |
| 92 | Temporal Lobe | ITG, Inferior Temporal Gyrus | A37elv, extreme lateroventral area37 |

| | | | |
|---|---|---|---|
| 93 | Temporal Lobe | ITG, Inferior Temporal Gyrus | A20r, rostral area 20 |
| 94 | Temporal Lobe | ITG, Inferior Temporal Gyrus | A20r, rostral area 20 |
| 95 | Temporal Lobe | ITG, Inferior Temporal Gyrus | A20il, intermediate lateral area 20 |
| 96 | Temporal Lobe | ITG, Inferior Temporal Gyrus | A20il, intermediate lateral area 20 |
| 97 | Temporal Lobe | ITG, Inferior Temporal Gyrus | A37vl, ventrolateral area 37 |
| 98 | Temporal Lobe | ITG, Inferior Temporal Gyrus | A37vl, ventrolateral area 37 |
| 99 | Temporal Lobe | ITG, Inferior Temporal Gyrus | A20cl, caudolateral of area 20 |
| 100 | Temporal Lobe | ITG, Inferior Temporal Gyrus | A20cl, caudolateral of area 20 |
| 101 | Temporal Lobe | ITG, Inferior Temporal Gyrus | A20cv, caudoventral of area 20 |
| 102 | Temporal Lobe | ITG, Inferior Temporal Gyrus | A20cv, caudoventral of area 20 |
| 103 | Temporal Lobe | FuG, Fusiform Gyrus | A20rv, rostroventral area 20 |
| 104 | Temporal Lobe | FuG, Fusiform Gyrus | A20rv, rostroventral area 20 |

| | | | |
|---|---|---|---|
| 108 | Temporal Lobe | FuG, Fusiform Gyrus | A37lv, lateroventral area37 |
| 135 | Parietal Lobe | IPL, Inferior Parietal Lobule | A39c, caudal area 39(PGp) |
| 136 | Parietal Lobe | IPL, Inferior Parietal Lobule | A39c, caudal area 39(PGp) |
| 137 | Parietal Lobe | IPL, Inferior Parietal Lobule | A39rd, rostrodorsal area 39(Hip3) |
| 138 | Parietal Lobe | IPL, Inferior Parietal Lobule | A39rd, rostrodorsal area 39(Hip3) |
| 141 | Parietal Lobe | IPL, Inferior Parietal Lobule | A40c, caudal area 40(PFm) |
| 143 | Parietal Lobe | IPL, Inferior Parietal Lobule | A39rv, rostroventral area 39(PGa) |
| 144 | Parietal Lobe | IPL, Inferior Parietal Lobule | A39rv, rostroventral area 39(PGa) |
| 151 | Parietal Lobe | Pcun, Precuneus | dmPOS, dorsomedial parietooccipital sulcus(PEr) |
| 152 | Parietal Lobe | Pcun, Precuneus | dmPOS, dorsomedial parietooccipital sulcus(PEr) |
| 153 | Parietal Lobe | Pcun, Precuneus | A31, area 31 (Lc1) |
| 154 | Parietal Lobe | Pcun, Precuneus | A31, area 31 (Lc1) |
| 175 | Limbic Lobe | CG, Cingulate Gyrus | A23d, dorsal area 23 |

| | | | |
|---|---|---|---|
| 178 | Limbic Lobe | CG, Cingulate Gyrus | A24rv, rostroventral area 24 |
| 179 | Limbic Lobe | CG, Cingulate Gyrus | A32p, pregenual area 32 |
| 180 | Limbic Lobe | CG, Cingulate Gyrus | A32p, pregenual area 32 |
| 183 | Limbic Lobe | CG, Cingulate Gyrus | A24cd, caudodorsal area 24 |
| 184 | Limbic Lobe | CG, Cingulate Gyrus | A24cd, caudodorsal area 24 |
| 185 | Limbic Lobe | CG, Cingulate Gyrus | A23c, caudal area 23 |
| 186 | Limbic Lobe | CG, Cingulate Gyrus | A23c, caudal area 23 |
| 187 | Limbic Lobe | CG, Cingulate Gyrus | A32sg, subgenual area 32 |
| 188 | Limbic Lobe | CG, Cingulate Gyrus | A32sg, subgenual area 32 |
| 190 | Occipital Lobe | MVOcC, MedioVentral Occipital Cortex | cLinG, caudal lingual gyrus |
| 209 | Occipital Lobe | LOcC, lateral Occipital Cortex | lsOccG, lateral superior occipital gyrus |
| 210 | Occipital Lobe | LOcC, lateral Occipital Cortex | lsOccG, lateral superior occipital gyrus |
| 212 | Subcortical Nuclei | Amyg, Amygdala | mAmyg, medial amygdala |
| 215 | Subcortical Nuclei | Hipp, Hippocampus | rHipp, rostral hippocampus |
| 216 | Subcortical Nuclei | Hipp, Hippocampus | rHipp, rostral hippocampus |

| | | | |
|---|---|---|---|
| 217 | Subcortical Nuclei | Hipp, Hippocampus | cHipp, caudal hippocampus |
| 218 | Subcortical Nuclei | Hipp, Hippocampus | cHipp, caudal hippocampus |
| 219 | Subcortical Nuclei | BG, Basal Ganglia | vCa, ventral caudate |
| 220 | Subcortical Nuclei | BG, Basal Ganglia | vCa, ventral caudate |
| 221 | Subcortical Nuclei | BG, Basal Ganglia | GP, globus pallidus |
| 222 | Subcortical Nuclei | BG, Basal Ganglia | GP, globus pallidus |
| 223 | Subcortical Nuclei | BG, Basal Ganglia | NAC, nucleus accumbens |
| 224 | Subcortical Nuclei | BG, Basal Ganglia | NAC, nucleus accumbens |
| 225 | Subcortical Nuclei | BG, Basal Ganglia | vmPu, ventromedial putamen |
| 226 | Subcortical Nuclei | BG, Basal Ganglia | vmPu, ventromedial putamen |
| 229 | Subcortical Nuclei | BG, Basal Ganglia | dlPu, dorsolateral putamen |
| 230 | Subcortical Nuclei | BG, Basal Ganglia | dlPu, dorsolateral putamen |
| 231 | Subcortical Nuclei | Tha, Thalamus | mPFtha, medial pre-frontal thalamus |
| 232 | Subcortical Nuclei | Tha, Thalamus | mPFtha, medial pre-frontal thalamus |
| 233 | Subcortical Nuclei | Tha, Thalamus | mPMtha, pre-motor thalamus |
| 234 | Subcortical Nuclei | Tha, Thalamus | mPMtha, pre-motor thalamus |

| | | | | |
|---|---|---|---|---|
| | 235 | Subcortical Nuclei | Tha, Thalamus | Stha, sensory thalamus |
| | 236 | Subcortical Nuclei | Tha, Thalamus | Stha, sensory thalamus |
| | 237 | Subcortical Nuclei | Tha, Thalamus | rTtha, rostral temporal thalamus |
| | 239 | Subcortical Nuclei | Tha, Thalamus | PPtha, posterior parietal thalamus |
| | 240 | Subcortical Nuclei | Tha, Thalamus | PPtha, posterior parietal thalamus |
| | 241 | Subcortical Nuclei | Tha, Thalamus | Otha, occipital thalamus |
| | 242 | Subcortical Nuclei | Tha, Thalamus | Otha, occipital thalamus |
| | 243 | Subcortical Nuclei | Tha, Thalamus | cTtha, caudal temporal thalamus |
| | 244 | Subcortical Nuclei | Tha, Thalamus | cTtha, caudal temporal thalamus |
| | 245 | Subcortical Nuclei | Tha, Thalamus | lPFtha, lateral pre-frontal thalamus |
| | 246 | Subcortical Nuclei | Tha, Thalamus | lPFtha, lateral pre-frontal thalamus |
| $SCZ_{C3}$ | 23 | Frontal Lobe | MFG, Middle Frontal Gyrus | A8vl, ventrolateral area 8 |
| | 24 | Frontal Lobe | MFG, Middle Frontal Gyrus | A8vl, ventrolateral area 8 |
| | 32 | Frontal Lobe | IFG, Inferior Frontal Gyrus | IFS, inferior frontal sulcus |
| | 63 | Frontal Lobe | PrG, Precentral Gyrus | A6cvl, caudal ventrolateral area 6 |

| | | | |
|---|---|---|---|
| 64 | Frontal Lobe | PrG, Precentral Gyrus | A6cvl, caudal ventrolateral area 6 |
| 71 | Temporal Lobe | STG, Superior Temporal Gyrus | A41/42, area 41/42 |
| 72 | Temporal Lobe | STG, Superior Temporal Gyrus | A41/42, area 41/42 |
| 73 | Temporal Lobe | STG, Superior Temporal Gyrus | TE1.0 and TE1.2 |
| 75 | Temporal Lobe | STG, Superior Temporal Gyrus | A22c, caudal area 22 |
| 76 | Temporal Lobe | STG, Superior Temporal Gyrus | A22c, caudal area 22 |
| 105 | Temporal Lobe | FuG, Fusiform Gyrus | A37mv, medioventral area37 |
| 106 | Temporal Lobe | FuG, Fusiform Gyrus | A37mv, medioventral area37 |
| 107 | Temporal Lobe | FuG, Fusiform Gyrus | A37lv, lateroventral area37 |
| 109 | Temporal Lobe | PhG, Parahippocampal Gyrus | A35/36r, rostral area 35/36 |
| 110 | Temporal Lobe | PhG, Parahippocampal Gyrus | A35/36r, rostral area 35/36 |
| 115 | Temporal Lobe | PhG, Parahippocampal Gyrus | A28/34, area 28/34 (EC, entorhinal cortex) |
| 116 | Temporal Lobe | PhG, Parahippocampal Gyrus | A28/34, area 28/34 (EC, entorhinal cortex) |

| | | | |
|---|---|---|---|
| 117 | Temporal Lobe | PhG, Parahippocampal Gyrus | TI, area TI(temporal agranular insular cortex) |
| 118 | Temporal Lobe | PhG, Parahippocampal Gyrus | TI, area TI(temporal agranular insular cortex) |
| 119 | Temporal Lobe | PhG, Parahippocampal Gyrus | TH, area TH (medial PPHC) |
| 120 | Temporal Lobe | PhG, Parahippocampal Gyrus | TH, area TH (medial PPHC) |
| 121 | Temporal Lobe | pSTS, posterior Superior Temporal Sulcus | rpSTS, rostroposterior superior temporal sulcus |
| 122 | Temporal Lobe | pSTS, posterior Superior Temporal Sulcus | rpSTS, rostroposterior superior temporal sulcus |
| 123 | Temporal Lobe | pSTS, posterior Superior Temporal Sulcus | cpSTS, caudoposterior superior temporal sulcus |
| 124 | Temporal Lobe | pSTS, posterior Superior Temporal Sulcus | cpSTS, caudoposterior superior temporal sulcus |
| 125 | Parietal Lobe | SPL, Superior Parietal Lobule | A7r, rostral area 7 |
| 126 | Parietal Lobe | SPL, Superior Parietal Lobule | A7r, rostral area 7 |
| 127 | Parietal Lobe | SPL, Superior Parietal Lobule | A7c, caudal area 7 |

| | | | |
|---|---|---|---|
| 128 | Parietal Lobe | SPL, Superior Parietal Lobule | A7c, caudal area 7 |
| 129 | Parietal Lobe | SPL, Superior Parietal Lobule | A5l, lateral area 5 |
| 130 | Parietal Lobe | SPL, Superior Parietal Lobule | A5l, lateral area 5 |
| 131 | Parietal Lobe | SPL, Superior Parietal Lobule | A7pc, postcentral area 7 |
| 132 | Parietal Lobe | SPL, Superior Parietal Lobule | A7pc, postcentral area 7 |
| 133 | Parietal Lobe | SPL, Superior Parietal Lobule | A7ip, intraparietal area 7(hIP3) |
| 134 | Parietal Lobe | SPL, Superior Parietal Lobule | A7ip, intraparietal area 7(hIP3) |
| 139 | Parietal Lobe | IPL, Inferior Parietal Lobule | A40rd, rostrodorsal area 40(PFt) |
| 140 | Parietal Lobe | IPL, Inferior Parietal Lobule | A40rd, rostrodorsal area 40(PFt) |
| 142 | Parietal Lobe | IPL, Inferior Parietal Lobule | A40c, caudal area 40(PFm) |
| 145 | Parietal Lobe | IPL, Inferior Parietal Lobule | A40rv, rostroventral area 40(PFop) |
| 146 | Parietal Lobe | IPL, Inferior Parietal Lobule | A40rv, rostroventral area 40(PFop) |
| 147 | Parietal Lobe | Pcun, Precuneus | A7m, medial area 7(PEp) |
| 148 | Parietal Lobe | Pcun, Precuneus | A7m, medial area 7(PEp) |

| | | | |
|---|---|---|---|
| 149 | Parietal Lobe | Pcun, Precuneus | A5m, medial area 5(PEm) |
| 150 | Parietal Lobe | Pcun, Precuneus | A5m, medial area 5(PEm) |
| 155 | Parietal Lobe | PoG, Postcentral Gyrus | A1/2/3ulhf, area 1/2/3(upper limb, head and face region) |
| 156 | Parietal Lobe | PoG, Postcentral Gyrus | A1/2/3ulhf, area 1/2/3(upper limb, head and face region) |
| 157 | Parietal Lobe | PoG, Postcentral Gyrus | A1/2/3tonIa, area 1/2/3(tongue and larynx region) |
| 158 | Parietal Lobe | PoG, Postcentral Gyrus | A1/2/3tonIa, area 1/2/3(tongue and larynx region) |
| 169 | Insular Lobe | INS, Insular Gyrus | vId/vIg, ventral dysgranular and granular insula |
| 170 | Insular Lobe | INS, Insular Gyrus | vId/vIg, ventral dysgranular and granular insula |
| 189 | Occipital Lobe | MVOcC, MedioVentral Occipital Cortex | cLinG, caudal lingual gyrus |
| 191 | Occipital Lobe | MVOcC, MedioVentral Occipital Cortex | rCunG, rostral cuneus gyrus |
| 192 | Occipital Lobe | MVOcC, MedioVentral Occipital Cortex | rCunG, rostral cuneus gyrus |

| | | | |
|---|---|---|---|
| 193 | Occipital Lobe | MVOcC, MedioVentral Occipital Cortex | cCunG, caudal cuneus gyrus |
| 194 | Occipital Lobe | MVOcC, MedioVentral Occipital Cortex | cCunG, caudal cuneus gyrus |
| 195 | Occipital Lobe | MVOcC, MedioVentral Occipital Cortex | rLinG, rostral lingual gyrus |
| 196 | Occipital Lobe | MVOcC, MedioVentral Occipital Cortex | rLinG, rostral lingual gyrus |
| 197 | Occipital Lobe | MVOcC, MedioVentral Occipital Cortex | vmPOS,ventromedial parietooccipital sulcus |
| 198 | Occipital Lobe | MVOcC, MedioVentral Occipital Cortex | vmPOS,ventromedial parietooccipital sulcus |
| 199 | Occipital Lobe | LOcC, lateral Occipital Cortex | mOccG, middle occipital gyrus |
| 200 | Occipital Lobe | LOcC, lateral Occipital Cortex | mOccG, middle occipital gyrus |
| 201 | Occipital Lobe | LOcC, lateral Occipital Cortex | V5/MT+, area V5/MT+ |
| 202 | Occipital Lobe | LOcC, lateral Occipital Cortex | V5/MT+, area V5/MT+ |

| | | | |
|---|---|---|---|
| 203 | Occipital Lobe | LOcC, lateral Occipital Cortex | OPC, occipital polar cortex |
| 204 | Occipital Lobe | LOcC, lateral Occipital Cortex | OPC, occipital polar cortex |
| 205 | Occipital Lobe | LOcC, lateral Occipital Cortex | iOccG, inferior occipital gyrus |
| 206 | Occipital Lobe | LOcC, lateral Occipital Cortex | iOccG, inferior occipital gyrus |
| 207 | Occipital Lobe | LOcC, lateral Occipital Cortex | msOccG, medial superior occipital gyrus |
| 208 | Occipital Lobe | LOcC, lateral Occipital Cortex | msOccG, medial superior occipital gyrus |
| 211 | Subcortical Nuclei | Amyg, Amygdala | mAmyg, medial amygdala |
| 213 | Subcortical Nuclei | Amyg, Amygdala | lAmyg, lateral amygdala |
| 214 | Subcortical Nuclei | Amyg, Amygdala | lAmyg, lateral amygdala |
| 227 | Subcortical Nuclei | BG, Basal Ganglia | dCa, dorsal caudate |
| 228 | Subcortical Nuclei | BG, Basal Ganglia | dCa, dorsal caudate |
| 238 | Subcortical Nuclei | Tha, Thalamus | rTtha, rostral temporal thalamus |

Supplementary Table 2. The table provides a detailed listing of the brain regions assigned to each cluster formed from applying AHC to the fMRI time series data.

| Subnetwork | Brainnetome Region Number | Lobe | Gyrus | Anatomical and modified Cyto-architectonic descriptions |
|---|---|---|---|---|
| $HC_{fMRI1}$ | 45 | Frontal Lobe | OrG, Orbital Gyrus | A11l, lateral area 11 |
| | 46 | Frontal Lobe | OrG, Orbital Gyrus | A11l, lateral area 11 |
| | 47 | Frontal Lobe | OrG, Orbital Gyrus | A11m, medial area 11 |
| | 48 | Frontal Lobe | OrG, Orbital Gyrus | A11m, medial area 11 |
| | 49 | Frontal Lobe | OrG, Orbital Gyrus | A13, area 13 |
| | 50 | Frontal Lobe | OrG, Orbital Gyrus | A13, area 13 |
| | 73 | Temporal Lobe | STG, Superior Temporal Gyrus | TE1.0 and TE1.2 |
| | 74 | Temporal Lobe | STG, Superior Temporal Gyrus | TE1.0 and TE1.2 |
| | 122 | Temporal Lobe | pSTS, posterior Superior Temporal Sulcus | rpSTS, rostroposterior superior temporal sulcus |
| | 165 | Insular Lobe | INS, Insular Gyrus | vIa, ventral agranular insula |
| | 166 | Insular Lobe | INS, Insular Gyrus | vIa, ventral agranular insula |
| | 167 | Insular Lobe | INS, Insular Gyrus | dIa, dorsal agranular insula |
| | 168 | Insular Lobe | INS, Insular Gyrus | dIa, dorsal agranular insula |

| | | | |
|---|---|---|---|
| 169 | Insular Lobe | INS, Insular Gyrus | vId/vIg, ventral dysgranular and granular insula |
| 171 | Insular Lobe | INS, Insular Gyrus | dIg, dorsal granular insula |
| 173 | Insular Lobe | INS, Insular Gyrus | dId, dorsal dysgranular insula |
| 175 | Limbic Lobe | CG, Cingulate Gyrus | A23d, dorsal area 23 |
| 176 | Limbic Lobe | CG, Cingulate Gyrus | A23d, dorsal area 23 |
| 181 | Limbic Lobe | CG, Cingulate Gyrus | A23v, ventral area 23 |
| 182 | Limbic Lobe | CG, Cingulate Gyrus | A23v, ventral area 23 |
| 183 | Limbic Lobe | CG, Cingulate Gyrus | A24cd, caudodorsal area 24 |
| 184 | Limbic Lobe | CG, Cingulate Gyrus | A24cd, caudodorsal area 24 |
| 185 | Limbic Lobe | CG, Cingulate Gyrus | A23c, caudal area 23 |
| 186 | Limbic Lobe | CG, Cingulate Gyrus | A23c, caudal area 23 |
| 187 | Limbic Lobe | CG, Cingulate Gyrus | A32sg, subgenual area 32 |
| 189 | Occipital Lobe | MVOcC, MedioVentral Occipital Cortex | cLinG, caudal lingual gyrus |
| 190 | Occipital Lobe | MVOcC, MedioVentral Occipital Cortex | cLinG, caudal lingual gyrus |
| 191 | Occipital Lobe | MVOcC, MedioVentral | rCunG, rostral cuneus gyrus |

| | | | |
|---|---|---|---|
| | | Occipital Cortex | |
| 192 | Occipital Lobe | MVOcC, MedioVentral Occipital Cortex | rCunG, rostral cuneus gyrus |
| 193 | Occipital Lobe | MVOcC, MedioVentral Occipital Cortex | cCunG, caudal cuneus gyrus |
| 194 | Occipital Lobe | MVOcC, MedioVentral Occipital Cortex | cCunG, caudal cuneus gyrus |
| 195 | Occipital Lobe | MVOcC, MedioVentral Occipital Cortex | rLinG, rostral lingual gyrus |
| 196 | Occipital Lobe | MVOcC, MedioVentral Occipital Cortex | rLinG, rostral lingual gyrus |
| 197 | Occipital Lobe | MVOcC, MedioVentral Occipital Cortex | vmPOS,ventromedial parietooccipital sulcus |
| 198 | Occipital Lobe | MVOcC, MedioVentral Occipital Cortex | vmPOS,ventromedial parietooccipital sulcus |
| 212 | Subcortical Nuclei | Amyg, Amygdala | mAmyg, medial amygdala |
| 213 | Subcortical Nuclei | Amyg, Amygdala | lAmyg, lateral amygdala |
| 214 | Subcortical Nuclei | Amyg, Amygdala | lAmyg, lateral amygdala |

| | | | | |
|---|---|---|---|---|
| $HC_{fMRI2}$ | 170 | Insular Lobe | INS, Insular Gyrus | vId/vIg, ventral dysgranular and granular insula |
| | 215 | Subcortical Nuclei | Hipp, Hippocampus | rHipp, rostral hippocampus |
| | 216 | Subcortical Nuclei | Hipp, Hippocampus | rHipp, rostral hippocampus |
| | 217 | Subcortical Nuclei | Hipp, Hippocampus | cHipp, caudal hippocampus |
| | 218 | Subcortical Nuclei | Hipp, Hippocampus | cHipp, caudal hippocampus |
| | 219 | Subcortical Nuclei | BG, Basal Ganglia | vCa, ventral caudate |
| | 220 | Subcortical Nuclei | BG, Basal Ganglia | vCa, ventral caudate |
| | 221 | Subcortical Nuclei | BG, Basal Ganglia | GP, globus pallidus |
| | 222 | Subcortical Nuclei | BG, Basal Ganglia | GP, globus pallidus |
| | 223 | Subcortical Nuclei | BG, Basal Ganglia | NAC, nucleus accumbens |
| | 224 | Subcortical Nuclei | BG, Basal Ganglia | NAC, nucleus accumbens |
| | 225 | Subcortical Nuclei | BG, Basal Ganglia | vmPu, ventromedial putamen |
| | 226 | Subcortical Nuclei | BG, Basal Ganglia | vmPu, ventromedial putamen |
| | 227 | Subcortical Nuclei | BG, Basal Ganglia | dCa, dorsal caudate |
| | 228 | Subcortical Nuclei | BG, Basal Ganglia | dCa, dorsal caudate |
| | 229 | Subcortical Nuclei | BG, Basal Ganglia | dlPu, dorsolateral putamen |

| | | | | |
|---|---|---|---|---|
| | 230 | Subcortical Nuclei | BG, Basal Ganglia | dlPu, dorsolateral putamen |
| | 231 | Subcortical Nuclei | Tha, Thalamus | mPFtha, medial pre-frontal thalamus |
| | 232 | Subcortical Nuclei | Tha, Thalamus | mPFtha, medial pre-frontal thalamus |
| | 233 | Subcortical Nuclei | Tha, Thalamus | mPMtha, pre-motor thalamus |
| | 234 | Subcortical Nuclei | Tha, Thalamus | mPMtha, pre-motor thalamus |
| $HC_{fMRI3}$ | 1 | Frontal Lobe | SFG, Superior Frontal Gyrus | A8m, medial area 8 |
| | 3 | Frontal Lobe | SFG, Superior Frontal Gyrus | A8dl, dorsolateral area 8 |
| | 4 | Frontal Lobe | SFG, Superior Frontal Gyrus | A8dl, dorsolateral area 8 |
| | 5 | Frontal Lobe | SFG, Superior Frontal Gyrus | A9l, lateral area 9 |
| | 6 | Frontal Lobe | SFG, Superior Frontal Gyrus | A9l, lateral area 9 |
| | 12 | Frontal Lobe | SFG, Superior Frontal Gyrus | A9m,medial area 9 |
| | 13 | Frontal Lobe | SFG, Superior Frontal Gyrus | A10m, medial area 10 |
| | 14 | Frontal Lobe | SFG, Superior Frontal Gyrus | A10m, medial area 10 |
| | 15 | Frontal Lobe | MFG, Middle Frontal Gyrus | A9/46d, dorsal area 9/46 |
| | 16 | Frontal Lobe | MFG, Middle Frontal Gyrus | A9/46d, dorsal area 9/46 |
| | 20 | Frontal Lobe | MFG, Middle Frontal Gyrus | A46, area 46 |

| | | | |
|---|---|---|---|
| 21 | Frontal Lobe | MFG, Middle Frontal Gyrus | A9/46v, ventral area 9/46 |
| 22 | Frontal Lobe | MFG, Middle Frontal Gyrus | A9/46v, ventral area 9/46 |
| 23 | Frontal Lobe | MFG, Middle Frontal Gyrus | A8vl, ventrolateral area 8 |
| 24 | Frontal Lobe | MFG, Middle Frontal Gyrus | A8vl, ventrolateral area 8 |
| 27 | Frontal Lobe | MFG, Middle Frontal Gyrus | A10l, lateral area10 |
| 28 | Frontal Lobe | MFG, Middle Frontal Gyrus | A10l, lateral area10 |
| 30 | Frontal Lobe | IFG, Inferior Frontal Gyrus | A44d,dorsal area 44 |
| 31 | Frontal Lobe | IFG, Inferior Frontal Gyrus | IFS, inferior frontal sulcus |
| 32 | Frontal Lobe | IFG, Inferior Frontal Gyrus | IFS, inferior frontal sulcus |
| 33 | Frontal Lobe | IFG, Inferior Frontal Gyrus | A45c, caudal area 45 |
| 34 | Frontal Lobe | IFG, Inferior Frontal Gyrus | A45c, caudal area 45 |
| 35 | Frontal Lobe | IFG, Inferior Frontal Gyrus | A45r, rostral area 45 |
| 36 | Frontal Lobe | IFG, Inferior Frontal Gyrus | A45r, rostral area 45 |
| 40 | Frontal Lobe | IFG, Inferior Frontal Gyrus | A44v, ventral area 44 |
| 52 | Frontal Lobe | OrG, Orbital Gyrus | A12/47l, lateral area 12/47 |
| 66 | Frontal Lobe | PCL, Paracentral Lobule | A1/2/3ll, area1/2/3 (lower limb region) |

| | | | |
|---|---|---|---|
| 67 | Frontal Lobe | PCL, Paracentral Lobule | A4ll, area 4, (lower limb region) |
| 68 | Frontal Lobe | PCL, Paracentral Lobule | A4ll, area 4, (lower limb region) |
| 69 | Temporal Lobe | STG, Superior Temporal Gyrus | A38m, medial area 38 |
| 70 | Temporal Lobe | STG, Superior Temporal Gyrus | A38m, medial area 38 |
| 71 | Temporal Lobe | STG, Superior Temporal Gyrus | A41/42, area 41/42 |
| 81 | Temporal Lobe | MTG, Middle Temporal Gyrus | A21c, caudal area 21 |
| 82 | Temporal Lobe | MTG, Middle Temporal Gyrus | A21c, caudal area 21 |
| 83 | Temporal Lobe | MTG, Middle Temporal Gyrus | A21r, rostral area 21 |
| 84 | Temporal Lobe | MTG, Middle Temporal Gyrus | A21r, rostral area 21 |
| 85 | Temporal Lobe | MTG, Middle Temporal Gyrus | A37dl, dorsolateral area37 |
| 86 | Temporal Lobe | MTG, Middle Temporal Gyrus | A37dl, dorsolateral area37 |
| 89 | Temporal Lobe | ITG, Inferior Temporal Gyrus | A20iv, intermediate ventral area 20 |
| 90 | Temporal Lobe | ITG, Inferior Temporal Gyrus | A20iv, intermediate ventral area 20 |
| 91 | Temporal Lobe | ITG, Inferior Temporal Gyrus | A37elv, extreme lateroventral area37 |
| 92 | Temporal Lobe | ITG, Inferior Temporal Gyrus | A37elv, extreme lateroventral area37 |

| | | | |
|---|---|---|---|
| 93 | Temporal Lobe | ITG, Inferior Temporal Gyrus | A20r, rostral area 20 |
| 94 | Temporal Lobe | ITG, Inferior Temporal Gyrus | A20r, rostral area 20 |
| 95 | Temporal Lobe | ITG, Inferior Temporal Gyrus | A20il, intermediate lateral area 20 |
| 96 | Temporal Lobe | ITG, Inferior Temporal Gyrus | A20il, intermediate lateral area 20 |
| 97 | Temporal Lobe | ITG, Inferior Temporal Gyrus | A37vl, ventrolateral area 37 |
| 99 | Temporal Lobe | ITG, Inferior Temporal Gyrus | A20cl, caudolateral of area 20 |
| 100 | Temporal Lobe | ITG, Inferior Temporal Gyrus | A20cl, caudolateral of area 20 |
| 101 | Temporal Lobe | ITG, Inferior Temporal Gyrus | A20cv, caudoventral of area 20 |
| 102 | Temporal Lobe | ITG, Inferior Temporal Gyrus | A20cv, caudoventral of area 20 |
| 103 | Temporal Lobe | FuG, Fusiform Gyrus | A20rv, rostroventral area 20 |
| 104 | Temporal Lobe | FuG, Fusiform Gyrus | A20rv, rostroventral area 20 |
| 109 | Temporal Lobe | PhG, Parahippocampal Gyrus | A35/36r, rostral area 35/36 |
| 110 | Temporal Lobe | PhG, Parahippocampal Gyrus | A35/36r, rostral area 35/36 |
| 111 | Temporal Lobe | PhG, Parahippocampal Gyrus | A35/36c, caudal area 35/36 |

| | | | |
|---|---|---|---|
| 116 | Temporal Lobe | PhG, Parahippocampal Gyrus | A28/34, area 28/34 (EC, entorhinal cortex) |
| 118 | Temporal Lobe | PhG, Parahippocampal Gyrus | TI, area TI(temporal agranular insular cortex) |
| 131 | Parietal Lobe | SPL, Superior Parietal Lobule | A7pc, postcentral area 7 |
| 132 | Parietal Lobe | SPL, Superior Parietal Lobule | A7pc, postcentral area 7 |
| 135 | Parietal Lobe | IPL, Inferior Parietal Lobule | A39c, caudal area 39(PGp) |
| 136 | Parietal Lobe | IPL, Inferior Parietal Lobule | A39c, caudal area 39(PGp) |
| 137 | Parietal Lobe | IPL, Inferior Parietal Lobule | A39rd, rostrodorsal area 39(Hip3) |
| 138 | Parietal Lobe | IPL, Inferior Parietal Lobule | A39rd, rostrodorsal area 39(Hip3) |
| 139 | Parietal Lobe | IPL, Inferior Parietal Lobule | A40rd, rostrodorsal area 40(PFt) |
| 140 | Parietal Lobe | IPL, Inferior Parietal Lobule | A40rd, rostrodorsal area 40(PFt) |
| 141 | Parietal Lobe | IPL, Inferior Parietal Lobule | A40c, caudal area 40(PFm) |
| 142 | Parietal Lobe | IPL, Inferior Parietal Lobule | A40c, caudal area 40(PFm) |
| 143 | Parietal Lobe | IPL, Inferior Parietal Lobule | A39rv, rostroventral area 39(PGa) |
| 144 | Parietal Lobe | IPL, Inferior Parietal Lobule | A39rv, rostroventral area 39(PGa) |

| | | | | |
|---|---|---|---|---|
| | 145 | Parietal Lobe | IPL, Inferior Parietal Lobule | A40rv, rostroventral area 40(PFop) |
| | 146 | Parietal Lobe | IPL, Inferior Parietal Lobule | A40rv, rostroventral area 40(PFop) |
| | 161 | Parietal Lobe | PoG, Postcentral Gyrus | A1/2/3tru, area1/2/3(trunk region) |
| | 200 | Occipital Lobe | LOcC, lateral Occipital Cortex | mOccG, middle occipital gyrus |
| | 201 | Occipital Lobe | LOcC, lateral Occipital Cortex | V5/MT+, area V5/MT+ |
| $HC_{fMRI4}$ | 235 | Subcortical Nuclei | Tha, Thalamus | Stha, sensory thalamus |
| | 236 | Subcortical Nuclei | Tha, Thalamus | Stha, sensory thalamus |
| | 237 | Subcortical Nuclei | Tha, Thalamus | rTtha, rostral temporal thalamus |
| | 238 | Subcortical Nuclei | Tha, Thalamus | rTtha, rostral temporal thalamus |
| | 239 | Subcortical Nuclei | Tha, Thalamus | PPtha, posterior parietal thalamus |
| | 240 | Subcortical Nuclei | Tha, Thalamus | PPtha, posterior parietal thalamus |
| | 241 | Subcortical Nuclei | Tha, Thalamus | Otha, occipital thalamus |
| | 242 | Subcortical Nuclei | Tha, Thalamus | Otha, occipital thalamus |
| | 243 | Subcortical Nuclei | Tha, Thalamus | cTtha, caudal temporal thalamus |
| | 244 | Subcortical Nuclei | Tha, Thalamus | cTtha, caudal temporal thalamus |

| | | | | |
|---|---|---|---|---|
| | 245 | Subcortical Nuclei | Tha, Thalamus | lPFtha, lateral pre-frontal thalamus |
| | 246 | Subcortical Nuclei | Tha, Thalamus | lPFtha, lateral pre-frontal thalamus |
| $HC_{fMRI5}$ | 2 | Frontal Lobe | SFG, Superior Frontal Gyrus | A8m, medial area 8 |
| | 7 | Frontal Lobe | SFG, Superior Frontal Gyrus | A6dl, dorsolateral area 6 |
| | 8 | Frontal Lobe | SFG, Superior Frontal Gyrus | A6dl, dorsolateral area 6 |
| | 9 | Frontal Lobe | SFG, Superior Frontal Gyrus | A6m, medial area 6 |
| | 10 | Frontal Lobe | SFG, Superior Frontal Gyrus | A6m, medial area 6 |
| | 11 | Frontal Lobe | SFG, Superior Frontal Gyrus | A9m,medial area 9 |
| | 17 | Frontal Lobe | MFG, Middle Frontal Gyrus | IFJ, inferior frontal junction |
| | 18 | Frontal Lobe | MFG, Middle Frontal Gyrus | IFJ, inferior frontal junction |
| | 19 | Frontal Lobe | MFG, Middle Frontal Gyrus | A46, area 46 |
| | 25 | Frontal Lobe | MFG, Middle Frontal Gyrus | A6vl, ventrolateral area 6 |
| | 26 | Frontal Lobe | MFG, Middle Frontal Gyrus | A6vl, ventrolateral area 6 |
| | 29 | Frontal Lobe | IFG, Inferior Frontal Gyrus | A44d,dorsal area 44 |
| | 37 | Frontal Lobe | IFG, Inferior Frontal Gyrus | A44op, opercular area 44 |
| | 38 | Frontal Lobe | IFG, Inferior Frontal Gyrus | A44op, opercular area 44 |

| | | | |
|---|---|---|---|
| 39 | Frontal Lobe | IFG, Inferior Frontal Gyrus | A44v, ventral area 44 |
| 41 | Frontal Lobe | OrG, Orbital Gyrus | A14m, medial area 14 |
| 42 | Frontal Lobe | OrG, Orbital Gyrus | A14m, medial area 14 |
| 43 | Frontal Lobe | OrG, Orbital Gyrus | A12/47o, orbital area 12/47 |
| 44 | Frontal Lobe | OrG, Orbital Gyrus | A12/47o, orbital area 12/47 |
| 51 | Frontal Lobe | OrG, Orbital Gyrus | A12/47l, lateral area 12/47 |
| 53 | Frontal Lobe | PrG, Precentral Gyrus | A4hf, area 4(head and face region) |
| 54 | Frontal Lobe | PrG, Precentral Gyrus | A4hf, area 4(head and face region) |
| 55 | Frontal Lobe | PrG, Precentral Gyrus | A6cdl, caudal dorsolateral area 6 |
| 56 | Frontal Lobe | PrG, Precentral Gyrus | A6cdl, caudal dorsolateral area 6 |
| 57 | Frontal Lobe | PrG, Precentral Gyrus | A4ul, area 4(upper limb region) |
| 58 | Frontal Lobe | PrG, Precentral Gyrus | A4ul, area 4(upper limb region) |
| 59 | Frontal Lobe | PrG, Precentral Gyrus | A4t, area 4(trunk region) |
| 60 | Frontal Lobe | PrG, Precentral Gyrus | A4t, area 4(trunk region) |
| 61 | Frontal Lobe | PrG, Precentral Gyrus | A4tl, area 4(tongue and larynx region) |
| 62 | Frontal Lobe | PrG, Precentral Gyrus | A4tl, area 4(tongue and larynx region) |

| | | | |
|---|---|---|---|
| 63 | Frontal Lobe | PrG, Precentral Gyrus | A6cvl, caudal ventrolateral area 6 |
| 64 | Frontal Lobe | PrG, Precentral Gyrus | A6cvl, caudal ventrolateral area 6 |
| 65 | Frontal Lobe | PCL, Paracentral Lobule | A1/2/3ll, area1/2/3 (lower limb region) |
| 72 | Temporal Lobe | STG, Superior Temporal Gyrus | A41/42, area 41/42 |
| 75 | Temporal Lobe | STG, Superior Temporal Gyrus | A22c, caudal area 22 |
| 76 | Temporal Lobe | STG, Superior Temporal Gyrus | A22c, caudal area 22 |
| 77 | Temporal Lobe | STG, Superior Temporal Gyrus | A38l, lateral area 38 |
| 78 | Temporal Lobe | STG, Superior Temporal Gyrus | A38l, lateral area 38 |
| 79 | Temporal Lobe | STG, Superior Temporal Gyrus | A22r, rostral area 22 |
| 80 | Temporal Lobe | STG, Superior Temporal Gyrus | A22r, rostral area 22 |
| 87 | Temporal Lobe | MTG, Middle Temporal Gyrus | aSTS, anterior superior temporal sulcus |
| 88 | Temporal Lobe | MTG, Middle Temporal Gyrus | aSTS, anterior superior temporal sulcus |
| 98 | Temporal Lobe | ITG, Inferior Temporal Gyrus | A37vl, ventrolateral area 37 |
| 105 | Temporal Lobe | FuG, Fusiform Gyrus | A37mv, medioventral area37 |

| | | | |
|---|---|---|---|
| 106 | Temporal Lobe | FuG, Fusiform Gyrus | A37mv, medioventral area37 |
| 107 | Temporal Lobe | FuG, Fusiform Gyrus | A37lv, lateroventral area37 |
| 108 | Temporal Lobe | FuG, Fusiform Gyrus | A37lv, lateroventral area37 |
| 112 | Temporal Lobe | PhG, Parahippocampal Gyrus | A35/36c, caudal area 35/36 |
| 113 | Temporal Lobe | PhG, Parahippocampal Gyrus | TL, area TL (lateral PPHC, posterior parahippocampal gyrus) |
| 114 | Temporal Lobe | PhG, Parahippocampal Gyrus | TL, area TL (lateral PPHC, posterior parahippocampal gyrus) |
| 115 | Temporal Lobe | PhG, Parahippocampal Gyrus | A28/34, area 28/34 (EC, entorhinal cortex) |
| 117 | Temporal Lobe | PhG, Parahippocampal Gyrus | TI, area TI(temporal agranular insular cortex) |
| 119 | Temporal Lobe | PhG, Parahippocampal Gyrus | TH, area TH (medial PPHC) |
| 120 | Temporal Lobe | PhG, Parahippocampal Gyrus | TH, area TH (medial PPHC) |
| 121 | Temporal Lobe | pSTS, posterior Superior Temporal Sulcus | rpSTS, rostroposterior superior temporal sulcus |

| | | | |
|---|---|---|---|
| 123 | Temporal Lobe | pSTS, posterior Superior Temporal Sulcus | cpSTS, caudoposterior superior temporal sulcus |
| 124 | Temporal Lobe | pSTS, posterior Superior Temporal Sulcus | cpSTS, caudoposterior superior temporal sulcus |
| 125 | Parietal Lobe | SPL, Superior Parietal Lobule | A7r, rostral area 7 |
| 126 | Parietal Lobe | SPL, Superior Parietal Lobule | A7r, rostral area 7 |
| 127 | Parietal Lobe | SPL, Superior Parietal Lobule | A7c, caudal area 7 |
| 128 | Parietal Lobe | SPL, Superior Parietal Lobule | A7c, caudal area 7 |
| 129 | Parietal Lobe | SPL, Superior Parietal Lobule | A5l, lateral area 5 |
| 130 | Parietal Lobe | SPL, Superior Parietal Lobule | A5l, lateral area 5 |
| 133 | Parietal Lobe | SPL, Superior Parietal Lobule | A7ip, intraparietal area 7(hIP3) |
| 134 | Parietal Lobe | SPL, Superior Parietal Lobule | A7ip, intraparietal area 7(hIP3) |
| 147 | Parietal Lobe | Pcun, Precuneus | A7m, medial area 7(PEp) |
| 148 | Parietal Lobe | Pcun, Precuneus | A7m, medial area 7(PEp) |
| 149 | Parietal Lobe | Pcun, Precuneus | A5m, medial area 5(PEm) |
| 150 | Parietal Lobe | Pcun, Precuneus | A5m, medial area 5(PEm) |
| 151 | Parietal Lobe | Pcun, Precuneus | dmPOS, dorsomedial parietooccipital sulcus(PEr) |

| | | | |
|---|---|---|---|
| 152 | Parietal Lobe | Pcun, Precuneus | dmPOS, dorsomedial parietooccipital sulcus(PEr) |
| 153 | Parietal Lobe | Pcun, Precuneus | A31, area 31 (Lc1) |
| 154 | Parietal Lobe | Pcun, Precuneus | A31, area 31 (Lc1) |
| 155 | Parietal Lobe | PoG, Postcentral Gyrus | A1/2/3ulhf, area 1/2/3(upper limb, head and face region) |
| 156 | Parietal Lobe | PoG, Postcentral Gyrus | A1/2/3ulhf, area 1/2/3(upper limb, head and face region) |
| 157 | Parietal Lobe | PoG, Postcentral Gyrus | A1/2/3tonIa, area 1/2/3(tongue and larynx region) |
| 158 | Parietal Lobe | PoG, Postcentral Gyrus | A1/2/3tonIa, area 1/2/3(tongue and larynx region) |
| 159 | Parietal Lobe | PoG, Postcentral Gyrus | A2, area 2 |
| 160 | Parietal Lobe | PoG, Postcentral Gyrus | A2, area 2 |
| 162 | Parietal Lobe | PoG, Postcentral Gyrus | A1/2/3tru, area1/2/3(trunk region) |
| 163 | Insular Lobe | INS, Insular Gyrus | G, hypergranular insula |
| 164 | Insular Lobe | INS, Insular Gyrus | G, hypergranular insula |
| 172 | Insular Lobe | INS, Insular Gyrus | dIg, dorsal granular insula |

| | | | |
|---|---|---|---|
| 174 | Insular Lobe | INS, Insular Gyrus | dId, dorsal dysgranular insula |
| 177 | Limbic Lobe | CG, Cingulate Gyrus | A24rv, rostroventral area 24 |
| 178 | Limbic Lobe | CG, Cingulate Gyrus | A24rv, rostroventral area 24 |
| 179 | Limbic Lobe | CG, Cingulate Gyrus | A32p, pregenual area 32 |
| 180 | Limbic Lobe | CG, Cingulate Gyrus | A32p, pregenual area 32 |
| 188 | Limbic Lobe | CG, Cingulate Gyrus | A32sg, subgenual area 32 |
| 199 | Occipital Lobe | LOcC, lateral Occipital Cortex | mOccG, middle occipital gyrus |
| 202 | Occipital Lobe | LOcC, lateral Occipital Cortex | V5/MT+, area V5/MT+ |
| 203 | Occipital Lobe | LOcC, lateral Occipital Cortex | OPC, occipital polar cortex |
| 204 | Occipital Lobe | LOcC, lateral Occipital Cortex | OPC, occipital polar cortex |
| 205 | Occipital Lobe | LOcC, lateral Occipital Cortex | iOccG, inferior occipital gyrus |
| 206 | Occipital Lobe | LOcC, lateral Occipital Cortex | iOccG, inferior occipital gyrus |
| 207 | Occipital Lobe | LOcC, lateral Occipital Cortex | msOccG, medial superior occipital gyrus |

| | | | | |
|---|---|---|---|---|
| | 208 | Occipital Lobe | LOcC, lateral Occipital Cortex | msOccG, medial superior occipital gyrus |
| | 209 | Occipital Lobe | LOcC, lateral Occipital Cortex | lsOccG, lateral superior occipital gyrus |
| | 210 | Occipital Lobe | LOcC, lateral Occipital Cortex | lsOccG, lateral superior occipital gyrus |
| | 211 | Subcortical Nuclei | Amyg, Amygdala | mAmyg, medial amygdala |
| $SCZ_{fMRI1}$ | 1 | Frontal Lobe | SFG, Superior Frontal Gyrus | A8m, medial area 8 |
| | 2 | Frontal Lobe | SFG, Superior Frontal Gyrus | A8m, medial area 8 |
| | 3 | Frontal Lobe | SFG, Superior Frontal Gyrus | A8dl, dorsolateral area 8 |
| | 4 | Frontal Lobe | SFG, Superior Frontal Gyrus | A8dl, dorsolateral area 8 |
| | 5 | Frontal Lobe | SFG, Superior Frontal Gyrus | A9l, lateral area 9 |
| | 6 | Frontal Lobe | SFG, Superior Frontal Gyrus | A9l, lateral area 9 |
| | 11 | Frontal Lobe | SFG, Superior Frontal Gyrus | A9m,medial area 9 |
| | 12 | Frontal Lobe | SFG, Superior Frontal Gyrus | A9m,medial area 9 |
| | 13 | Frontal Lobe | SFG, Superior Frontal Gyrus | A10m, medial area 10 |
| | 14 | Frontal Lobe | SFG, Superior Frontal Gyrus | A10m, medial area 10 |
| | 15 | Frontal Lobe | MFG, Middle Frontal Gyrus | A9/46d, dorsal area 9/46 |

| | | | |
|---|---|---|---|
| 16 | Frontal Lobe | MFG, Middle Frontal Gyrus | A9/46d, dorsal area 9/46 |
| 17 | Frontal Lobe | MFG, Middle Frontal Gyrus | IFJ, inferior frontal junction |
| 18 | Frontal Lobe | MFG, Middle Frontal Gyrus | IFJ, inferior frontal junction |
| 19 | Frontal Lobe | MFG, Middle Frontal Gyrus | A46, area 46 |
| 20 | Frontal Lobe | MFG, Middle Frontal Gyrus | A46, area 46 |
| 21 | Frontal Lobe | MFG, Middle Frontal Gyrus | A9/46v, ventral area 9/46 |
| 22 | Frontal Lobe | MFG, Middle Frontal Gyrus | A9/46v, ventral area 9/46 |
| 23 | Frontal Lobe | MFG, Middle Frontal Gyrus | A8vl, ventrolateral area 8 |
| 24 | Frontal Lobe | MFG, Middle Frontal Gyrus | A8vl, ventrolateral area 8 |
| 26 | Frontal Lobe | MFG, Middle Frontal Gyrus | A6vl, ventrolateral area 6 |
| 27 | Frontal Lobe | MFG, Middle Frontal Gyrus | A10l, lateral area10 |
| 28 | Frontal Lobe | MFG, Middle Frontal Gyrus | A10l, lateral area10 |
| 29 | Frontal Lobe | IFG, Inferior Frontal Gyrus | A44d,dorsal area 44 |
| 30 | Frontal Lobe | IFG, Inferior Frontal Gyrus | A44d,dorsal area 44 |
| 31 | Frontal Lobe | IFG, Inferior Frontal Gyrus | IFS, inferior frontal sulcus |
| 32 | Frontal Lobe | IFG, Inferior Frontal Gyrus | IFS, inferior frontal sulcus |

| | | | |
|---|---|---|---|
| 33 | Frontal Lobe | IFG, Inferior Frontal Gyrus | A45c, caudal area 45 |
| 34 | Frontal Lobe | IFG, Inferior Frontal Gyrus | A45c, caudal area 45 |
| 35 | Frontal Lobe | IFG, Inferior Frontal Gyrus | A45r, rostral area 45 |
| 36 | Frontal Lobe | IFG, Inferior Frontal Gyrus | A45r, rostral area 45 |
| 37 | Frontal Lobe | IFG, Inferior Frontal Gyrus | A44op, opercular area 44 |
| 38 | Frontal Lobe | IFG, Inferior Frontal Gyrus | A44op, opercular area 44 |
| 40 | Frontal Lobe | IFG, Inferior Frontal Gyrus | A44v, ventral area 44 |
| 41 | Frontal Lobe | OrG, Orbital Gyrus | A14m, medial area 14 |
| 42 | Frontal Lobe | OrG, Orbital Gyrus | A14m, medial area 14 |
| 43 | Frontal Lobe | OrG, Orbital Gyrus | A12/47o, orbital area 12/47 |
| 44 | Frontal Lobe | OrG, Orbital Gyrus | A12/47o, orbital area 12/47 |
| 47 | Frontal Lobe | OrG, Orbital Gyrus | A11m, medial area 11 |
| 48 | Frontal Lobe | OrG, Orbital Gyrus | A11m, medial area 11 |
| 51 | Frontal Lobe | OrG, Orbital Gyrus | A12/47l, lateral area 12/47 |
| 52 | Frontal Lobe | OrG, Orbital Gyrus | A12/47l, lateral area 12/47 |
| 69 | Temporal Lobe | STG, Superior Temporal Gyrus | A38m, medial area 38 |

| | | | |
|---|---|---|---|
| 70 | Temporal Lobe | STG, Superior Temporal Gyrus | A38m, medial area 38 |
| 71 | Temporal Lobe | STG, Superior Temporal Gyrus | A41/42, area 41/42 |
| 75 | Temporal Lobe | STG, Superior Temporal Gyrus | A22c, caudal area 22 |
| 78 | Temporal Lobe | STG, Superior Temporal Gyrus | A38l, lateral area 38 |
| 80 | Temporal Lobe | STG, Superior Temporal Gyrus | A22r, rostral area 22 |
| 81 | Temporal Lobe | MTG, Middle Temporal Gyrus | A21c, caudal area 21 |
| 82 | Temporal Lobe | MTG, Middle Temporal Gyrus | A21c, caudal area 21 |
| 83 | Temporal Lobe | MTG, Middle Temporal Gyrus | A21r, rostral area 21 |
| 84 | Temporal Lobe | MTG, Middle Temporal Gyrus | A21r, rostral area 21 |
| 85 | Temporal Lobe | MTG, Middle Temporal Gyrus | A37dl, dorsolateral area37 |
| 86 | Temporal Lobe | MTG, Middle Temporal Gyrus | A37dl, dorsolateral area37 |
| 87 | Temporal Lobe | MTG, Middle Temporal Gyrus | aSTS, anterior superior temporal sulcus |
| 88 | Temporal Lobe | MTG, Middle Temporal Gyrus | aSTS, anterior superior temporal sulcus |
| 89 | Temporal Lobe | ITG, Inferior Temporal Gyrus | A20iv, intermediate ventral area 20 |
| 90 | Temporal Lobe | ITG, Inferior Temporal Gyrus | A20iv, intermediate ventral area 20 |

| | | | |
|---|---|---|---|
| 91 | Temporal Lobe | ITG, Inferior Temporal Gyrus | A37elv, extreme lateroventral area37 |
| 92 | Temporal Lobe | ITG, Inferior Temporal Gyrus | A37elv, extreme lateroventral area37 |
| 93 | Temporal Lobe | ITG, Inferior Temporal Gyrus | A20r, rostral area 20 |
| 94 | Temporal Lobe | ITG, Inferior Temporal Gyrus | A20r, rostral area 20 |
| 95 | Temporal Lobe | ITG, Inferior Temporal Gyrus | A20il, intermediate lateral area 20 |
| 96 | Temporal Lobe | ITG, Inferior Temporal Gyrus | A20il, intermediate lateral area 20 |
| 97 | Temporal Lobe | ITG, Inferior Temporal Gyrus | A37vl, ventrolateral area 37 |
| 98 | Temporal Lobe | ITG, Inferior Temporal Gyrus | A37vl, ventrolateral area 37 |
| 99 | Temporal Lobe | ITG, Inferior Temporal Gyrus | A20cl, caudolateral of area 20 |
| 100 | Temporal Lobe | ITG, Inferior Temporal Gyrus | A20cl, caudolateral of area 20 |
| 101 | Temporal Lobe | ITG, Inferior Temporal Gyrus | A20cv, caudoventral of area 20 |
| 102 | Temporal Lobe | ITG, Inferior Temporal Gyrus | A20cv, caudoventral of area 20 |
| 103 | Temporal Lobe | FuG, Fusiform Gyrus | A20rv, rostroventral area 20 |
| 104 | Temporal Lobe | FuG, Fusiform Gyrus | A20rv, rostroventral area 20 |

| | | | |
|---|---|---|---|
| 109 | Temporal Lobe | PhG, Parahippocampal Gyrus | A35/36r, rostral area 35/36 |
| 110 | Temporal Lobe | PhG, Parahippocampal Gyrus | A35/36r, rostral area 35/36 |
| 111 | Temporal Lobe | PhG, Parahippocampal Gyrus | A35/36c, caudal area 35/36 |
| 116 | Temporal Lobe | PhG, Parahippocampal Gyrus | A28/34, area 28/34 (EC, entorhinal cortex) |
| 118 | Temporal Lobe | PhG, Parahippocampal Gyrus | TI, area TI(temporal agranular insular cortex) |
| 131 | Parietal Lobe | SPL, Superior Parietal Lobule | A7pc, postcentral area 7 |
| 132 | Parietal Lobe | SPL, Superior Parietal Lobule | A7pc, postcentral area 7 |
| 134 | Parietal Lobe | SPL, Superior Parietal Lobule | A7ip, intraparietal area 7(hIP3) |
| 135 | Parietal Lobe | IPL, Inferior Parietal Lobule | A39c, caudal area 39(PGp) |
| 136 | Parietal Lobe | IPL, Inferior Parietal Lobule | A39c, caudal area 39(PGp) |
| 137 | Parietal Lobe | IPL, Inferior Parietal Lobule | A39rd, rostrodorsal area 39(Hip3) |
| 138 | Parietal Lobe | IPL, Inferior Parietal Lobule | A39rd, rostrodorsal area 39(Hip3) |
| 139 | Parietal Lobe | IPL, Inferior Parietal Lobule | A40rd, rostrodorsal area 40(PFt) |
| 140 | Parietal Lobe | IPL, Inferior Parietal Lobule | A40rd, rostrodorsal area 40(PFt) |

| | | | |
|---|---|---|---|
| 141 | Parietal Lobe | IPL, Inferior Parietal Lobule | A40c, caudal area 40(PFm) |
| 142 | Parietal Lobe | IPL, Inferior Parietal Lobule | A40c, caudal area 40(PFm) |
| 143 | Parietal Lobe | IPL, Inferior Parietal Lobule | A39rv, rostroventral area 39(PGa) |
| 144 | Parietal Lobe | IPL, Inferior Parietal Lobule | A39rv, rostroventral area 39(PGa) |
| 145 | Parietal Lobe | IPL, Inferior Parietal Lobule | A40rv, rostroventral area 40(PFop) |
| 146 | Parietal Lobe | IPL, Inferior Parietal Lobule | A40rv, rostroventral area 40(PFop) |
| 151 | Parietal Lobe | Pcun, Precuneus | dmPOS, dorsomedial parietooccipital sulcus(PEr) |
| 153 | Parietal Lobe | Pcun, Precuneus | A31, area 31 (Lc1) |
| 154 | Parietal Lobe | Pcun, Precuneus | A31, area 31 (Lc1) |
| 179 | Limbic Lobe | CG, Cingulate Gyrus | A32p, pregenual area 32 |
| 180 | Limbic Lobe | CG, Cingulate Gyrus | A32p, pregenual area 32 |
| 199 | Occipital Lobe | LOcC, lateral Occipital Cortex | mOccG, middle occipital gyrus |
| 200 | Occipital Lobe | LOcC, lateral Occipital Cortex | mOccG, middle occipital gyrus |

| | | | | |
|---|---|---|---|---|
| | 201 | Occipital Lobe | LOcC, lateral Occipital Cortex | V5/MT+, area V5/MT+ |
| | 202 | Occipital Lobe | LOcC, lateral Occipital Cortex | V5/MT+, area V5/MT+ |
| | 203 | Occipital Lobe | LOcC, lateral Occipital Cortex | OPC, occipital polar cortex |
| | 204 | Occipital Lobe | LOcC, lateral Occipital Cortex | OPC, occipital polar cortex |
| | 205 | Occipital Lobe | LOcC, lateral Occipital Cortex | iOccG, inferior occipital gyrus |
| | 207 | Occipital Lobe | LOcC, lateral Occipital Cortex | msOccG, medial superior occipital gyrus |
| | 208 | Occipital Lobe | LOcC, lateral Occipital Cortex | msOccG, medial superior occipital gyrus |
| | 209 | Occipital Lobe | LOcC, lateral Occipital Cortex | lsOccG, lateral superior occipital gyrus |
| | 210 | Occipital Lobe | LOcC, lateral Occipital Cortex | lsOccG, lateral superior occipital gyrus |
| $SCZ_{fMRI2}$ | 45 | Frontal Lobe | OrG, Orbital Gyrus | A11l, lateral area 11 |
| | 46 | Frontal Lobe | OrG, Orbital Gyrus | A11l, lateral area 11 |
| | 49 | Frontal Lobe | OrG, Orbital Gyrus | A13, area 13 |
| | 50 | Frontal Lobe | OrG, Orbital Gyrus | A13, area 13 |

| | | | |
|---|---|---|---|
| 60 | Frontal Lobe | PrG, Precentral Gyrus | A4t, area 4(trunk region) |
| 73 | Temporal Lobe | STG, Superior Temporal Gyrus | TE1.0 and TE1.2 |
| 114 | Temporal Lobe | PhG, Parahippocampal Gyrus | TL, area TL (lateral PPHC, posterior parahippocampal gyrus) |
| 120 | Temporal Lobe | PhG, Parahippocampal Gyrus | TH, area TH (medial PPHC) |
| 162 | Parietal Lobe | PoG, Postcentral Gyrus | A1/2/3tru, area1/2/3(trunk region) |
| 163 | Insular Lobe | INS, Insular Gyrus | G, hypergranular insula |
| 164 | Insular Lobe | INS, Insular Gyrus | G, hypergranular insula |
| 165 | Insular Lobe | INS, Insular Gyrus | vIa, ventral agranular insula |
| 169 | Insular Lobe | INS, Insular Gyrus | vId/vIg, ventral dysgranular and granular insula |
| 170 | Insular Lobe | INS, Insular Gyrus | vId/vIg, ventral dysgranular and granular insula |
| 171 | Insular Lobe | INS, Insular Gyrus | dIg, dorsal granular insula |
| 175 | Limbic Lobe | CG, Cingulate Gyrus | A23d, dorsal area 23 |
| 176 | Limbic Lobe | CG, Cingulate Gyrus | A23d, dorsal area 23 |

| | | | |
|---|---|---|---|
| 177 | Limbic Lobe | CG, Cingulate Gyrus | A24rv, rostroventral area 24 |
| 178 | Limbic Lobe | CG, Cingulate Gyrus | A24rv, rostroventral area 24 |
| 181 | Limbic Lobe | CG, Cingulate Gyrus | A23v, ventral area 23 |
| 182 | Limbic Lobe | CG, Cingulate Gyrus | A23v, ventral area 23 |
| 183 | Limbic Lobe | CG, Cingulate Gyrus | A24cd, caudodorsal area 24 |
| 184 | Limbic Lobe | CG, Cingulate Gyrus | A24cd, caudodorsal area 24 |
| 187 | Limbic Lobe | CG, Cingulate Gyrus | A32sg, subgenual area 32 |
| 188 | Limbic Lobe | CG, Cingulate Gyrus | A32sg, subgenual area 32 |
| 195 | Occipital Lobe | MVOcC, MedioVentral Occipital Cortex | rLinG, rostral lingual gyrus |
| 196 | Occipital Lobe | MVOcC, MedioVentral Occipital Cortex | rLinG, rostral lingual gyrus |
| 212 | Subcortical Nuclei | Amyg, Amygdala | mAmyg, medial amygdala |
| 214 | Subcortical Nuclei | Amyg, Amygdala | lAmyg, lateral amygdala |
| 219 | Subcortical Nuclei | BG, Basal Ganglia | vCa, ventral caudate |
| 220 | Subcortical Nuclei | BG, Basal Ganglia | vCa, ventral caudate |

| | | | | |
|---|---|---|---|---|
| | 221 | Subcortical Nuclei | BG, Basal Ganglia | GP, globus pallidus |
| | 222 | Subcortical Nuclei | BG, Basal Ganglia | GP, globus pallidus |
| | 223 | Subcortical Nuclei | BG, Basal Ganglia | NAC, nucleus accumbens |
| | 224 | Subcortical Nuclei | BG, Basal Ganglia | NAC, nucleus accumbens |
| | 225 | Subcortical Nuclei | BG, Basal Ganglia | vmPu, ventromedial putamen |
| | 226 | Subcortical Nuclei | BG, Basal Ganglia | vmPu, ventromedial putamen |
| | 227 | Subcortical Nuclei | BG, Basal Ganglia | dCa, dorsal caudate |
| | 228 | Subcortical Nuclei | BG, Basal Ganglia | dCa, dorsal caudate |
| | 229 | Subcortical Nuclei | BG, Basal Ganglia | dlPu, dorsolateral putamen |
| | 230 | Subcortical Nuclei | BG, Basal Ganglia | dlPu, dorsolateral putamen |
| | 231 | Subcortical Nuclei | Tha, Thalamus | mPFtha, medial pre-frontal thalamus |
| | 232 | Subcortical Nuclei | Tha, Thalamus | mPFtha, medial pre-frontal thalamus |
| | 233 | Subcortical Nuclei | Tha, Thalamus | mPMtha, pre-motor thalamus |
| | 234 | Subcortical Nuclei | Tha, Thalamus | mPMtha, pre-motor thalamus |
| $SCZ_{fMRI3}$ | 185 | Limbic Lobe | CG, Cingulate Gyrus | A23c, caudal area 23 |
| | 186 | Limbic Lobe | CG, Cingulate Gyrus | A23c, caudal area 23 |

| | | | |
|---|---|---|---|
| 215 | Subcortical Nuclei | Hipp, Hippocampus | rHipp, rostral hippocampus |
| 216 | Subcortical Nuclei | Hipp, Hippocampus | rHipp, rostral hippocampus |
| 217 | Subcortical Nuclei | Hipp, Hippocampus | cHipp, caudal hippocampus |
| 218 | Subcortical Nuclei | Hipp, Hippocampus | cHipp, caudal hippocampus |
| 235 | Subcortical Nuclei | Tha, Thalamus | Stha, sensory thalamus |
| 236 | Subcortical Nuclei | Tha, Thalamus | Stha, sensory thalamus |
| 237 | Subcortical Nuclei | Tha, Thalamus | rTtha, rostral temporal thalamus |
| 238 | Subcortical Nuclei | Tha, Thalamus | rTtha, rostral temporal thalamus |
| 239 | Subcortical Nuclei | Tha, Thalamus | PPtha, posterior parietal thalamus |
| 240 | Subcortical Nuclei | Tha, Thalamus | PPtha, posterior parietal thalamus |
| 241 | Subcortical Nuclei | Tha, Thalamus | Otha, occipital thalamus |
| 242 | Subcortical Nuclei | Tha, Thalamus | Otha, occipital thalamus |
| 243 | Subcortical Nuclei | Tha, Thalamus | cTtha, caudal temporal thalamus |
| 244 | Subcortical Nuclei | Tha, Thalamus | cTtha, caudal temporal thalamus |
| 245 | Subcortical Nuclei | Tha, Thalamus | lPFtha, lateral pre-frontal thalamus |
| 246 | Subcortical Nuclei | Tha, Thalamus | lPFtha, lateral pre-frontal thalamus |

| | | | | |
|---|---|---|---|---|
| $SCZ_{fMRI4}$ | 7 | Frontal Lobe | SFG, Superior Frontal Gyrus | A6dl, dorsolateral area 6 |
| | 8 | Frontal Lobe | SFG, Superior Frontal Gyrus | A6dl, dorsolateral area 6 |
| | 9 | Frontal Lobe | SFG, Superior Frontal Gyrus | A6m, medial area 6 |
| | 10 | Frontal Lobe | SFG, Superior Frontal Gyrus | A6m, medial area 6 |
| | 25 | Frontal Lobe | MFG, Middle Frontal Gyrus | A6vl, ventrolateral area 6 |
| | 39 | Frontal Lobe | IFG, Inferior Frontal Gyrus | A44v, ventral area 44 |
| | 53 | Frontal Lobe | PrG, Precentral Gyrus | A4hf, area 4(head and face region) |
| | 54 | Frontal Lobe | PrG, Precentral Gyrus | A4hf, area 4(head and face region) |
| | 55 | Frontal Lobe | PrG, Precentral Gyrus | A6cdl, caudal dorsolateral area 6 |
| | 56 | Frontal Lobe | PrG, Precentral Gyrus | A6cdl, caudal dorsolateral area 6 |
| | 57 | Frontal Lobe | PrG, Precentral Gyrus | A4ul, area 4(upper limb region) |
| | 58 | Frontal Lobe | PrG, Precentral Gyrus | A4ul, area 4(upper limb region) |
| | 59 | Frontal Lobe | PrG, Precentral Gyrus | A4t, area 4(trunk region) |
| | 61 | Frontal Lobe | PrG, Precentral Gyrus | A4tl, area 4(tongue and larynx region) |
| | 62 | Frontal Lobe | PrG, Precentral Gyrus | A4tl, area 4(tongue and larynx region) |
| | 63 | Frontal Lobe | PrG, Precentral Gyrus | A6cvl, caudal ventrolateral area 6 |

| | | | |
|---|---|---|---|
| 64 | Frontal Lobe | PrG, Precentral Gyrus | A6cvl, caudal ventrolateral area 6 |
| 65 | Frontal Lobe | PCL, Paracentral Lobule | A1/2/3ll, area1/2/3 (lower limb region) |
| 66 | Frontal Lobe | PCL, Paracentral Lobule | A1/2/3ll, area1/2/3 (lower limb region) |
| 67 | Frontal Lobe | PCL, Paracentral Lobule | A4ll, area 4, (lower limb region) |
| 68 | Frontal Lobe | PCL, Paracentral Lobule | A4ll, area 4, (lower limb region) |
| 72 | Temporal Lobe | STG, Superior Temporal Gyrus | A41/42, area 41/42 |
| 74 | Temporal Lobe | STG, Superior Temporal Gyrus | TE1.0 and TE1.2 |
| 76 | Temporal Lobe | STG, Superior Temporal Gyrus | A22c, caudal area 22 |
| 77 | Temporal Lobe | STG, Superior Temporal Gyrus | A38l, lateral area 38 |
| 79 | Temporal Lobe | STG, Superior Temporal Gyrus | A22r, rostral area 22 |
| 105 | Temporal Lobe | FuG, Fusiform Gyrus | A37mv, medioventral area37 |
| 106 | Temporal Lobe | FuG, Fusiform Gyrus | A37mv, medioventral area37 |
| 107 | Temporal Lobe | FuG, Fusiform Gyrus | A37lv, lateroventral area37 |
| 108 | Temporal Lobe | FuG, Fusiform Gyrus | A37lv, lateroventral area37 |

| | | | |
|---|---|---|---|
| 112 | Temporal Lobe | PhG, Parahippocampal Gyrus | A35/36c, caudal area 35/36 |
| 113 | Temporal Lobe | PhG, Parahippocampal Gyrus | TL, area TL (lateral PPHC, posterior parahippocampal gyrus) |
| 115 | Temporal Lobe | PhG, Parahippocampal Gyrus | A28/34, area 28/34 (EC, entorhinal cortex) |
| 117 | Temporal Lobe | PhG, Parahippocampal Gyrus | TI, area TI(temporal agranular insular cortex) |
| 119 | Temporal Lobe | PhG, Parahippocampal Gyrus | TH, area TH (medial PPHC) |
| 121 | Temporal Lobe | pSTS, posterior Superior Temporal Sulcus | rpSTS, rostroposterior superior temporal sulcus |
| 122 | Temporal Lobe | pSTS, posterior Superior Temporal Sulcus | rpSTS, rostroposterior superior temporal sulcus |
| 123 | Temporal Lobe | pSTS, posterior Superior Temporal Sulcus | cpSTS, caudoposterior superior temporal sulcus |
| 124 | Temporal Lobe | pSTS, posterior Superior Temporal Sulcus | cpSTS, caudoposterior superior temporal sulcus |
| 125 | Parietal Lobe | SPL, Superior Parietal Lobule | A7r, rostral area 7 |
| 126 | Parietal Lobe | SPL, Superior Parietal Lobule | A7r, rostral area 7 |

| | | | |
|---|---|---|---|
| 127 | Parietal Lobe | SPL, Superior Parietal Lobule | A7c, caudal area 7 |
| 128 | Parietal Lobe | SPL, Superior Parietal Lobule | A7c, caudal area 7 |
| 129 | Parietal Lobe | SPL, Superior Parietal Lobule | A5l, lateral area 5 |
| 130 | Parietal Lobe | SPL, Superior Parietal Lobule | A5l, lateral area 5 |
| 133 | Parietal Lobe | SPL, Superior Parietal Lobule | A7ip, intraparietal area 7(hIP3) |
| 147 | Parietal Lobe | Pcun, Precuneus | A7m, medial area 7(PEp) |
| 148 | Parietal Lobe | Pcun, Precuneus | A7m, medial area 7(PEp) |
| 149 | Parietal Lobe | Pcun, Precuneus | A5m, medial area 5(PEm) |
| 150 | Parietal Lobe | Pcun, Precuneus | A5m, medial area 5(PEm) |
| 152 | Parietal Lobe | Pcun, Precuneus | dmPOS, dorsomedial parietooccipital sulcus(PEr) |
| 155 | Parietal Lobe | PoG, Postcentral Gyrus | A1/2/3ulhf, area 1/2/3(upper limb, head and face region) |
| 156 | Parietal Lobe | PoG, Postcentral Gyrus | A1/2/3ulhf, area 1/2/3(upper limb, head and face region) |
| 157 | Parietal Lobe | PoG, Postcentral Gyrus | A1/2/3tonIa, area 1/2/3(tongue and larynx region) |
| 158 | Parietal Lobe | PoG, Postcentral Gyrus | A1/2/3tonIa, area 1/2/3(tongue and larynx region) |

| | | | |
|---|---|---|---|
| 159 | Parietal Lobe | PoG, Postcentral Gyrus | A2, area 2 |
| 160 | Parietal Lobe | PoG, Postcentral Gyrus | A2, area 2 |
| 161 | Parietal Lobe | PoG, Postcentral Gyrus | A1/2/3tru, area1/2/3(trunk region) |
| 166 | Insular Lobe | INS, Insular Gyrus | vIa, ventral agranular insula |
| 167 | Insular Lobe | INS, Insular Gyrus | dIa, dorsal agranular insula |
| 168 | Insular Lobe | INS, Insular Gyrus | dIa, dorsal agranular insula |
| 172 | Insular Lobe | INS, Insular Gyrus | dIg, dorsal granular insula |
| 173 | Insular Lobe | INS, Insular Gyrus | dId, dorsal dysgranular insula |
| 174 | Insular Lobe | INS, Insular Gyrus | dId, dorsal dysgranular insula |
| 189 | Occipital Lobe | MVOcC, MedioVentral Occipital Cortex | cLinG, caudal lingual gyrus |
| 190 | Occipital Lobe | MVOcC, MedioVentral Occipital Cortex | cLinG, caudal lingual gyrus |
| 191 | Occipital Lobe | MVOcC, MedioVentral Occipital Cortex | rCunG, rostral cuneus gyrus |
| 192 | Occipital Lobe | MVOcC, MedioVentral | rCunG, rostral cuneus gyrus |

| | | | |
|---|---|---|---|
| | | Occipital Cortex | |
| 193 | Occipital Lobe | MVOcC, MedioVentral Occipital Cortex | cCunG, caudal cuneus gyrus |
| 194 | Occipital Lobe | MVOcC, MedioVentral Occipital Cortex | cCunG, caudal cuneus gyrus |
| 197 | Occipital Lobe | MVOcC, MedioVentral Occipital Cortex | vmPOS,ventromedial parietooccipital sulcus |
| 198 | Occipital Lobe | MVOcC, MedioVentral Occipital Cortex | vmPOS,ventromedial parietooccipital sulcus |
| 206 | Occipital Lobe | LOcC, lateral Occipital Cortex | iOccG, inferior occipital gyrus |
| 211 | Subcortical Nuclei | Amyg, Amygdala | mAmyg, medial amygdala |
| 213 | Subcortical Nuclei | Amyg, Amygdala | lAmyg, lateral amygdala |
